# A perinuclear microtubule-organizing center controls nuclear positioning and basement membrane secretion

**DOI:** 10.1101/2019.12.24.888065

**Authors:** Yiming Zheng, Rebecca A. Buchwalter, Chunfeng Zheng, Elise M. Wight, Jieyan V. Chen, Timothy L. Megraw

## Abstract

Non-centrosomal microtubule-organizing centers (ncMTOCs) have a variety of roles presumed to serve the diverse functions of the range of cell types in which they are found. ncMTOCs are diverse in their composition, subcellular localization, and function. Here we report a novel perinuclear MTOC in *Drosophila* fat body cells that is anchored by Msp300/Nesprin at the cytoplasmic surface of the nucleus. Msp300 recruits the MT minus-end protein Patronin/CAMSAP, which functions redundantly with Ninein to further recruit the MT polymerase Msps/XMAP215 to assemble non-centrosomal MTs and does so independently of the widespread MT nucleation factor *γ*-tubulin. Functionally, the fat body ncMTOC and the radial MT arrays it organizes is essential for nuclear positioning and for secretion of basement membrane components via retrograde dynein-dependent endosomal trafficking that restricts plasma membrane growth. Together, this study identifies a perinuclear ncMTOC with unique architecture and MT regulation properties that serves vital functions.

**Highlights:**

- A novel perinuclear MTOC in differentiated fat body cells
- The predominant nucleator, *γ*-tubulin, is not required at the fat body ncMTOC
- Msp300/Nesprin organizes the ncMTOC at the nuclear surface by recruiting Patronin/CAMSAP and the spectraplakin Shot
- Patronin cooperates with Ninein to control MT assembly at the fat body ncMTOC by recruiting Msps
- Msps, a MT polymerase, is essential for radial MT elongation from the fat body ncMTOC
- Patronin and Msps associate
- The ncMTOC and radial MTs, but not actin, control nuclear positioning in the fat body
- The fat body MTOC controls retrograde endocytic trafficking to regulate plasma membrane growth and secretion of basement membrane proteins

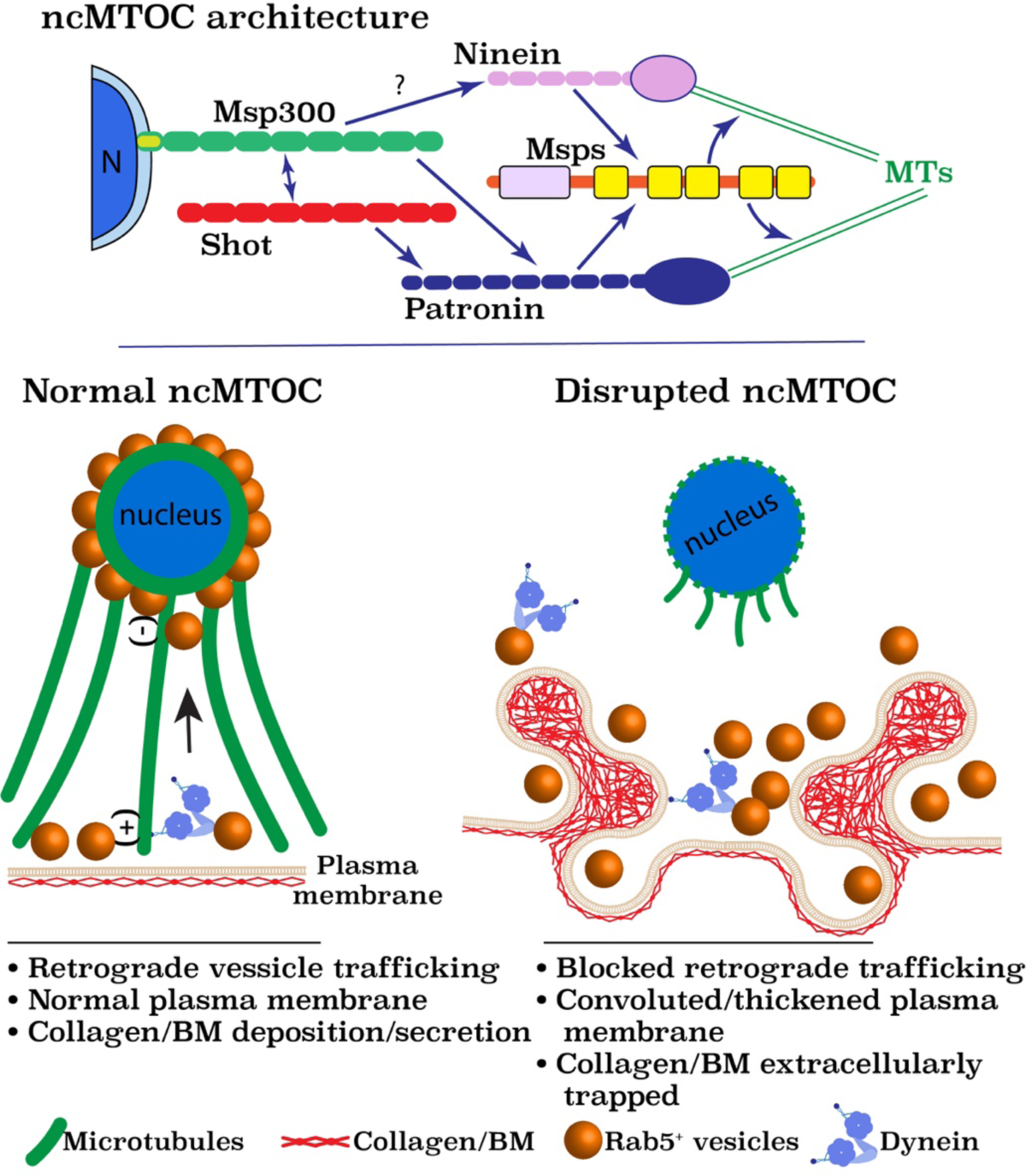

## Introduction

Microtubule (MT) organization supports critical cellular functions, including assembly of the mitotic spindle apparatus during cell division, establishment of cell polarity and cell shape, and by serving as a conduit for intracellular trafficking. The best-known microtubule-organizing center (MTOC) in animal cells is the centrosome, which organizes MT arrays from a supramolecular matrix called the pericentriolar material (PCM). However, various cell types across species, after exit from the cell cycle and upon differentiation, lack a functional centrosome. In these cases, non-centrosomal MTOCs (ncMTOCs) function as alternative sites to accommodate the organization of MT networks specialized for differentiated cell types (Muroyama and Lechler, 2017; Sanchez and Feldman, 2017; Martin and Akhmanova, 2018; Tillery et al., 2018). Whereas knowledge of the centrosome is extensive, the roles ncMTOCs play in cells, their molecular compositions, and how they are anchored to specific subcellular sites remains largely unknown.

Generation of an ncMTOC requires the nucleation, stabilization, and anchoring of MT minus-ends, generally achieved by MT minus-end-associated proteins. Compared to the relatively large number of MT plus-end proteins, few MT minus-end proteins have been identified. Gamma-tubulin (*γ*-tubulin), a conserved and essential MT minus-end protein that nucleates centrosomal MTs (Kollman et al., 2011; Oakley et al., 2015; Farache et al., 2018), also nucleates and anchors non-centrosomal MTs in many differentiated cell types (Chen et al., 2017; Muroyama and Lechler, 2017; Sanchez and Feldman, 2017; Tillery et al., 2018). The XMAP215/Msps/Stu2/ Dis1/Alp14/ZYG-9/ch-TOG/MOR1 is an ancient family of MT polymerases that was recently identified as a new *in vivo* MT nucleator at centrosomes and spindle pole bodies through its association with *γ*-tubulin (Flor-Parra et al., 2018; Gunzelmann et al., 2018; Thawani et al., 2018); a role for the MT polymerase at nucleating MTs at ncMTOCs has not been reported. Additionally, the recently identified CAMSAP/Patronin family of MT minus-end proteins protects MT minus-ends from Kinesin-13 family depolymerases (Goodwin and Vale, 2010; Hendershott and Vale, 2014; Atherton et al., 2017). CAMSAP/Patronin has emerged as a critical player at ncMTOCs via unclear mechanisms. Ninein is another MT minus-end anchoring protein, but little is known of its mechanisms of action. To understand the diversity of ncMTOCs and how they serve the unique needs of the diverse cell types where they are organized, it is essential to determine how ncMTOCs are assembled and what the key effectors of MT assembly are.

Here we report the discovery of an ncMTOC that is assembled on the surface of nuclei in *Drosophila* larval fat body cells, a differentiated cell type that has critical secretory functions and serves the metabolic needs of the organism. Assembly of this perinuclear ncMTOC requires the Msp300/Nesprin (nuclear envelope spectrin repeat protein) as a primary organizer/anchor. Msp300 recruits Patronin, which functions redundantly with Ninein for MT assembly at the ncMTOC. Patronin and Ninein cooperate to recruit Msps, which is essential for the elongation of MTs from the fat body ncMTOC. Recruitment of Msps and MT assembly at the ncMTOC are independent of *γ*-tubulin. Functionally, this ncMTOC is necessary for: 1) the regulation of nuclear positioning, 2) controlling plasma membrane growth by facilitating dynein/Rab5-mediated retrograde endosomal trafficking of excess plasma membrane, and 3) the secretion of collagen IV and other basement membrane (BM) proteins. Together, this study identifies a unique perinuclear ncMTOC with novel MT assembly mechanisms that controls physiological roles of fat body cells by supporting nuclear positioning and vital secretory functions of fat body cells.

## Results

### A perinuclear MTOC is assembled in fat body cells

Postmitotic polyploid cell types in *Drosophila*, such as salivary gland, midgut, or malpighian tubule cells, lack centrosomes (Schoenfelder et al., 2014). We found that centrosomes are also lost in polyploid fat body cells (Figure 1A), which are functionally similar to vertebrate liver and adipose cells including high metabolic and secretory activities (Hoshizaki et al., 1994; Arrese and Soulages, 2010; Frawley and Orr-Weaver, 2015; Droujinine and Perrimon, 2016).

**Figure 1.**
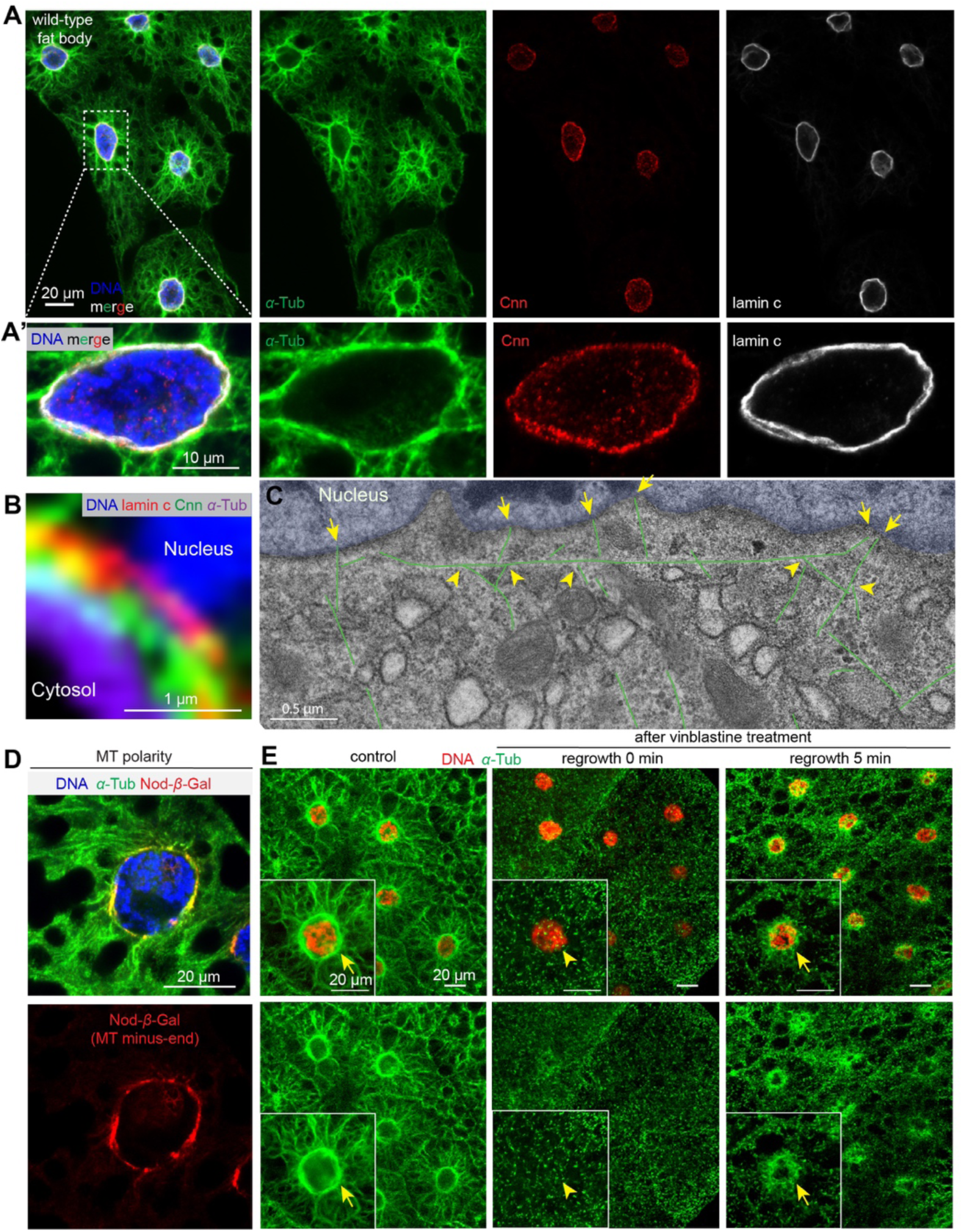
Fat body cells assemble a perinuclear ncMTOC. **A, A’**) Immunofluorescent (IF) staining of fixed fat body tissues from third instar larvae labelled to reveal MTs (*α*-tubulin), the centrosomal protein Cnn, nuclear lamin C and DNA (DAPI). (**A’**) shows magnified view of a fat body cell. (**B**) A close-up view of the nuclear surface showing MTs, Cnn, lamin and nuclear DNA. (**C**) TEM image of fat body MTs; a stitched composite of three images. MTs are colored green and the nucleus is colored in blue. Arrows: MTs anchored at nuclear surface. Arrowheads: MTs that branch from the circumferential MTs. (**D**) Localization of the Nod-*β*-gal fusion protein at the nuclear surface marks the MT minus-ends. (**E**) MT regrowth in fat body cells 5 min after recovery from vinblastine treatment shows MT assembly enriched at the perinuclear ncMTOC. Insets show magnification of a fat body cell; arrows point to sites of MT regrowth. Scale bars: 20 µm (**A**, **D, E**), 10 µm (**A**’), 1 µm (**B**), 0.5 µm **(C**).

Fat body cells have a prominent perinuclear organization of MTs (Figure 1). In each fat body cell MTs are highly enriched circumferentially at the nuclear surface and also radiate outward toward the plasma membrane (Figure 1A, A’). These observations indicate that the nuclear surface houses an ncMTOC. Consistent with this, the centrosomal protein Centrosomin (Cnn) and MTs are positioned on the cytoplasmic face of the nuclear envelope relative to intranuclear lamin C (Figure 1B). Electron microscopy imaging confirmed the presence of MTs oriented circumferentially and perpendicularly (radial) to the nuclear surface (Figure 1C). Moreover, two general populations of radial MTs are evident; those anchored at the nuclear surface, presumably by their minus ends (arrows) and another group that appear to branch from the circumferential MTs (arrowheads). Consistently, labelling of the MT minus ends with a Nod-*β*-gal fusion protein (Clark et al., 1997) shows that MTs are anchored and enriched at the MTOC on the nuclear surface (Figure 1D). Moreover, MT regrowth experiments show that the nuclear surface is the primary site for MT assembly in fat body cells (Figure 1E).

Cold treatment, which typically causes MT polymers to disassemble, did not overtly impact the MT array organized at the fat body nuclear surface (Figure 2A), indicating that these MTs are highly stable. Consistent with this, fat body MTs are acetylated and polyglutamylated (Figure 2B, C), two tubulin post-translational modifications that are associated with stabilized MTs (Song and Brady, 2015; Wloga et al., 2017). Together, these data demonstrate a novel MTOC at the nuclear surface of fat body cells that is enriched in stable MTs.

**Figure 2.**
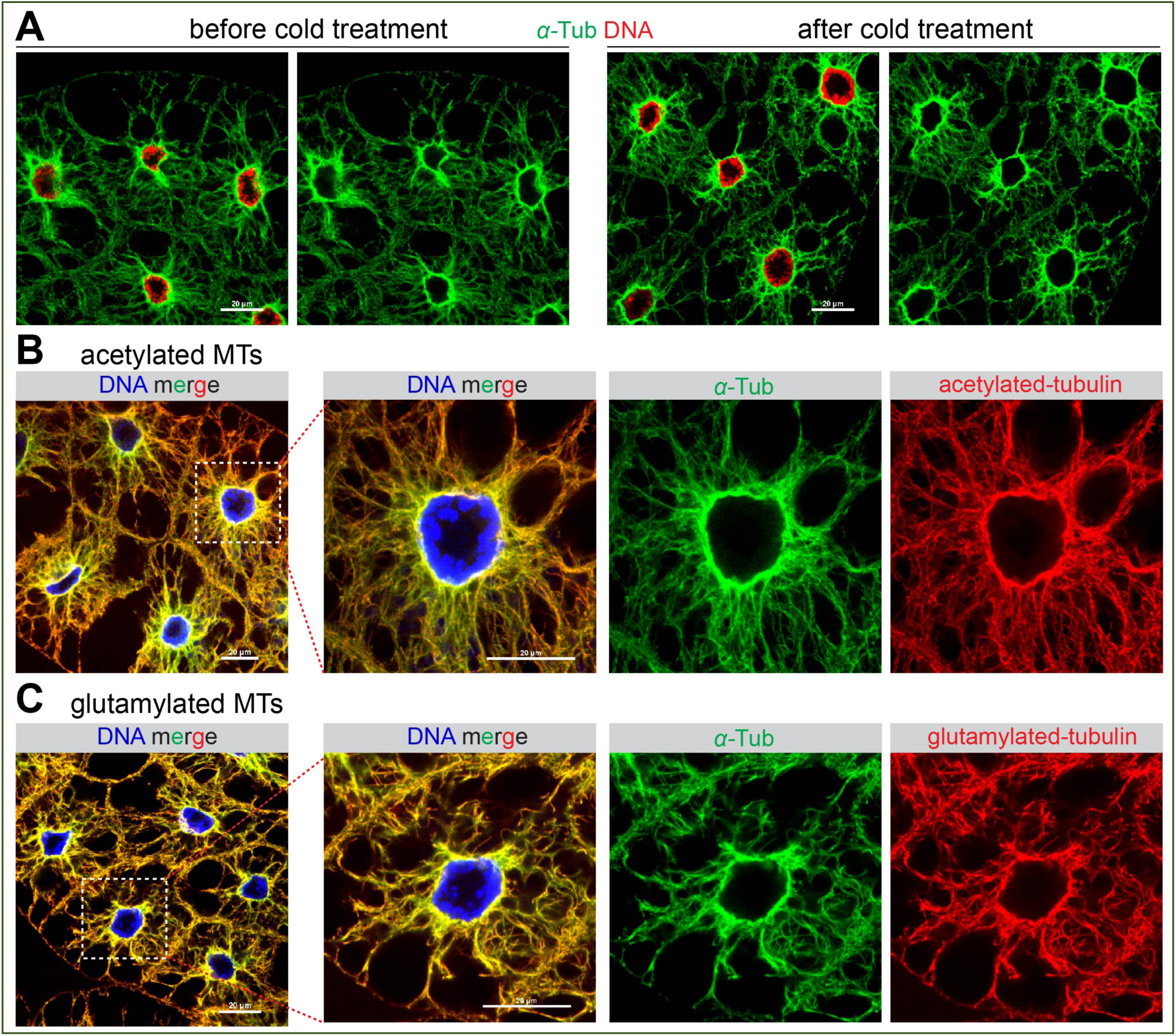
Fat body MTs are stabilized. (**A**) Images of IF stained fat body cells with or without cold treatment for 1 hr, which did not cause significant reduction of the MT array. (**B**, **C**) IF staining shows that the fat body contains MTs that are acetylated (**B**) and polyglutamylated (**C**), two post-translational modifications attributed to MT stabilization. Scale bars: 20 µm.

### Fat body microtubules, but not actin, are essential for nuclear positioning

To examine a role for the MT array in fat body cells we disrupted MTs directly using two approaches, both involving the generation of genetically mosaic “flipout” clones. In the first approach, we knocked down expression of the genes encoding the MT subunits, *α*-tubulin and *β*-tubulin, by RNAi. The flipout clones where RNAi knockdown occured were marked with GFP expression adjacent to the control cells, enabling comparison of knockdown cell phenotypes side-by-side with control cells (Figure 3A, B). Knockdown of either tubulin subunit significantly disrupted MTs and also the expression/stability of the other subunit; i.e. knockdown of *α*-tubulin resulted in reduced *β*-tubulin expression and vice versa (Figure 3A’, B’).

**Figure 3.**
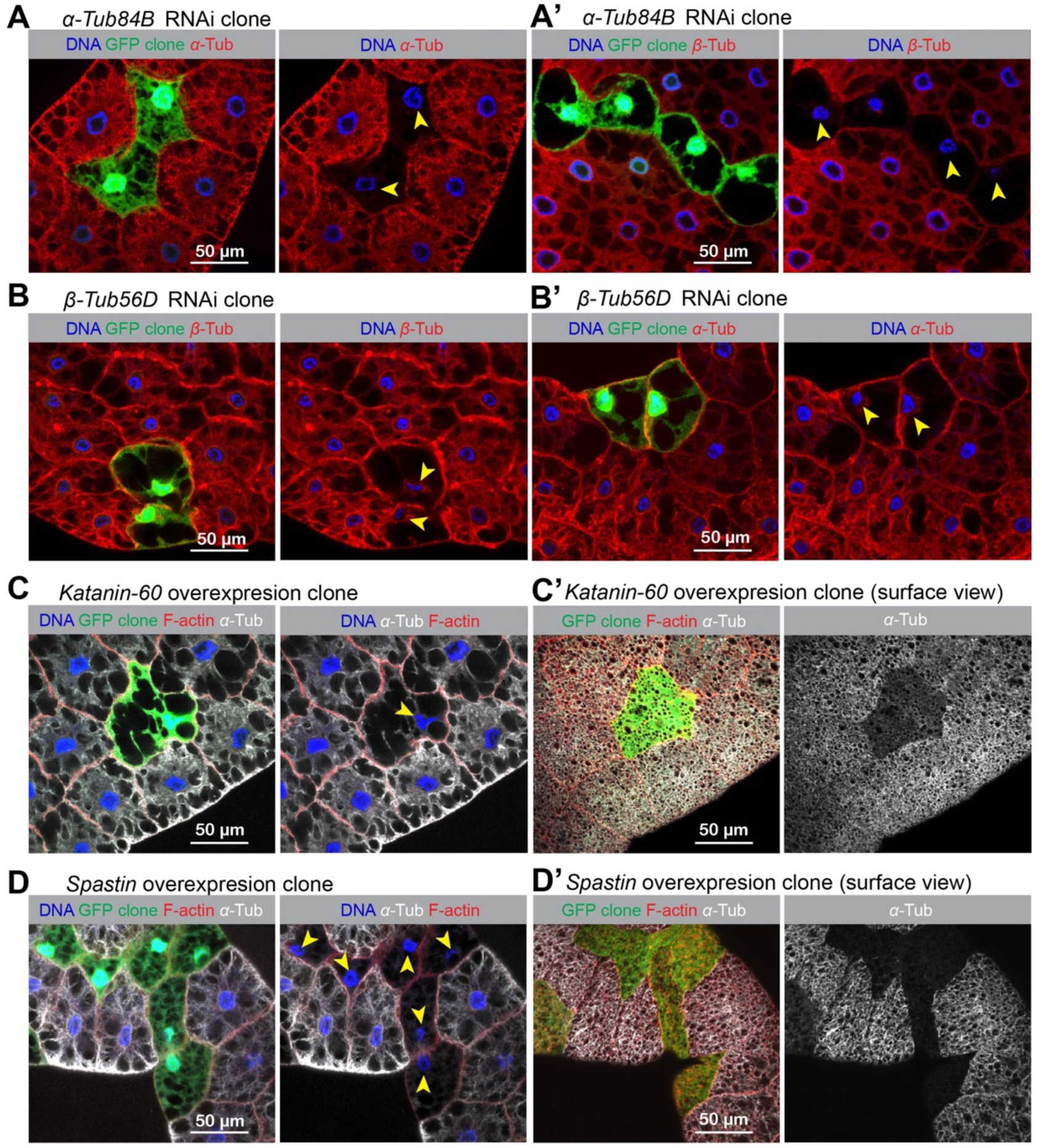
Microtubule disruption after tubulin knockdown or overexpression of MT severing enzyme. Expression of the *α*- and *β*-tubulin subunits were blocked by siRNA-mediated knockdown in somatic clones marked with GFP expression and generated using FLP-FRT mediated recombination. Arrowheads indicate nuclei that have lost centricity in effected clones. (**A**, **A**’) knockdown of *α-Tub84B* significantly reduces expression of *α*-tubulin (**A**) and also btubulin (**A**’). (**B**, **B**’) Likewise, knockdown of *β-Tub56D* blocks expression/stability of both tubulin subunits. (**C**, **C**’, **D**, **D**’) Overexpression of MT severing enzymes Spastin (**C**, **C**’) or Katanin-60 (**D**, **D**’) in GFP-marked clones also results in MT disruption. Scale bars: 50 μm.

In the second approach, we overexpressed the MT-severing enzymes Spastin or Katanin-60 to disrupt MTs. In GFP-positive clones that overexpressed each severing enzyme, fat body MTs were disrupted (Figure 3C, C’, D, D’). In each fat body cell, the nucleus resides in the geometric center (centroid) (Figure 4A). When fat body MTs were disrupted by either approach, nuclear positioning was significantly affected. In tubulin knockdown cells or Spastin or Katanin-60 overexpressing cells, the normally centroid nucleus was mispositioned (Figure 4A). In contrast to MT disruption, blocking actin nucleation by *Arp2* knockdown or *Rho1* knockdown reduced F-actin signal but did not disrupt nuclear positioning (Figure 4B). These data show that MTs, but not actin, are necessary for the proper positioning of nuclei in fat body cells, and that nuclear positioning can be used as a methodological readout for proper MT assembly or ncMTOC function in the fat body.

**Figure 4.**
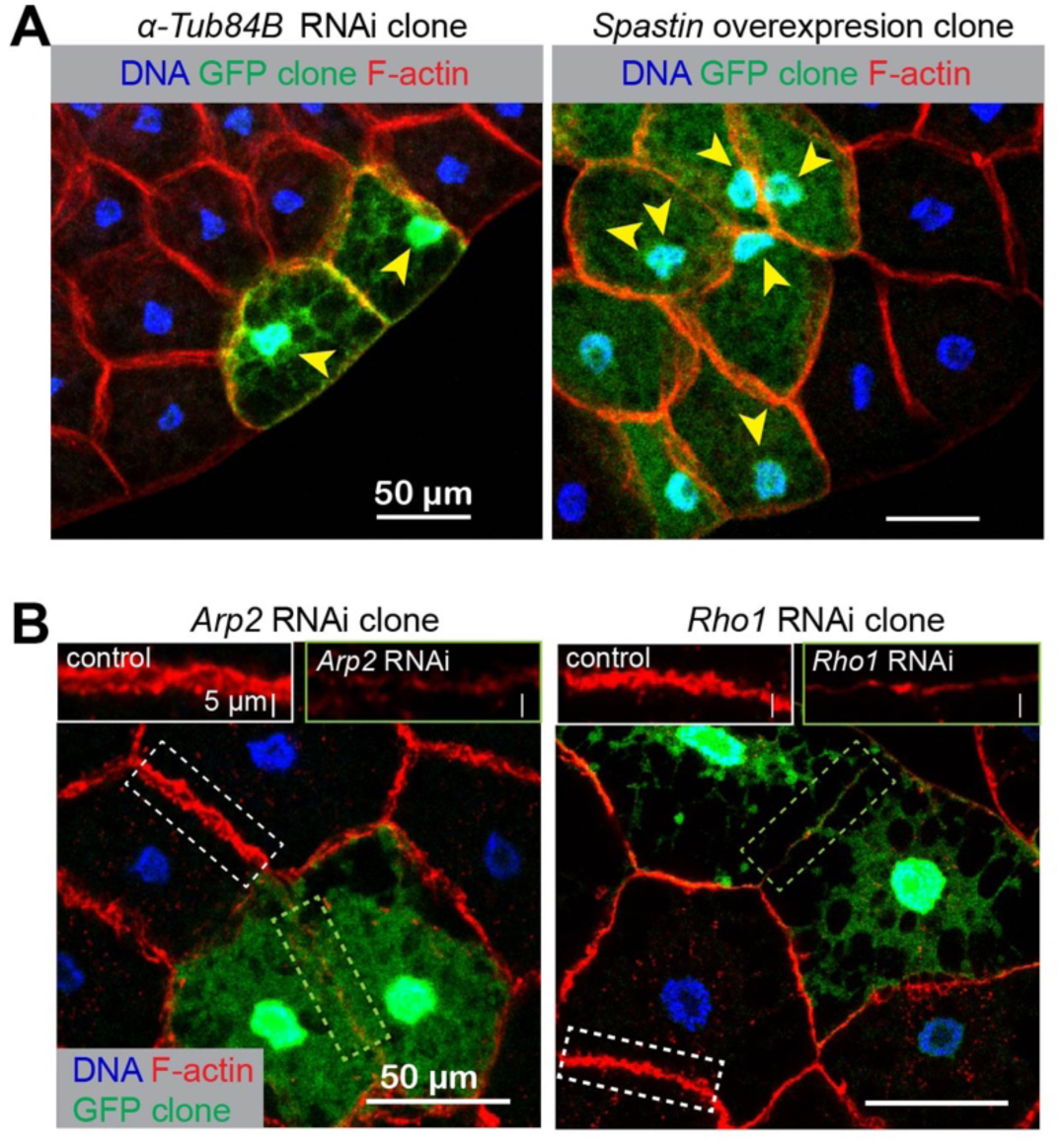
Microtubule, but not actin disruption impairs nuclear positioning. (**A**) All of these MT disruptions result in a loss of nuclear centricity in fat body cells, which are also indicated with arrowheads in Figure 3A-D’. Quantification of nuclear positioning is shown in Figure 8B. (**B**) Disruption of actin assembly by knockdown of *Arp2*, an actin nucleator, or *Rho1*, a GTPase critical for actin assembly, did not impair nuclear positioning. Insets show the reduced assembly of cortical F-actin in *Arp2* or *Rho1* RNAi cell compared to the control cell. Quantification of nuclear positioning is shown in Figure 8B. Scale bars: 50 μm.

### Centrosomal proteins are localized at the fat body ncMTOC but are not required for MT assembly

We next determined the requirement of centrosomal proteins and MT regulatory proteins for fat body MTOC functions using nuclear positioning as an assay. We surveyed mutants or RNAi-mediated knockdowns for their requirement to position the nucleus at the centroid relative to cortical F-actin or myr-RFP (myristoylation domain of Src fused to mRFP), both of which mark the cell boundary. We also surveyed the localization of centrosomal proteins and MT regulators at the fat body perinuclear MTOC using available antibodies and/or expression of tagged transgenic proteins. These data are summarized in Table 1. Validation of antibody and transgenic expression reagents for several key centrosomal components are shown in Figure 5.

**Figure 5.**
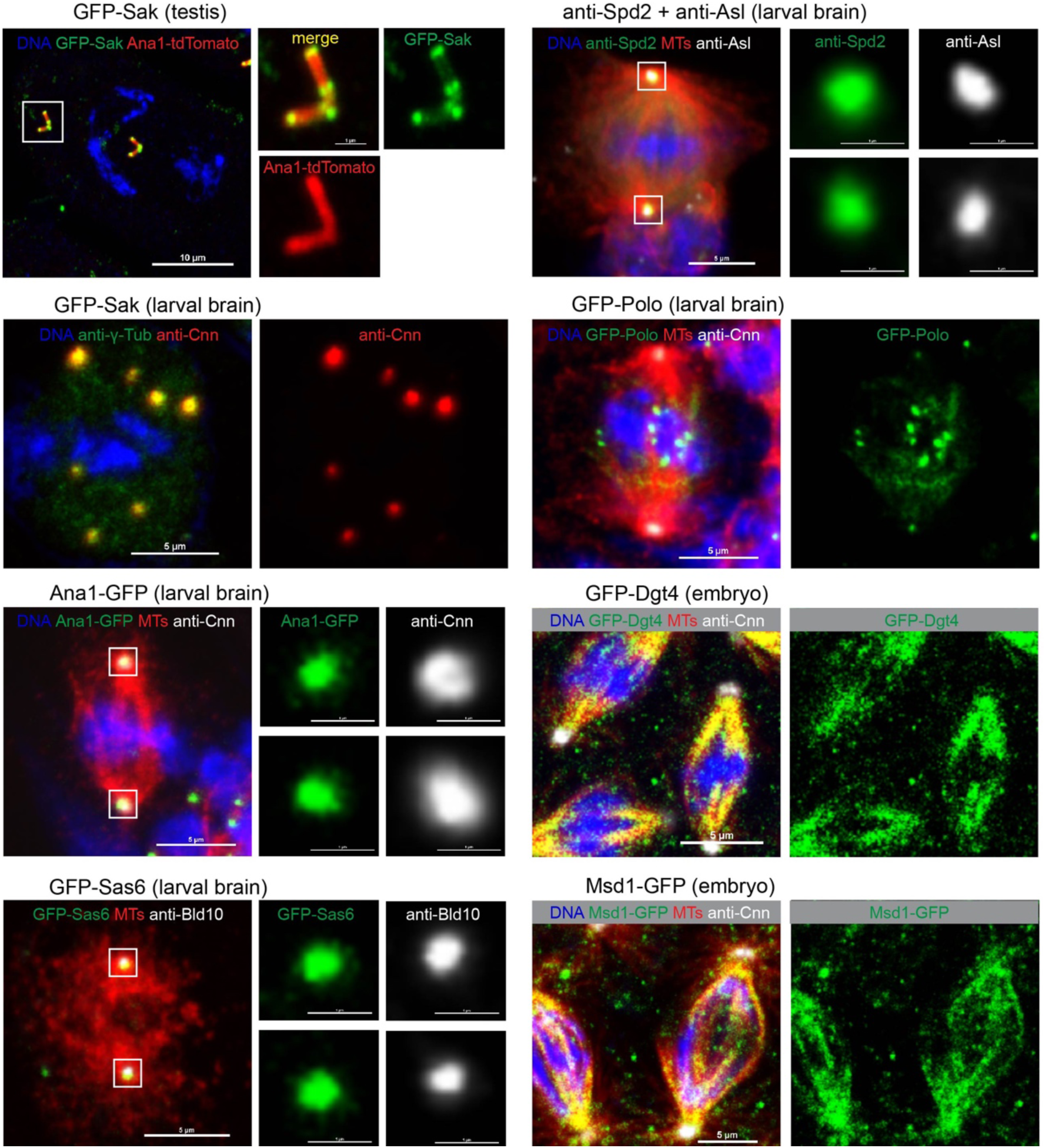
Verification of key centrosomal markers. All these centrosomal transgenes or antibodies have been published previously, except for the GFP-Dgt4 transgene, and were verified here by examining expression and localization in embryos or larval brains.

**Table 1.**
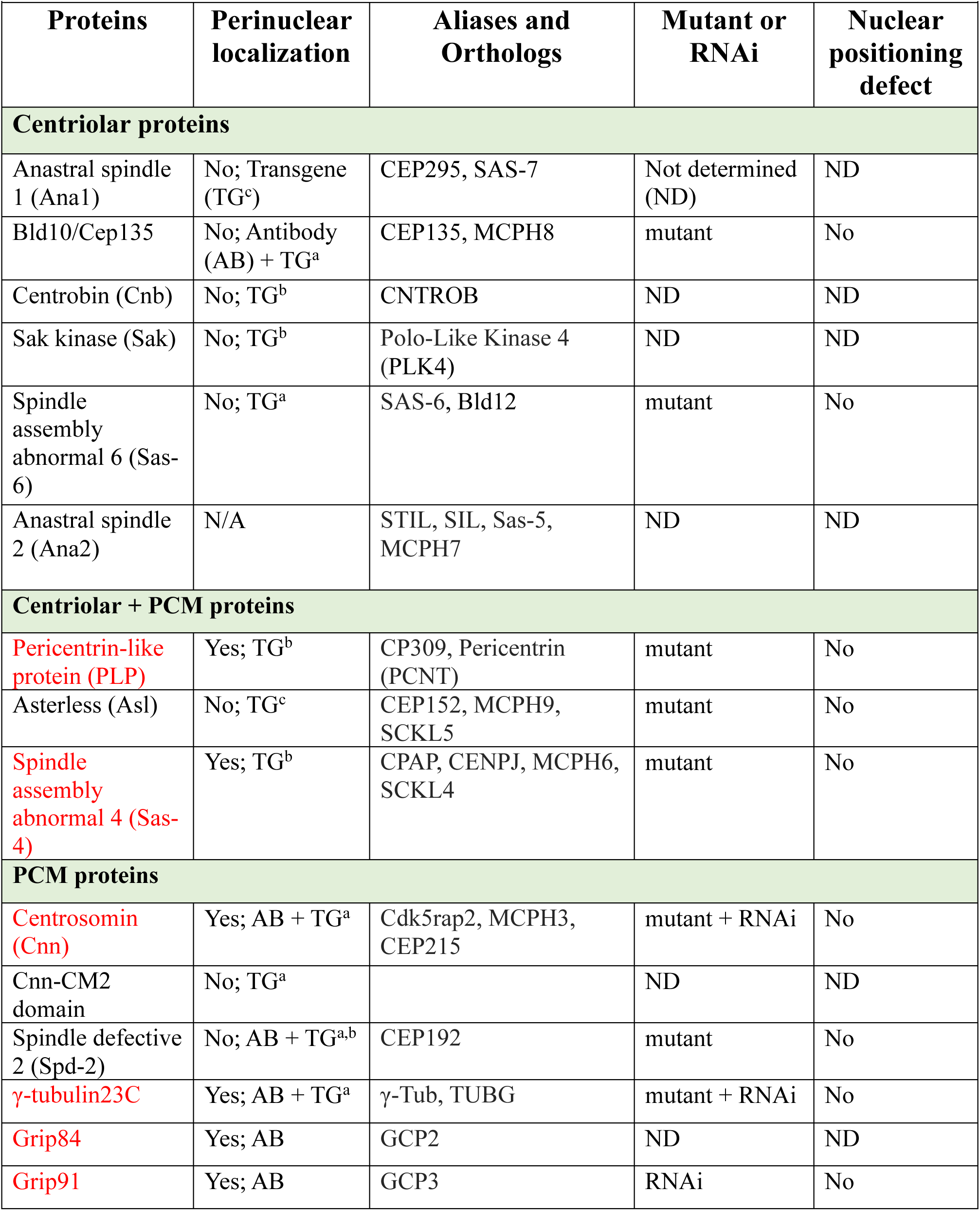
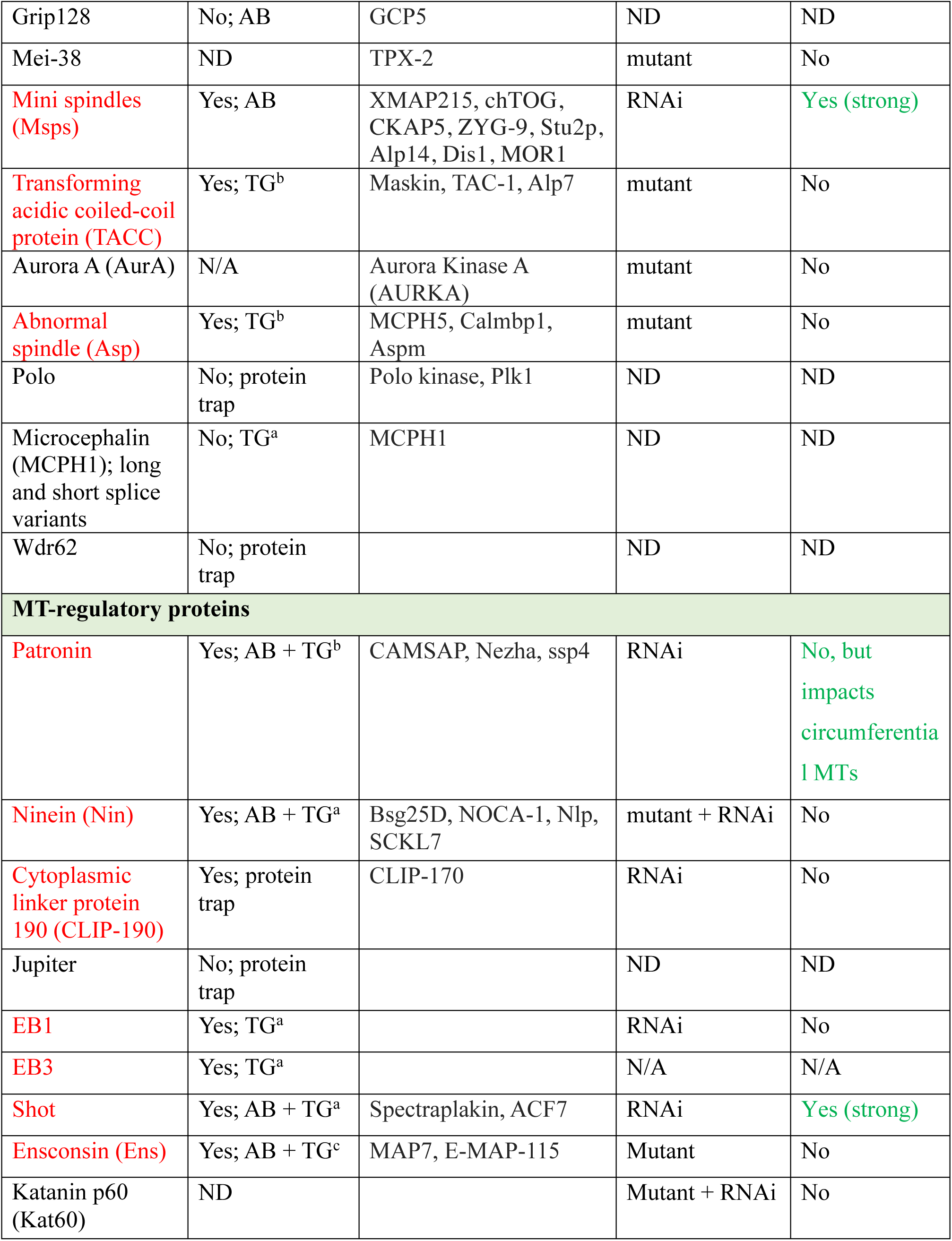
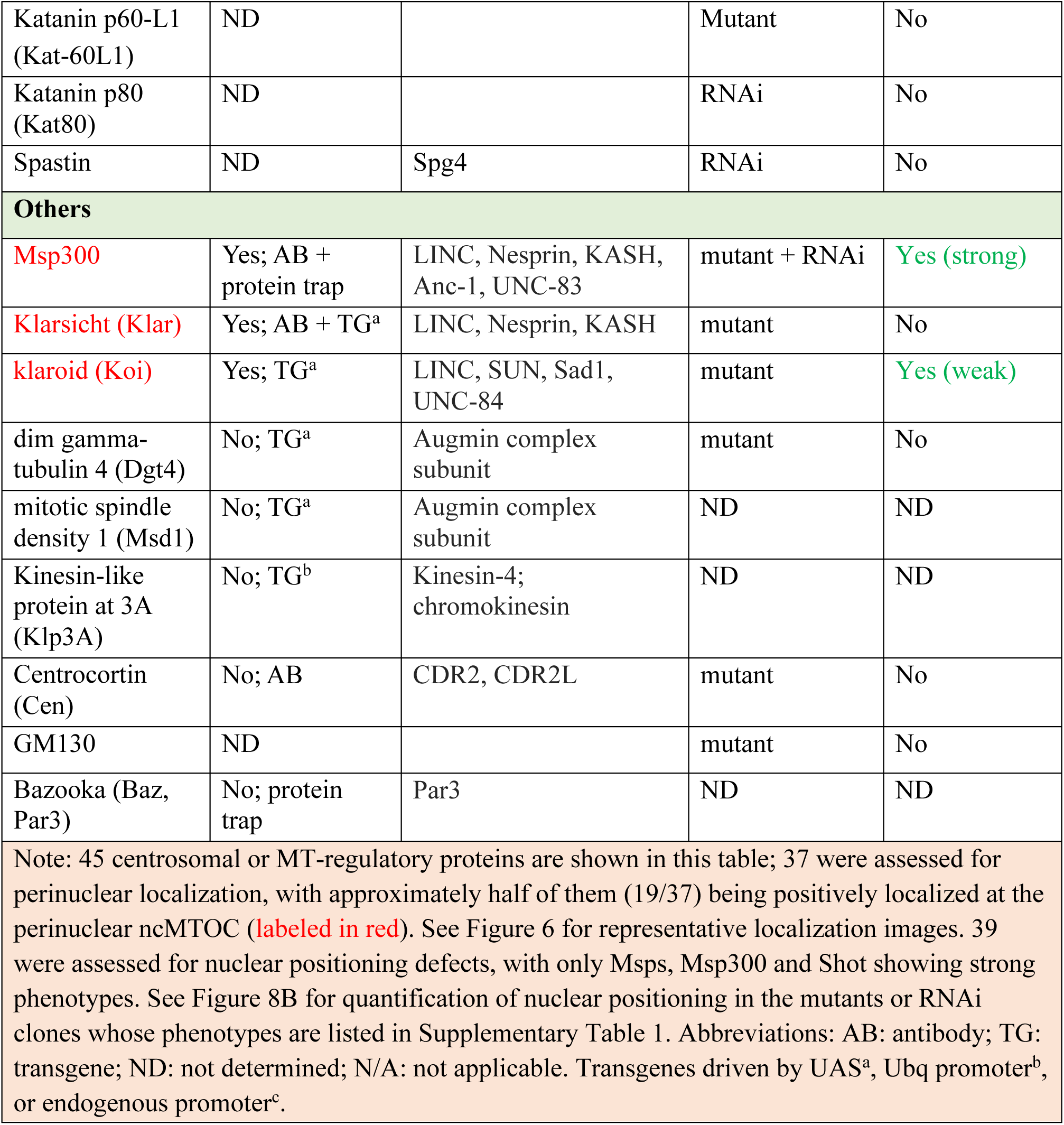
Survey of fat body ncMTOC components and key assembly regulators.

This survey showed that centrosome PCM proteins were localized at the fat body MTOC, but core centriolar proteins were absent (Figure 6, Table 1). The major core PCM components were present, with the exception of Spd-2 (Figure 6, Table 1). Since some of the transgenic proteins were expressed ectopically, it is possible that some of the positive localizations determined in this manner do not reflect endogenous protein localization. Collectively, the fat body MTOC contains some proteins in common with the centrosome PCM, but also some distinct components including Patronin (see below).

**Figure 6.**
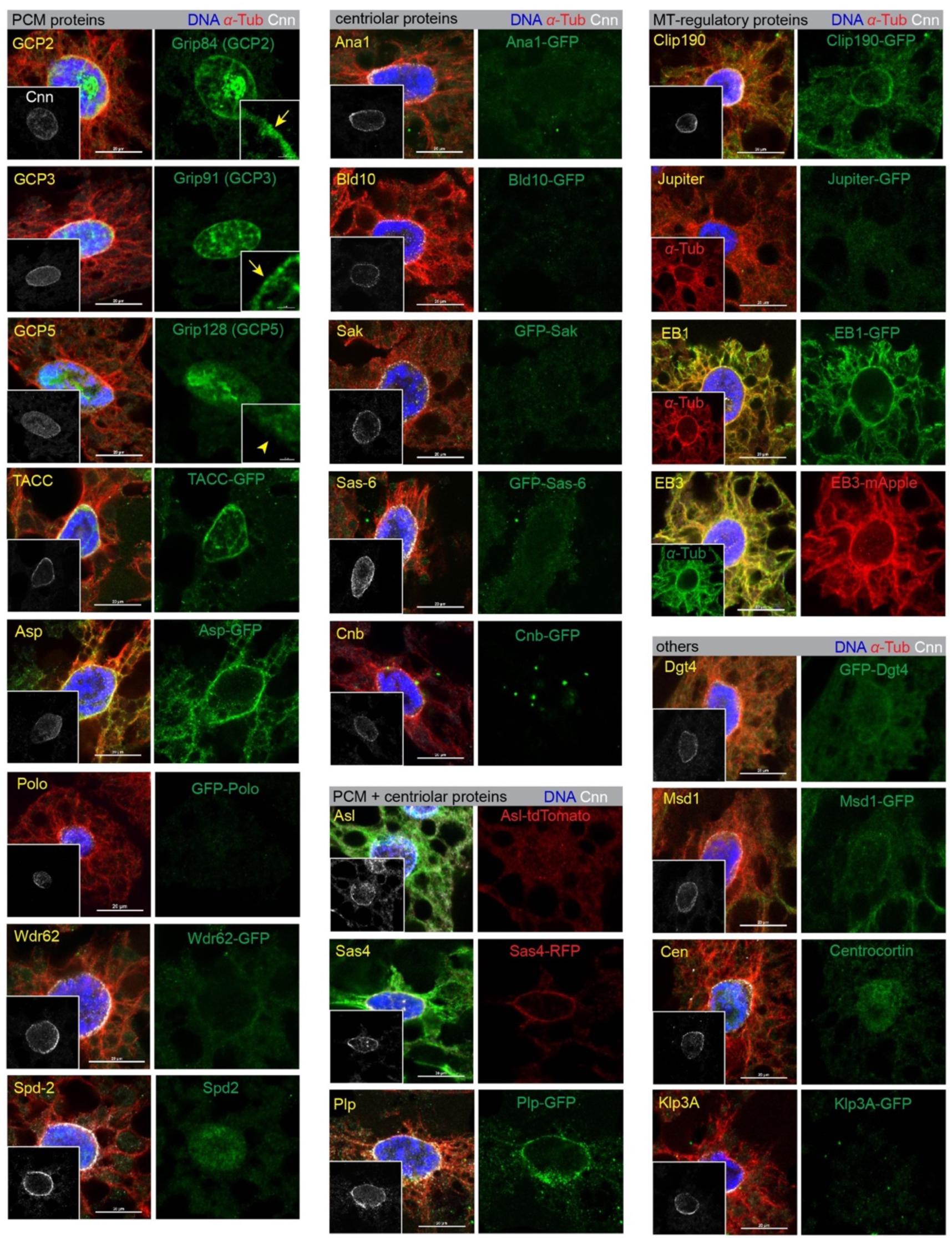
Survey of localization of centrosomal proteins and MT regulators at the fat body ncMTOC. Images of fat body cells stained for Cnn (insets) and counterstained in green or red for the indicated proteins. See Table 1 for the full list. Scale bars: 20 μm.

No single centrosomal protein gene mutant that we tested disrupted fat body nuclear centricity or ncMTOC assembly (Table 1). Among those tested were mutations in *cnn*, *nin* (*Bsg25D*), *plp*, *asp* (Figure 7A), *spd-2*, *sas-4*, *asl*, *sas-6, bld10* (*Cep135*), *centrocortin* (not shown), *tacc,* and *aurA* (see below). Reasoning that there might be functional redundancy in MT assembly, as is the case with Cnn and Spd-2 at centrosomes (Conduit et al., 2014), we tested pairwise combinations of mutants or RNAi knockdowns, but still found no combination of centrosomal proteins that was required for the fat body ncMTOC to assemble MTs or affect nuclear positioning (Figure 7B). These data indicate that although the fat body ncMTOC shares a majority of the core centrosome PCM components, it may not require them for MTOC function.

**Figure 7.**
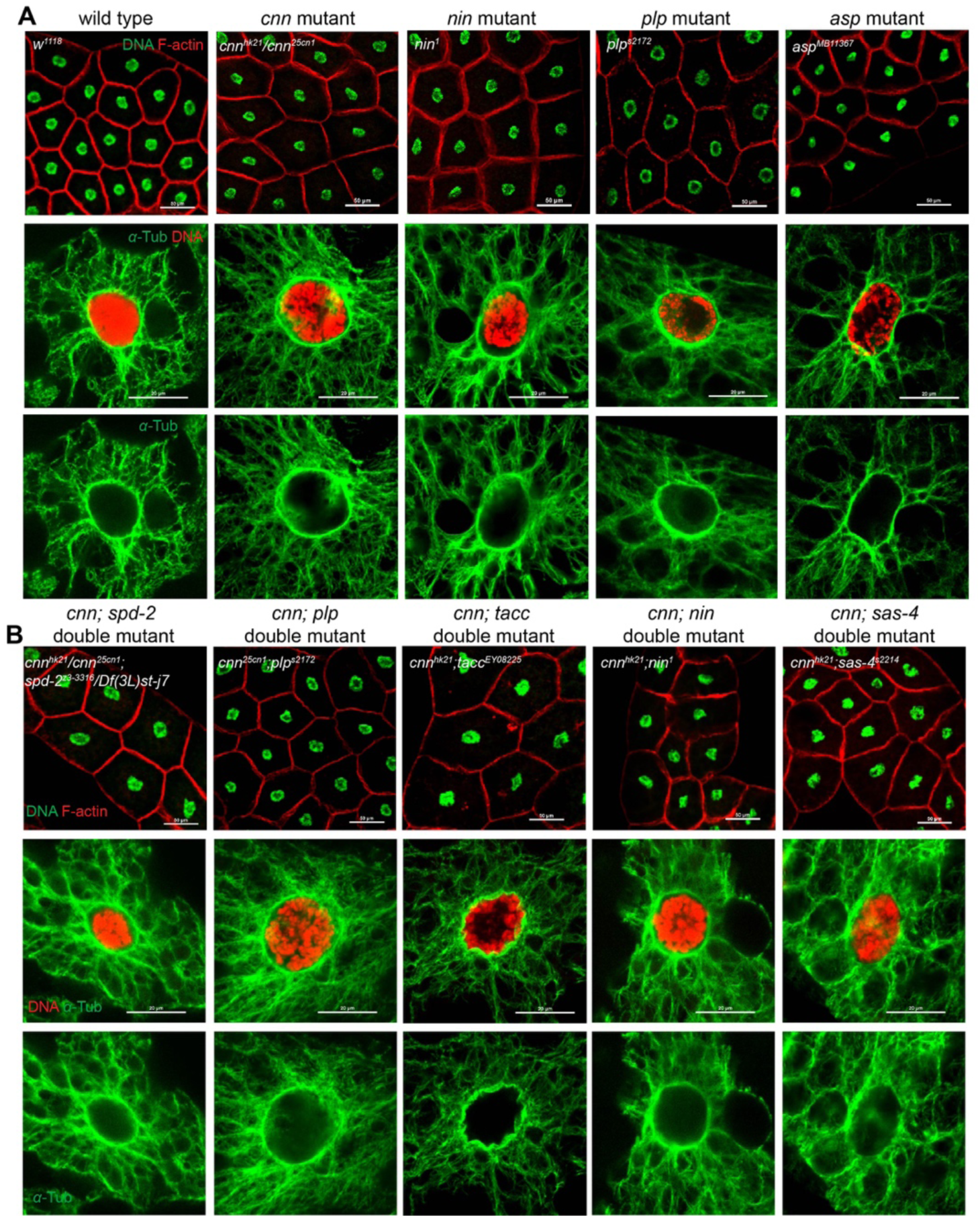
Single or double mutant combinations of centrosomal protein genes do not overtly impair MT assembly at the fat body ncMTOC. (A) Fat body cells were stained for DNA (Hoechst) and filamentous actin (CF568-Phalloidin) to assay for nuclear centricity. (B) MTs (anti-*α*-tubulin) and DNA (DAPI) were stained to assay MT organization. None of the single or double mutants tested showed defects in nuclear positioning or MT assembly at the MTOC. Quantifications of nuclear positioning (top panel) are shown in Figure 8B. Scale bars: 50 μm (top panels), 20 μm (middle and bottom panels).

### Nuclear positioning as a readout for MT assembly at the fat body ncMTOC

We next surveyed candidate MTOC components and MT regulators using nuclear positioning as a readout (Figure 8A), since nuclear positioning in the fat body requires integrity of the MT network. *α*-tubulin knockdown and Spastin overexpression were included for positive controls, where nuclear positioning was significantly disrupted (Figure 8B). Interestingly, when the MTOC was impaired, displaced nuclei showed little movement *in vivo* (Supplemental Movies 1 and 2).

**Figure 8.**
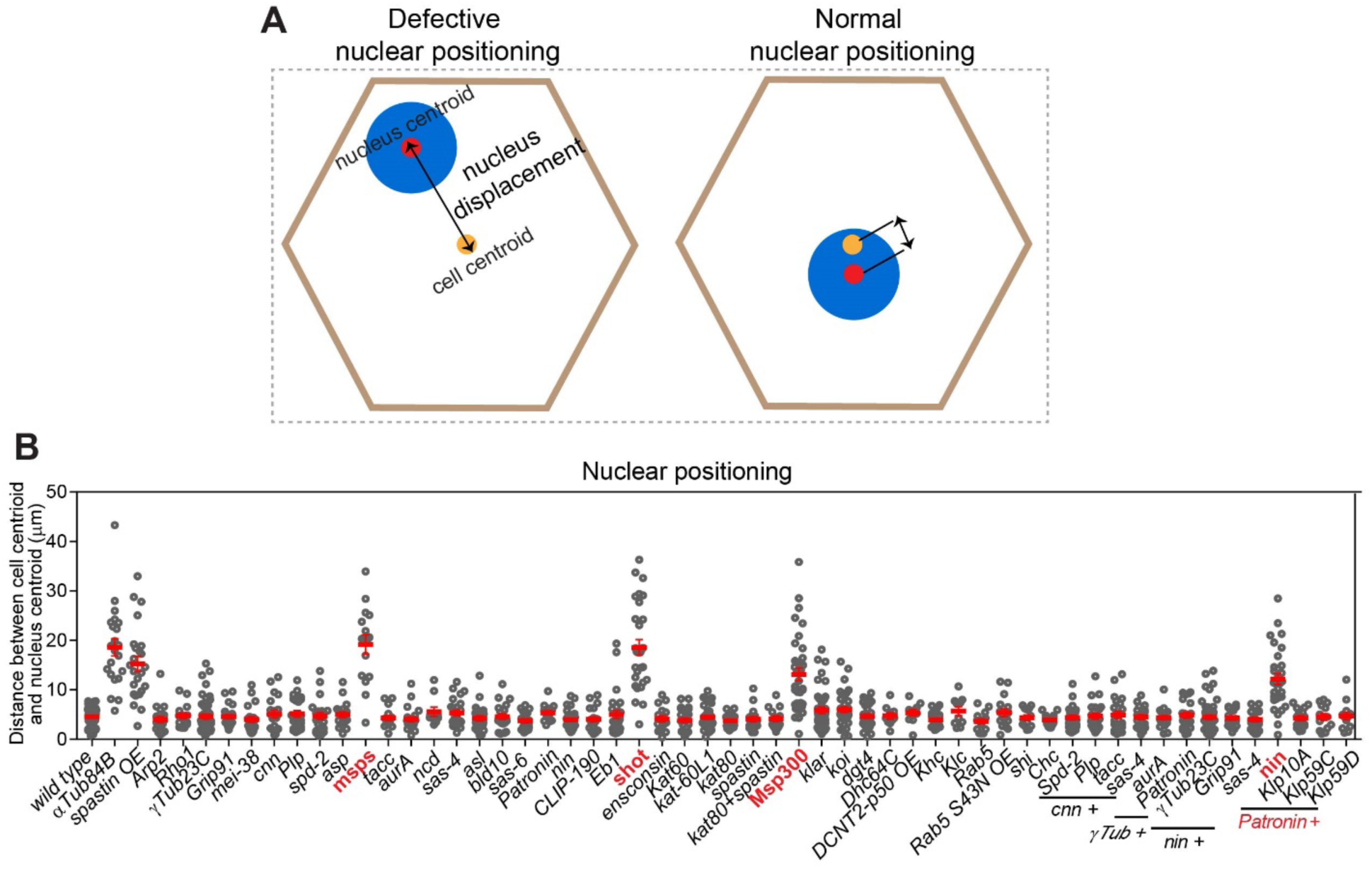
Nuclear positioning as a screen for MT assembly at the fat body ncMTOC. (**A**) Cartoon illustrating nuclear displacement measurements in a fat body cell with normal (left) or defective (right) nuclear positioning. (**B**) Quantification of nuclear positioning in the indicated genotypes. The positive hits are marked in red. Data are shown as the means ± s.e.m.

We screened most centrosomal genes and MT-regulatory genes and also included double combinations (see details below).

### *γ*-tubulin is not required for MT assembly at the fat body ncMTOC

*γ*-tubulin, partnering with *γ*-tubulin complex proteins (GCPs) to form the *γ*-tubulin ring complex (*γ*-TuRC), is widely employed as a MT nucleator or anchor at centrosomes and other MTOCs (Kollman et al., 2011; Lin et al., 2015; Oakley et al., 2015; Muroyama and Lechler, 2017; Sanchez and Feldman, 2017; Farache et al., 2018; Sallee et al., 2018; Tillery et al., 2018). Compared to the larval brain, the fat body expresses significantly less *γ*-tubulin, about 120-fold less (Figure 9A), but is detectable by IF and is localized to the ncMTOC throughout larval developmental stages (Figure 9C, D), together with other *γ*-TuRC components including GCP2 and GCP3 (Figure 6).

**Figure 9.**
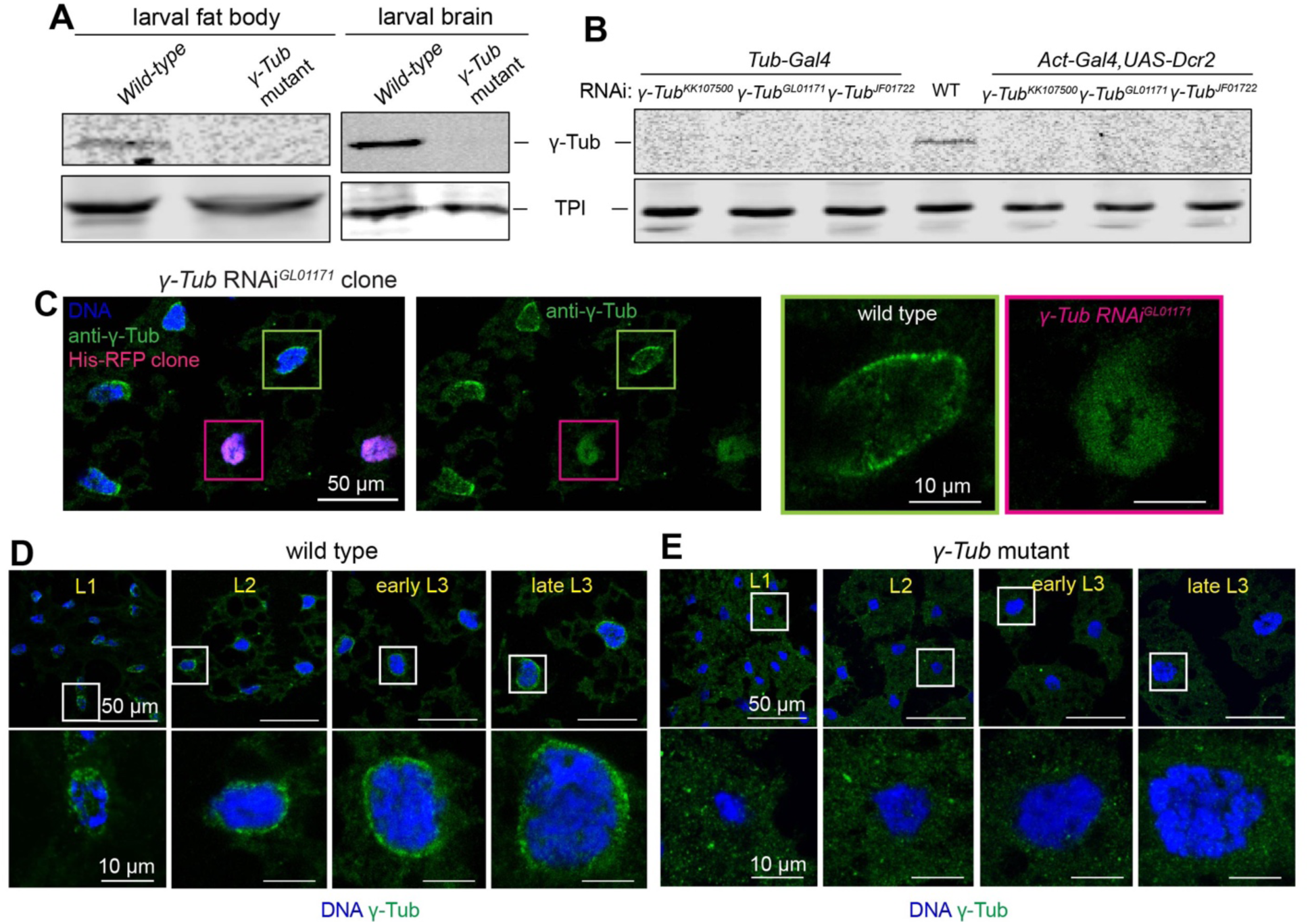
*γ*-tubulin is expressed and localized to perinuclear MTOC in fat body cells throughout larval developmental stages. (**A**) Western blot detection of *γ*-tubulin in wild-type and *γTub23C* mutant larval fat body lysates or larval brain lysates. Wild type is *w^1118^*, *γTub* mutant is *γ-Tub23C^A15-2^/Df (2L)JS17*. Triose-phosphate isomerase (TPI) is the loading control. (**B**) Western blot detection of *γ*-tubulin in whole larval lysates from wild-type and three independent *γTub23C* RNAi lines driven by two strong ubiquitous promoters: Tub-Gal4 and with Act5C-Gal4 plus UAS-Dicer-2 for increased knockdown efficiency. (**C**) Images showing *γ*-tubulin staining in *γ-Tub23C* ^GL01171^ RNAi fat body clone marked by His-RFP. Scale bar, 50 μm, zoomed images (10 μm) (**D**, **E**) Images showing g-tubulin staining in wild-type (**D**) and *γ-Tub23C^A15-2^/Df* mutant (**E**) fat body at different developmental stages as indicated. Scale bars: 50 μm (top panel), 10 μm (bottom panel).

To investigate the requirement for *γ*-tubulin at the fat body ncMTOC, we examined a null mutant and three independent RNAi lines, all of which showed depletion by western blotting and/or no detectable protein at the nuclear surface in the larval fat body throughout developmental stages (Figure 9). The lack of detectable signal in 1^st^ instar larvae indicates that the mutant effectively eliminated expression, and that there is no significant residual maternal supply in fat body cells prior to the third instar larval stage where most of our experiments were performed. Surprisingly, we found that fat body cells lacking *γ*-tubulin (*γTub23C*) or GCP3 (*grip91*) had normal nuclear positioning and MT assembly from the ncMTOC (Figures 10, 11, 15).

**Figure 10.**
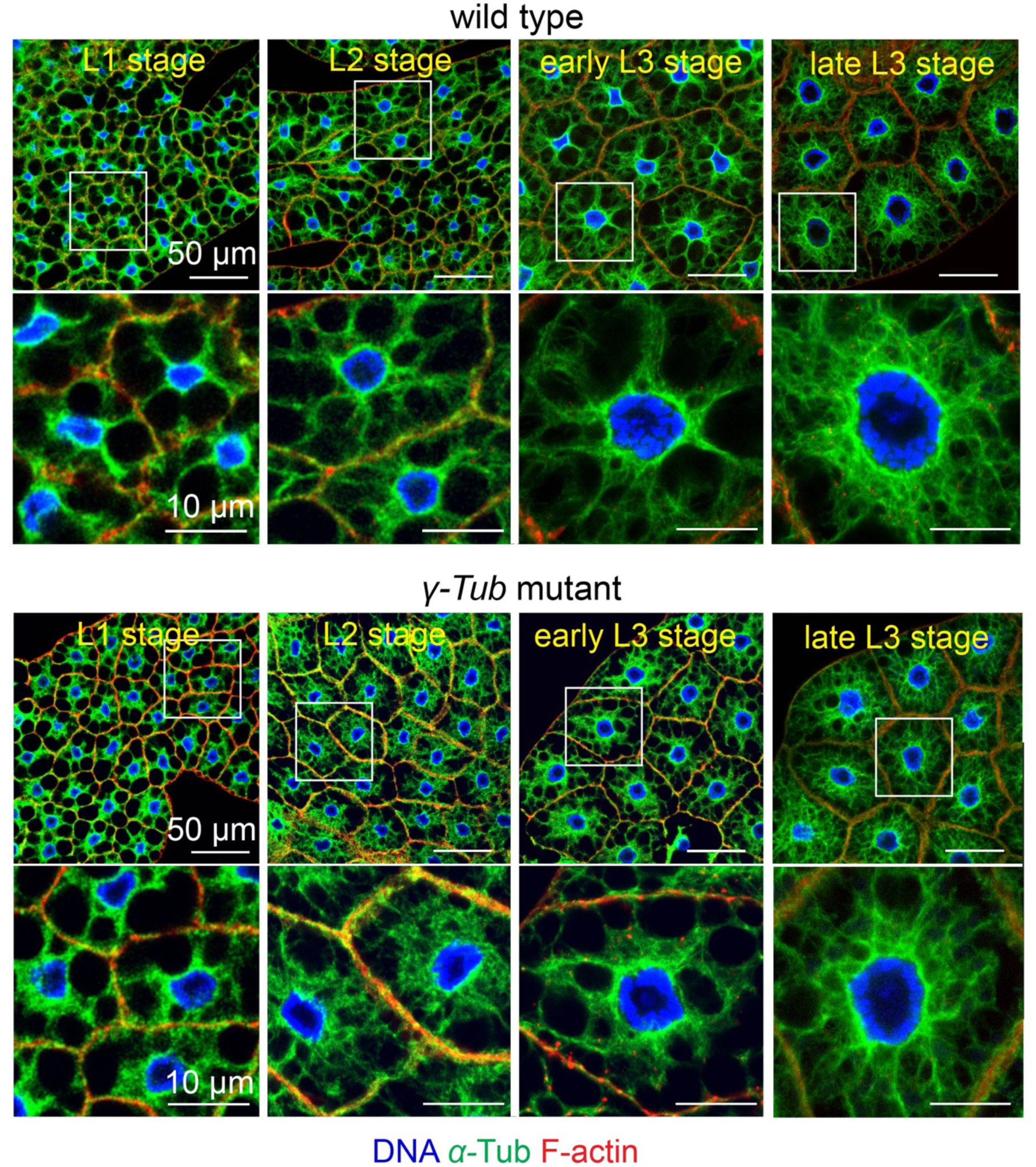
*γ*-tubulin is not required for MT assembly at the fat body ncMTOC. Images showing nuclear positioning and MT assembly in wild-type and *γ-Tub23C* mutant (*γ-Tub23C^A15-2^*/Df) fat bodies throughout 1^st^, 2^nd^, early 3^rd^ instar and late 3^rd^ instar larvae. Scale bars: 50 μm (top panel), 10 μm (bottom panel).

**Figure 11.**
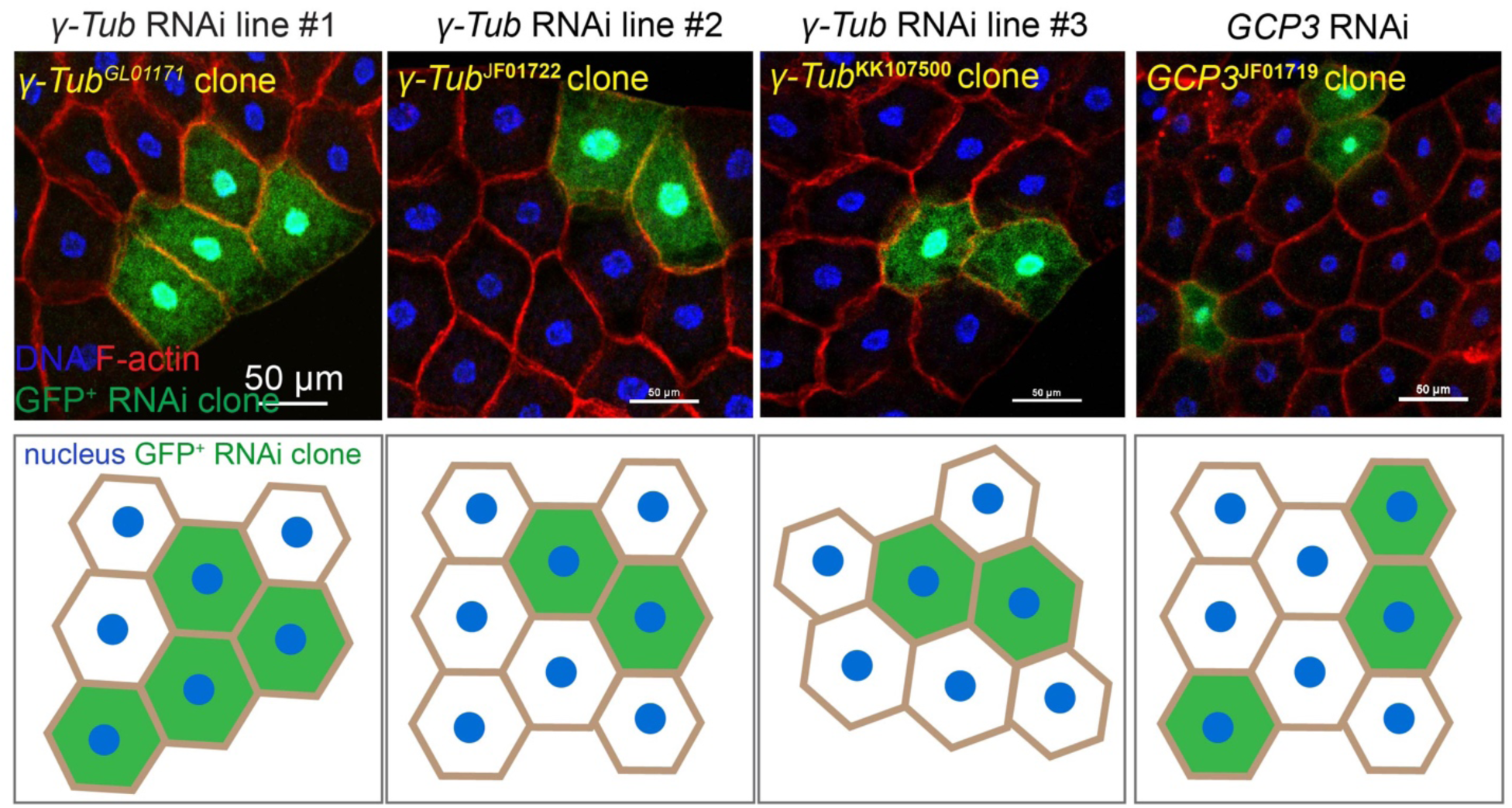
*γ*-tubulin is not required for nuclear positioning at the fat body cells. Images showing nuclear positioning in fat body clones with the indicated RNAi, including three independent *γ-Tub23C* RNAi lines and one *GCP3* RNAi line. The bottom panels are cartoons depicting the nuclear positioning in the indicated RNAi clones. Scale bars: 50 μm.

Since fat body MTs are highly stabilized, a requirement for *γ*-tubulin during early stages of fat body MTOC, when some residual maternal supply might persist, could be masked by MT stabilization in the *γ*-tubulin mutant. To address this possibility, we performed MT regrowth in *γ*-tubulin-depleted RNAi clones, and compared MT regrowth in wild type and RNAi cells side-by-side in late (third instar) stages. We found that MT regrowth was indistinguishable between wild-type and *γ*-tubulin-depleted fat body cells (Figure 12), indicating that *γ*-tubulin has no significant role in MT assembly at the fat body ncMTOC. Together, these results show that *γ*-tubulin is not required for the assembly of MTs at the fat body ncMTOC.

**Figure 12.**
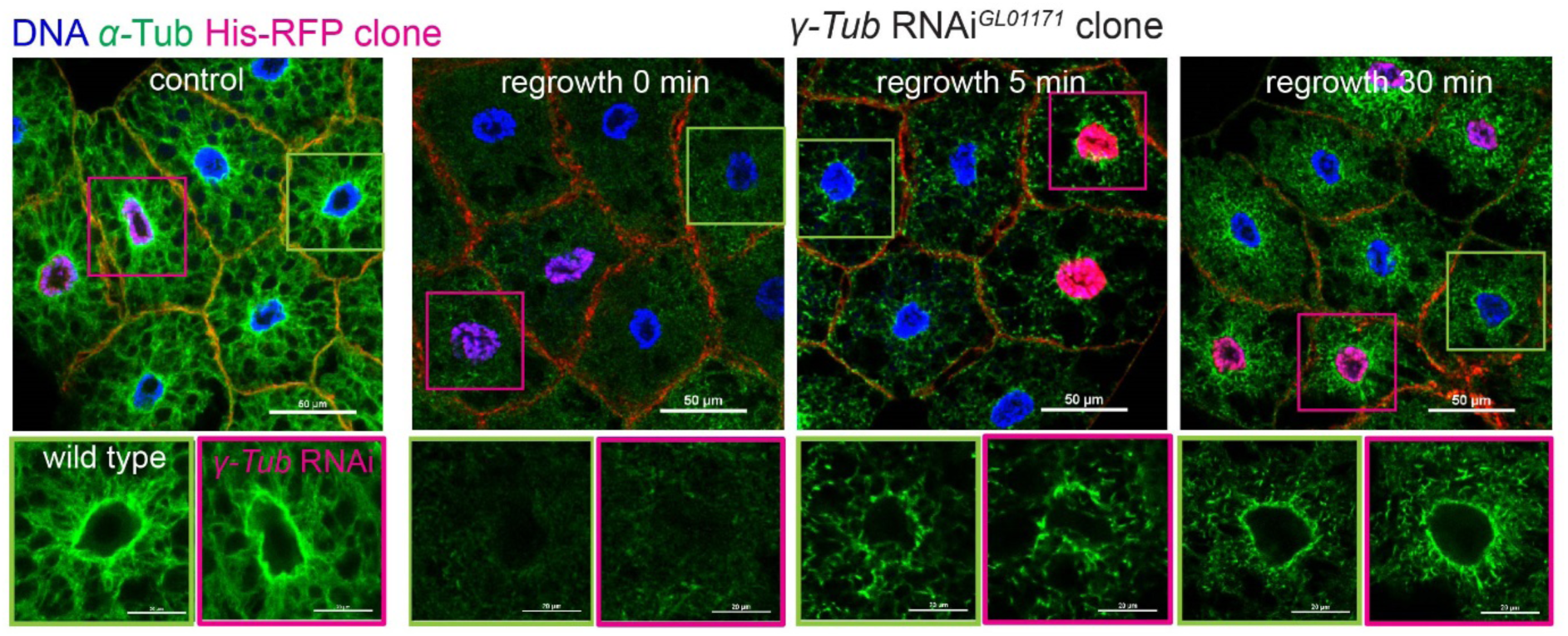
MT regrowth is normal in the absence of *γ*-tubulin at the fat body ncMTOC. Images showing MT regrowth in wild-type and *γ-Tub23C* ^GL01171^ RNAi clones marked by His-RFP. Control: no vinblastine treatment; the other three groups were treated with vinblastine and underwent MT regrowth for 0, 5, 30 min as indicated. Scale bars: 50 μm (top panel), 20 μm (bottom panel).

*γ*-tubulin is the major regulator of centrosomal MTs while AurA independently regulates the remainder (Strome et al., 2001; Hannak et al., 2002; Motegi et al., 2006). However, co-depletion of *γTub23C* and *aurA* had no effects on fat body nuclear positioning and ncMTOC assembly (Figure 13), consistent with a similar lack of requirement for these MT regulators at the apical ncMTOC in *C. elegans* embryonic intestinal epithelial cells (Sallee et al., 2018). Therefore, the widespread MT nucleator *γ*-tubulin is not required for MT assembly at the fat body ncMTOC.

**Figure 13.**
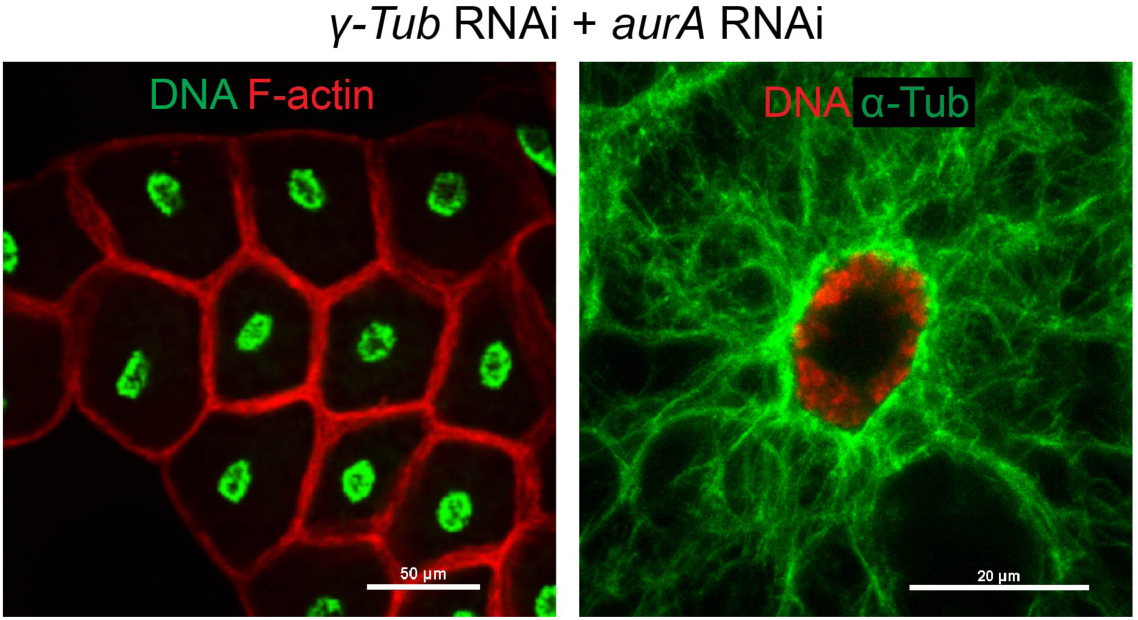
Co-depletion of *γ*-tubulin and AurA does not affect fat body ncMTOC. Images showing nuclear positioning (left panel) and MT assembly (right panel) in fat body after double knockdown of *γTub23C* and *aurA*. Scale bars: 50 μm (left), 20 μm (right).

### Msps is required for the fat body ncMTOC assembly independent of conventional partners

Among the 23 mutants or RNAi knockdowns tested in our survey for regulators of the fat body ncMTOC, *minispindles* (*msps*) RNAi knockdown (Figure 14B, C) singularly caused a severe defect in nuclear centricity (Figure 14D), indicating that Msps is essential for fat body ncMTOC activity. Msps, the homolog of XMAP215/Stu2/Dis1/Alp14/ZYG-9/ch-TOG, functions at the growing MT plus end as a processive MT polymerase (Brouhard et al., 2008; Al-Bassam and Chang, 2011; Wieczorek et al., 2015; Geyer et al., 2018). Msps localizes to the fat body perinuclear MTOC (Figure 14A, Table 1) and *msps* knockdown had a severe impact on radial MT assembly at the fat body MTOC. Significantly, the dense circumferential MTs at the nuclear surface appeared unaffected, and only the MTs that radiate outward from the MTOC were diminished (Figure 15), indicating that MT elongation is specifically impaired.

**Figure 14.**
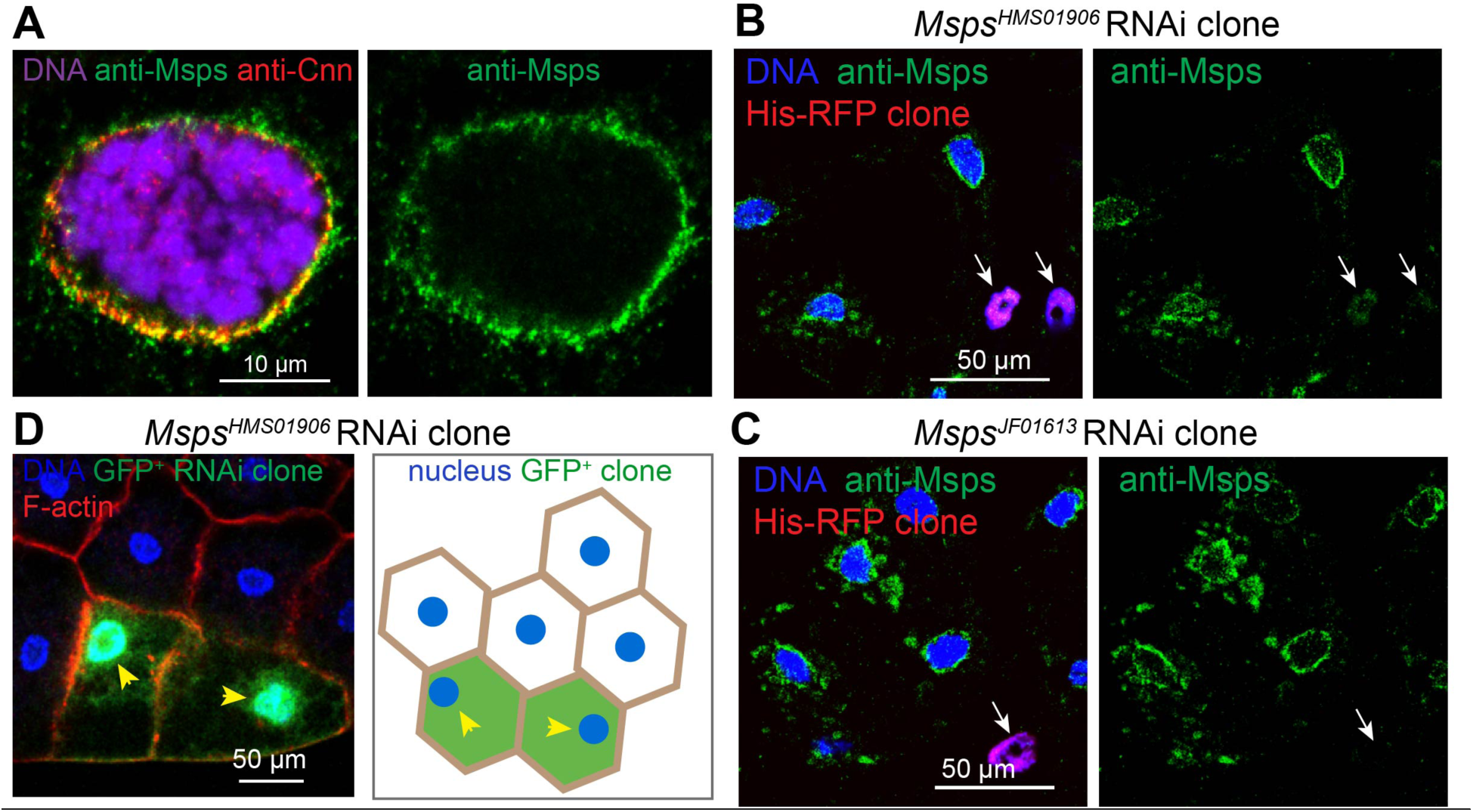
Msps is essential for nuclear positioning. (**A**) Msps is localized to perinuclear site in fat body. Images showing Msps staining in wildtype fat body cells that are co-stained with Cnn and DAPI. Scale bar, 10 μm. (**B**, **C**) Validation of two independent Msps RNAi lines. Staining for Msps in *msps* RNAi fat body clones marked by His-RFP. Arrow indicates the loss of perinuclear Msps signal in an *msps* RNAi cell. Scale bars: 50 μm. (**D**) Msps RNAi GFP clones show defective nuclear positioning. Left panel: fat bodies containing control and GFP-marked knockdown clones were co-stained for DNA (Hoechst) and actin (CF568-Phalloidin) to assess nuclear centricity. Right panel: cartoon depicting nuclear positioning in the Msps RNAi clones. Arrowheads indicate a loss of nuclear centricity. Quantification of nuclear positioning is shown in Figure 8B. Scale bar, 50 μm.

**Figure 15.**
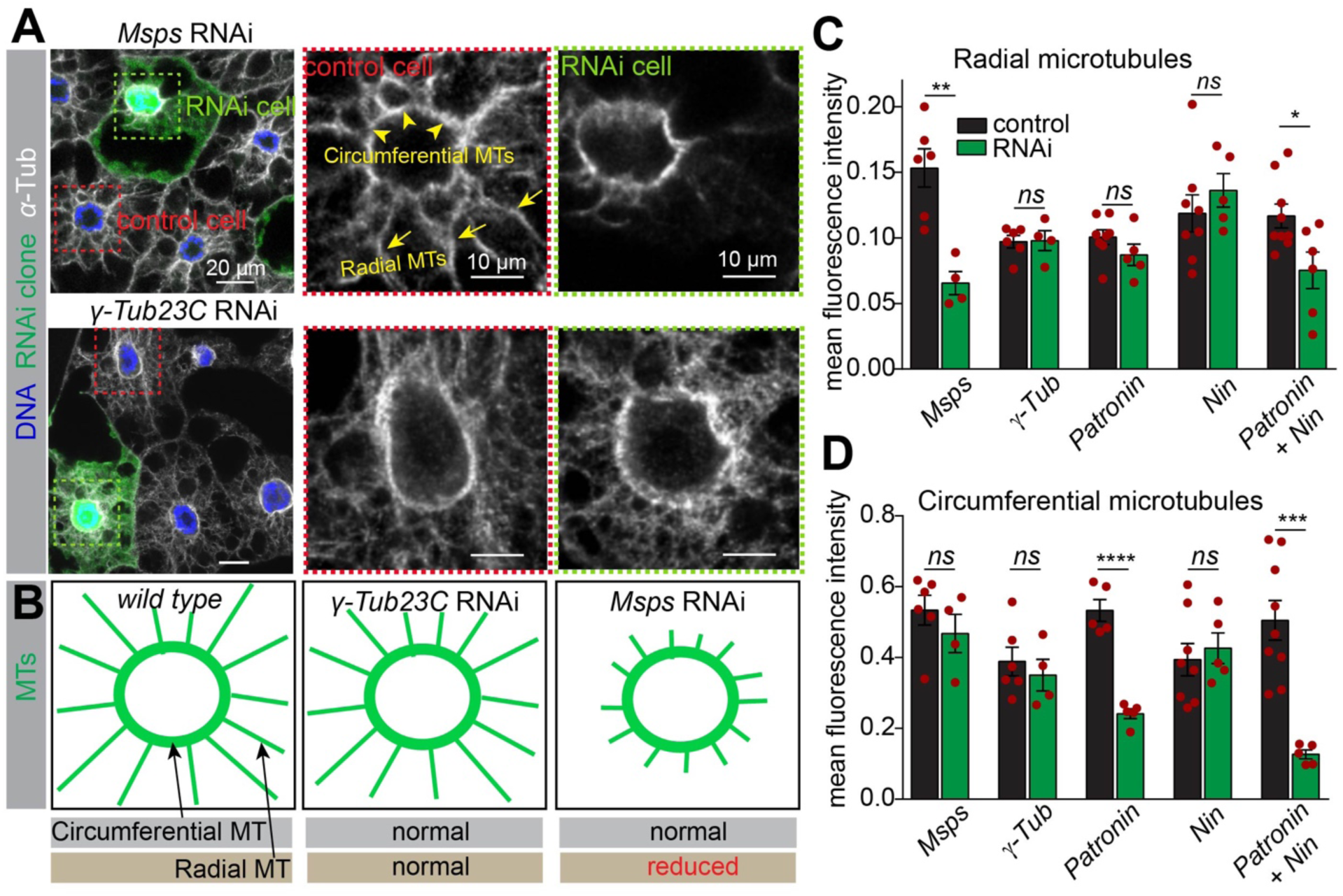
Msps is required for MT growth independently of *γ*-tubulin. (**A**) Msps depletion disrupts radial MTs, but not circumferential MTs, while *γ*-tubulin depletion does not affect either class of MT. Mosaic fat bodies were stained for MTs (anti-α-tubulin) and DNA to assess MT assembly defects at the perinuclear ncMTOC. Control and GFP^+^ RNAi knockdown cells were boxed with dashed red and green lines, respectively in the left panels. Middle and right panels show close-up images of the boxed MT staining in the control and RNAi cells. Circumferential MTs (arrowheads) and radial MTs (arrows) are indicated in a control cell. Scale bars: 20 μm (left panel), 10 μm (middle and right panels). (**B**) Diagrams depicting circumferential and radial MT assembly phenotypes in fat body cells from wild type and the indicated RNAi knockdowns. (**C, D**) Quantification of mean fluorescent intensity of radial microtubules (**C**) and circumferential microtubules (**D**) in the indicated RNAi knockdown clones compared to internal control cells. Data are shown as the means ± s.e.m. Statistics by two-tailed Student’s *t*-test. *msps* RNAi vs control, *p*=0.002 (**, radial), *p*=0.3572 (ns, circumferential); *γ-Tub23C* RNAi vs control, *p*=0. 9316 (ns, radial), *p*=0.5535 (ns, circumferential); *Patronin* RNAi vs control, *p*=0.1845 (ns, radial), *p*<0.0001 (****, circumferential); *Nin* vs control, *p*=0.4156 (ns, radial), *p*=0.6417 (ns, circumferential); *Patronin* RNAi + *Nin* RNAi vs control, *p*=0.0213 (*, radial), *p*=0.0004 (***, circumferential).

Tacc is a major protein partner of Msps, and its localization at centrosomes is regulated by Aurora A kinase (Cullen and Ohkura, 2001; Lee et al., 2001; Barros et al., 2005). In oocytes, Msps is transported to spindle poles by the Kinesin-14 motor Ncd (Cullen and Ohkura, 2001; Lee et al., 2001; Barros et al., 2005). However, none of these components are required for Msps recruitment, nor ncMTOC assembly or nuclear centricity in the fat body (Figure 16), indicating a distinct mechanism for Msps deployment at the fat body ncMTOC.

**Figure 16.**
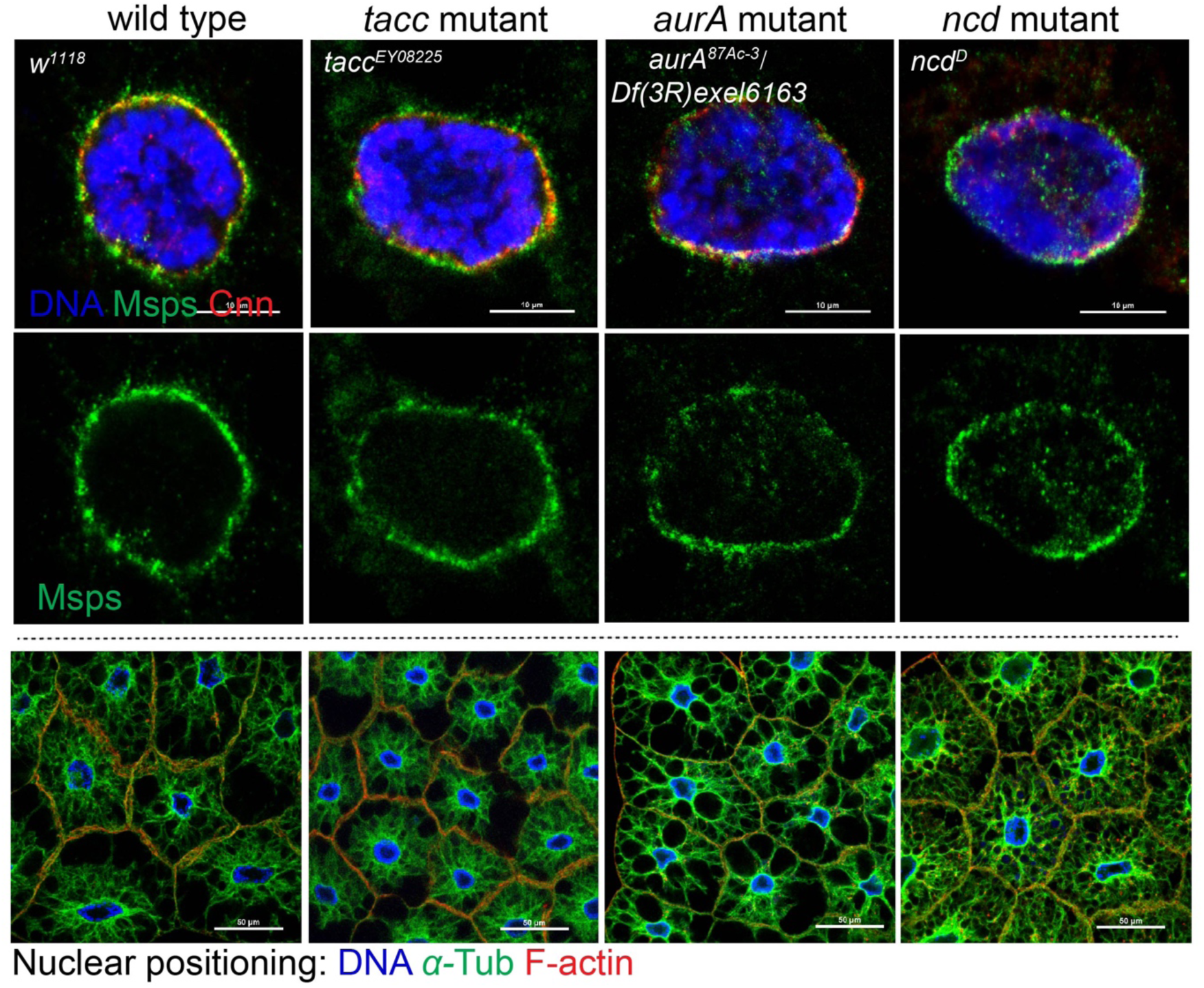
Msps localization at the fat body ncMTOC does not require known regulators, including TACC, AurA and Ncd. Msps localization at centrosomes or spindle poles relies on Tacc, Aurora A kinase phosphorylation of Tacc, and the Kinesin-14 motor Ncd. None of these components are required for Msps localization at the fat body ncMTOC (top and middle panels), or for fat body nuclear positioning and MT organization (bottom panels). Fat bodies were stained with antibodies against Msps and Cnn, or for MTs using a FITC-conjugated DM1A antibody, and for DNA with DAPI, and actin with CF568-conjugated phalloidin. Nuclear positioning was quantified in Figure 8B. Scale bars: 10 μm (top and middle panels), 50 μm (bottom panels).

It was recently shown that the Msps homologs XMAP215, Stu2, and Alp14 function as MT nucleators by directly binding to and cooperating with *γ*-tubulin complexes for MT nucleation (Flor-Parra et al., 2018; Gunzelmann et al., 2018; Thawani et al., 2018). However, the role for Msps in the fat body ncMTOC is clearly distinct from the cooperative one it plays with the *γ*-TuRC in these other contexts because the *γ*-TuRC is not required for the fat body ncMTOC. Previous work supports a role for Msps and Stu2 in MT growth independent of *γ*-tubulin in interphase S2 cells and yeast, respectively (Rogers et al., 2008; Gunzelmann et al., 2018).

Together, these data show that Msps is required for MT assembly at the fat body ncMTOC, specifically for radial MT elongation, independently of previously described mechanisms, including the recently identified *γ*-tubulin–dependent nucleation seen with Msps homologs.

### Patronin controls circumferential MT assembly at the ncMTOC

Patronin/CAMSAP family proteins are MT minus-end-associated proteins that stabilize MTs and have emerged as critical MT assembly factors at ncMTOCs in a variety of organisms and cell types (Wu and Akhmanova, 2017). Patronin localized at the fat body MTOC (Figure 17A, Table 1), however, its knockdown (Figure 17B) had no significant impact on nuclear centricity (Figure 8B, 17C), although the MT array was generally reduced in Patronin knockdown clones (Figure 17D). Interestingly, in Patronin knockdown fat body cells, examined in three independent RNAi lines, the circumferential MTs were significantly affected but the radial MTs were not (Figure 17D, F, and quantification in Figure 15C, D), suggesting that Patronin is involved in stabilization or assembly of the MTs most proximal to the MTOC.

**Figure 17.**
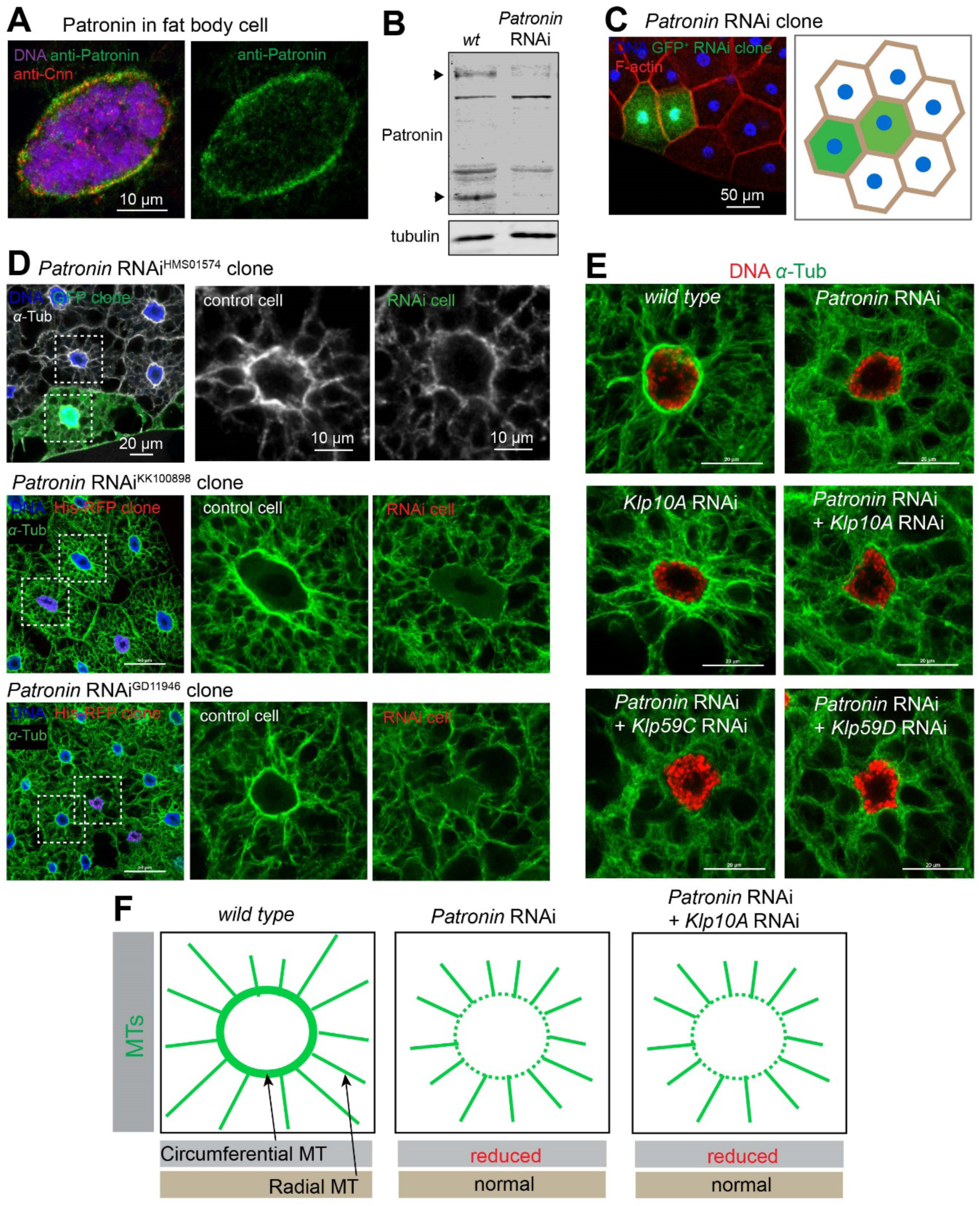
Patronin is required for assembly of circumferential MTs at the fat body ncMTOC. (**A**) Patronin is perinuclear in fat body cells. Images of fat body cells stained for Cnn in red and counterstained in green for Patronin by antibody staining. Scale bar, 10 µm. (**B**) Western blot of Patronin in larval fat body lysates from wild-type and *Patronin* RNAi (HMS01574) knockdown with the SPARC-Gal4 fat body driver. Arrows indicate the expected Patronin isoform band of approximately 180 kDa, and another putative isoform at approximately 70 kDa. (**C**) Patronin RNAi (HMS01574) clones show normal nuclear positioning. Left panel: fat bodies containing control and GFP+ knockdown clones were co-stained for DNA (Hoechst) and actin (CF568-Phalloidin) to assess nuclear centricity. Right panel: illustration depicting nuclear positioning in the Patronin RNAi clones. Quantification is shown in Figure 8B. Scale bar, 50 µm. (**D**) Three independent RNAi lines show that Patronin depletion disrupts circumferential MTs, not radial MTs. Mosaic fat bodies were stained for MTs (anti-*α*-tubulin) and DNA to assess MT assembly defects at the perinuclear ncMTOC. Middle and right panels showed close-up images of the MT staining in the control and RNAi cells. MT quantification is shown in Figure 15C, D. (**E**) Patronin regulation of circumferential MT assembly appears not antagonized by Kinesin-13 MT depolymerases. Images show MT staining in wild-type, single knockdown of Patronin or the MT depolymerase Klp10A (Kinesin-13), and double knockdowns of Patronin and Klp10A, and of Patronin and with other MT depolymerase paralogs *Klp59C* or *Klp59D*. Scale bars: 20 µm. (**F**) Diagram depicting changes in the two classes of MTs in the indicated groups.

How Patronin functions to assemble MTs at the ncMTOC remains incompletely understood. Patronin/CAMSAP stabilizes MT minus-ends and antagonizes the activity of Kinesin-13 family depolymerases (Goodwin and Vale, 2010; Hendershott and Vale, 2014; Atherton et al., 2017). However, this antagonism does not prevail at the fat body ncMTOC because Kinesin-13 (*Klp10A*) knockdown, or other MT depolymerases *Klp59C* or *Klp59D* did not suppress the reduced circumferential MTs in Patronin knockdown cells (Figure 17E, F).

CAMSAP/Patronin associates with the MT severing enzyme Katanin (Jiang et al., 2014; Nashchekin et al., 2016; Jiang et al., 2018), an association proposed to remodel MTs with the ability to amplify MTs (Roll-Mecak and Vale, 2006; Sharp and Ross, 2012; Lindeboom et al., 2013; Vemu et al., 2018; Kuo et al., 2019). However, in *Drosophila* fat body cells, loss of MT severing enzymes katanin p60 (kat60), katanin p60-like 1(kat-60L1), the katanin regulatory subunit p80 (kat80) or spastin, the other major MT severing enzyme, did not overtly impact perinuclear MT assembly or nuclear positioning (Figure 18A, B). The co-depletion of katanin and spastin also had no impact (Figure 18B).

**Figure 18.**
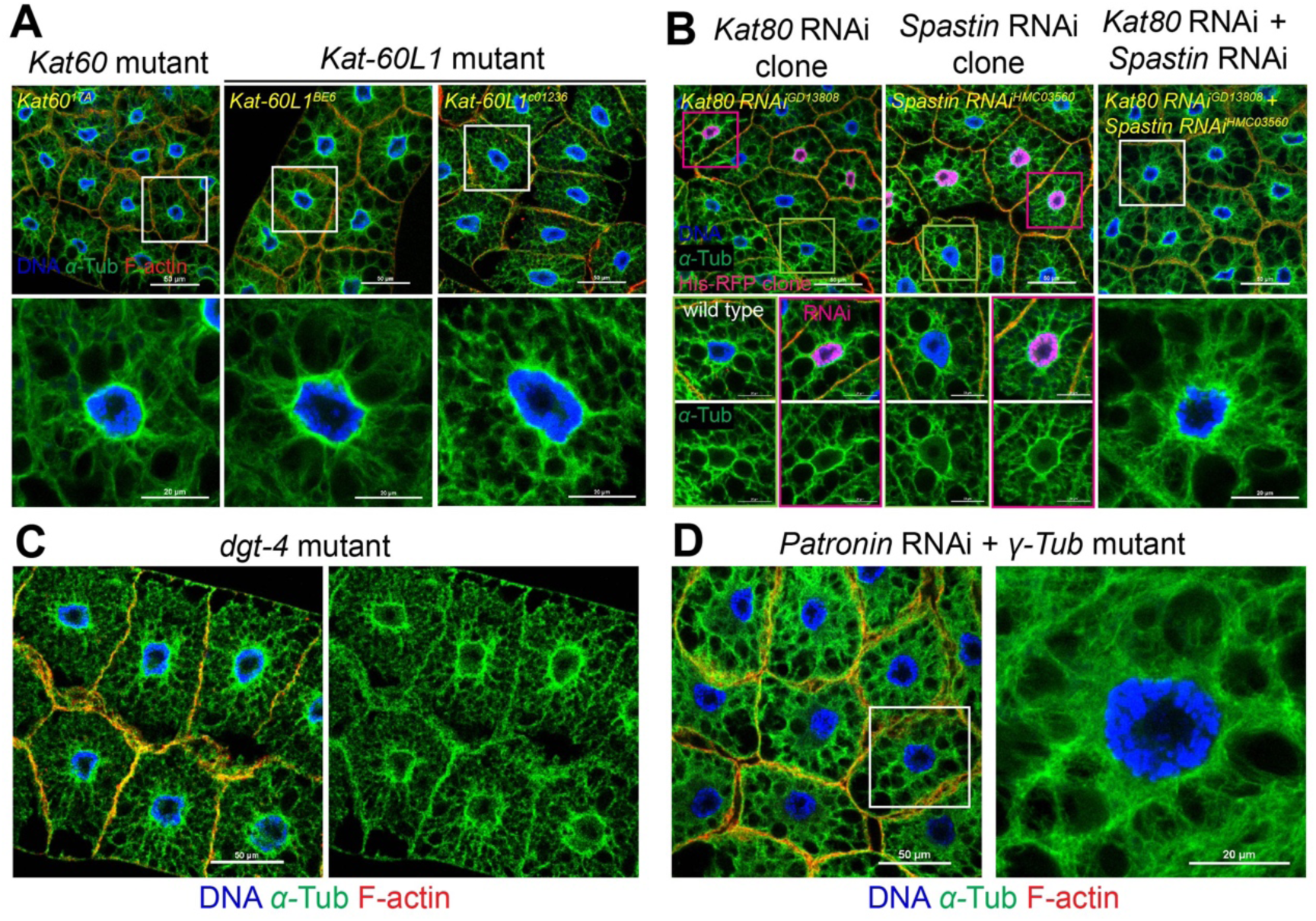
MT severing and amplification mechanisms do not prevail at fat body. (**A**) Mutants for MT severing enzymes katanin p60 (*Kat60^17A^*) or katanin p60-like 1 (*Kat-60L1^BE6^*, *Kat-60L1^c01236^*) do not affect nuclear positioning (top) or MT assembly (bottom). Nuclear quantification is shown in Figure 8B. Scale bar, 50 µm (top panels), 20 µm (bottom panels). (**B**) Knockdown of MT severing enzymes *Kat80* or *Spastin* in GFP-marked clones or double knockdown of *Kat80* and *Spastin* (not assayed in clones) does not affect nuclear positioning (top) or MT organization (bottom). Nuclear quantification is shown in Figure 8B. Scale bar, 50 μm (top panel), 20 μm (middle and bottom panels). (**C**) Mutant for *dgt4*, an augmin complex subunit, does not affect nuclear positioning (left) orMT assembly (right). Nuclear quantification is shown in Figure 8B. Scale bar, 50 μm. (**D**) *γTub23C* mutation (*γTub23C^A15-2^/Df*) together with *Patronin* RNAi knockdown does not affect nuclear positioning (left) or radial MT assembly (right). Nuclear quantification is shown in Figure 8B. Scale bar, 50 µm (left), 20 µm (right).

Another proposed MT amplification mechanism involves the augmin/HAUS complex, which generates new MTs along pre-existing ones through *γ*-TuRC-mediated nucleation and promotes mitotic spindle MT assembly (Goshima et al., 2008; Lawo et al., 2009) or post-mitotic non-centrosomal MT assembly (Sanchez-Huertas et al., 2016; Cunha-Ferreira et al., 2018). We tested a mutant for *dgt4*, a core augmin complex subunit, and found no defects in nuclear centricity or perinuclear MT assembly in fat body cells (Figure 18C). These data indicate that these two potential MT amplification mechanisms are not functionally required for MT assembly at the fat body ncMTOC.

Taken together, these results show that the ncMTOC regulator Patronin/CAMSAP is partially responsible for MT assembly at the fat body MTOC, but does not function via the known mechanisms established in other contexts, including the reported antagonism with Kinesin-13 and cooperativity with the severing enzyme Katanin.

### Patronin cooperates with Ninein to organize the fat body ncMTOC by recruiting Msps

We reasoned that Patronin might work cooperatively or in parallel with other minus-end proteins to assemble the fat body ncMTOC. We investigated *γ*-tubulin and Ninein (Nin), two other MT minus-end proteins. CAMSAP and *γ*-tubulin are proposed to act sequentially in the generation of non-centrosomal MTs in neurons where *γ*-tubulin initiates MT nucleation and CAMSAP stabilizes MTs (Yau et al., 2014). In the *C. elegans* larval epidermis, Patronin works in parallel with *γ*-tubulin at ncMTOCs (Wang et al., 2015b). In *Drosophila* fat body cells, however, co-depletion of *Patronin* and *γTub23C* did not enhance the MT-organization phenotype of *Patronin* single knockdown (Figure 18D), indicating that *γ*-tubulin does not function redundantly with Patronin in MT assembly at the fat body ncMTOC.

Nin is a MT minus-end protein with anchoring function at the centriole subdistal appendages in mammals (Mogensen et al., 2000; Delgehyr et al., 2005), but Nin also has ncMTOC roles in mammals, *Drosophila* and *C. elegans* (Mogensen et al., 2000; Wang et al., 2015b; Zheng et al., 2016; Goldspink et al., 2017). While Nin’s molecular mechanisms of action are not well understood, the conserved N-terminal domain of *Drosophila* Nin can associate directly with MTs (Kowanda et al., 2016). Nin localizes to *Drosophila* muscle and wing ncMTOCs (Zheng et al., 2016) and to the apparent ncMTOC in oocytes (Kowanda et al., 2016). We found that Nin localized to the fat body perinuclear ncMTOC (Figure 19A). Mutations in *ninein* are viable in *Drosophila* and mouse (Kowanda et al., 2016; Zheng et al., 2016; Lecland et al., 2019; Rosen et al., 2019), and *Nin* mutants or knockdowns (Figure 19B, C) showed no overt effects on fat body nuclear positioning (Figure 19D) or MT assembly (Figure 19E, E’, and quantification in Figure 15C, D).

**Figure 19.**
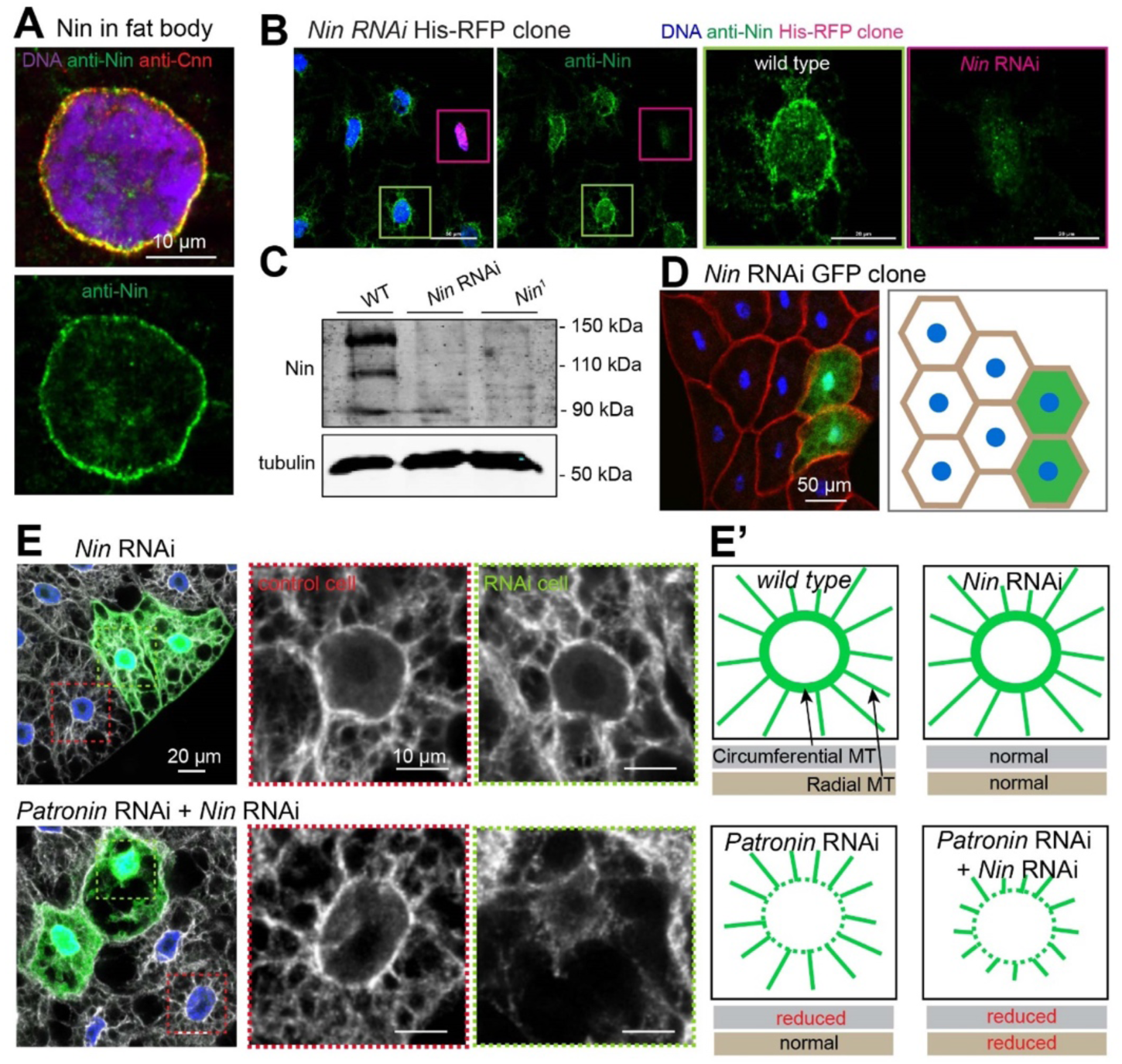
Nin alone is not essential, but its cooperation with Patronin is essential for MT assembly at the fat body ncMTOC. (**A**) Nin is perinuclear in fat body cells. Images of fat body cells stained with antibodies against Cnn in red and Nin in green. Scale bar, 10 μm. (**B**) Verification of *Nin* RNAi by staining. Images showing Nin staining in a *Nin*^HMJ23837^ RNAi fat body clone marked by His-RFP. Scale bars: 50 μm (left two images), 20 μm (right two images). (**C**) Verification of *Nin* RNAi and mutant. Western blot detection of Nin in larval brain lysates from wild-type, *Nin^1^* (null) mutant, and *Nin*^HMJ23837^ RNAi knockdown driven by Tub-Gal4. (**D**) Nin RNAi GFP+ clones show normal nuclear positioning. Left panel: fat bodies containing control and GFP+ knockdown clones were co-stained for DNA (Hoechst) and actin (CF568-Phalloidin) to assess nuclear centricity. Right panel: illustration depicting nuclear positioning in the *Nin* RNAi clones. Quantification shown in Figure 8B. Scale bar, 50 μm. (**E, E’**) Nin depletion disrupts neither circumferential nor radial MTs, while co-depletion of Patronin and Nin disrupts both classes of MTs. Mosaic fat bodies were stained for MTs (anti-α-tubulin) and DNA to assess MT assembly defects at the perinuclear ncMTOC. Control and GFP^+^ RNAi knockdown cells were boxed with dashed red and green lines, respectively. Middle and right panels show close-up images of MT staining in the control and RNAi cells. Diagram in E’ depicts changes in two classes of MTs after indicated RNAi. MT quantification is shown in Figure 15C, D. Scale bars: 20 μm (left panel), 10 μm (middle and right panels).

However, double knockdown of *Patronin* and *Nin* significantly disrupted nuclear positioning (Figure 19E, E’). Moreover, *Patronin Nin* double knockdown impaired both radial and circumferential MT assembly, while *Patronin* knockdown mostly affected circumferential MTs and *Nin* knockdown had no overt effects on its own (Figure 19E, E, also see quantification in Figure 15C, D). These data demonstrate that Patronin and Nin function redundantly in MT assembly at the fat body ncMTOC and indicates that each can effectively compensate for the other to control assembly of radial MTs.

Nin associates with *γ*-tubulin (Delgehyr et al., 2005; Zheng et al., 2016). In *C. elegans*, *γ*-tubulin recruits NOCA-1 (a *nin* homolog) to the ncMTOC in epidermal cells, where they function together and in parallel to Patronin to regulate non-centrosomal MT assembly (Wang et al., 2015b). However, at the fat body ncMTOC, *γ*-tubulin is not required to recruit Nin (Figure 20A), and *γTub23C* knockdown or mutant did not enhance *Patronin* knockdown phenotypes (Figure 18D). Moreover, *Nin* knockdown together with *γTub23C* or *gcp3* had no effect on nuclear positioning in fat body cells (Figure 20B), indicating that Nin and *γ*-tubulin/*γ*-TuRC do not act in a parallel or cooperative manner at the fat body ncMTOC. Altogether, these findings show that Nin and Patronin cooperate in MT assembly at the ncMTOC without functional involvement of *γ*-tubulin.

**Figure 20.**
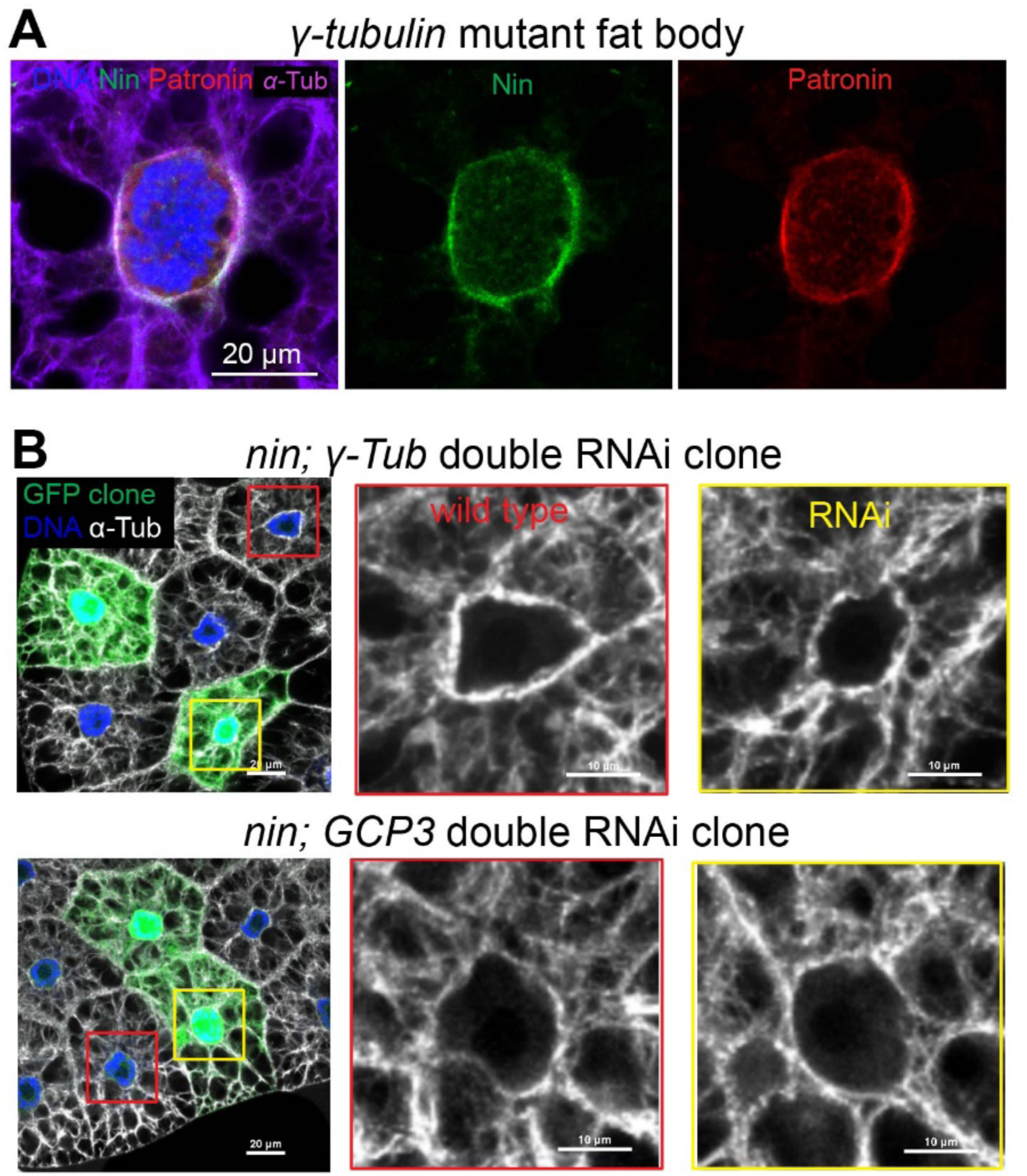
Nin does not cooperate with *γ*-tubulin at the fat body ncMTOC. (**A**) *γ*-tubulin does not recruit Nin or Patronin. Staining for Nin (green), Patronin (red) and MTs (purple) in *γTub23C^A15-2^/Df* mutant larval fat body. Scale bar, 20 μm. (**B**) Double knockdown of *Nin* and *γTub23C,* or *Nin* and *GCP3* does not affect nuclear positioning and MT assembly in fat body clones. Nuclear quantification is shown in Figure 8B. Scale bar, 20 μm (left panel), 10 μm (middle and right panels)

Since Patronin and Ninein are required for assembly of both the radial and circumferential MTs at the fat body MTOC, and because Msps depletion only affects radial MTs, we examined whether Patronin and Ninein are required to recruit Msps to the ncMTOC. Knockdown of *Patronin* reduced Msps recruitment to the nuclear surface to 37.0% of control, while *Nin* knockdown had no significant impact (Figure 21A, B). Knockdown of *Patronin* and *Nin* together, however, significantly diminished Msps localization to 19.6% relative to control (Figure 21A, B). We used CoIP assays to further show that Patronin associates with Msps in a protein complex (Figure 21C).

**Figure 21.**
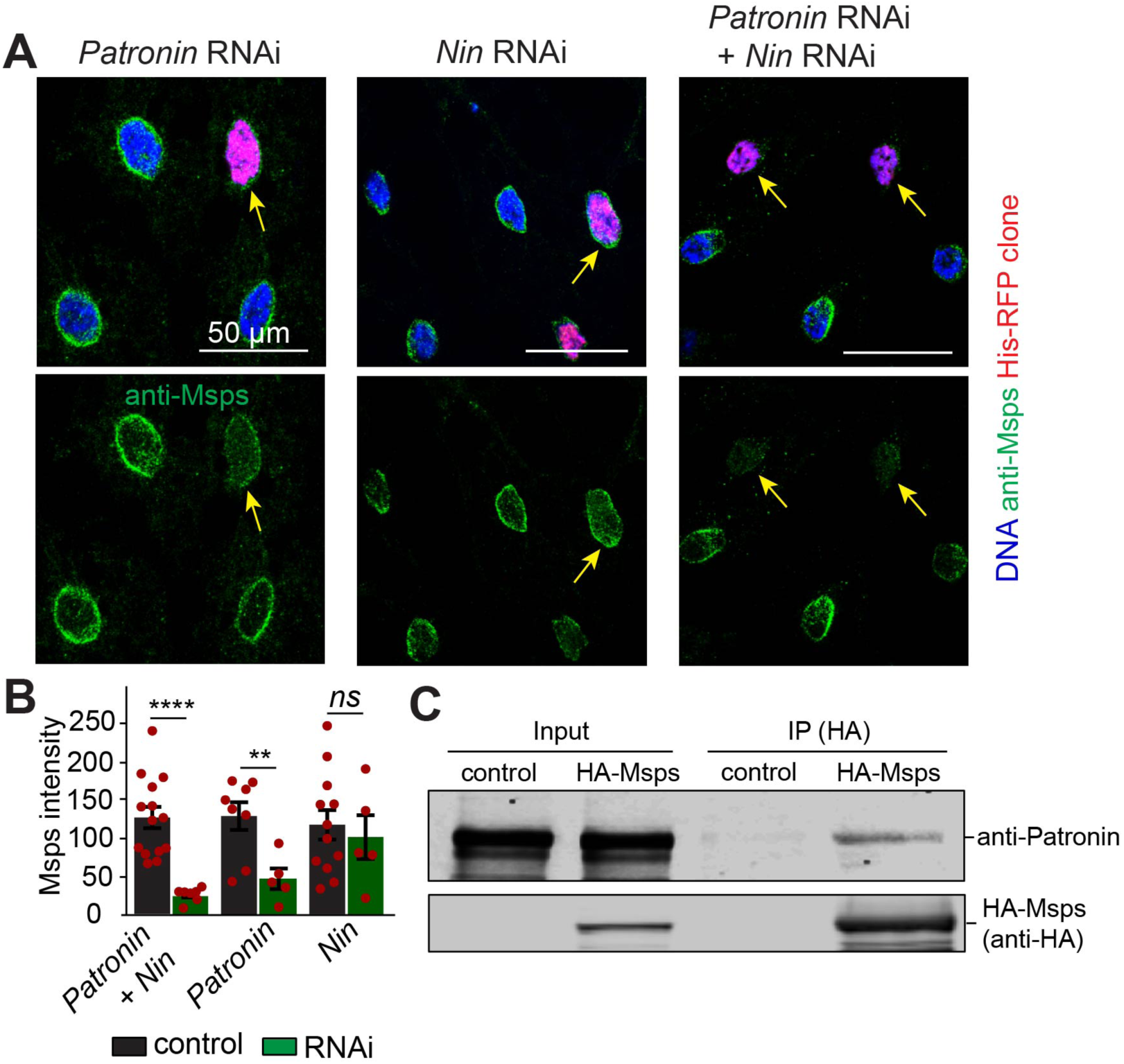
Patronin and Nin cooperate to recruit Msps to the ncMTOC. (**A**) Patronin has a significant role in recruiting Msps and Nin compensates when Patronin is depleted. Msps localization at the nuclear surface (arrows) in the indicated RNAi knockdown clones marked with His-RFP. Scale bar, 50 μm. (**B**) Quantification of Msps perinuclear signal shown in A. Data are shown as the means ± s.e.m. Statistics by two-tailed Student’s *t*-test. *Patronin* RNAi + *Nin* RNAi vs control, *p*<0.0001 (****), *Patronin* RNAi vs control, *p*=0.00867 (**), *Nin* RNAi vs control, *p*=0.6572 (*ns*). (**C**) HA-Msps (anti-HA) coimmunoprecipitation with endogenous Patronin (anti-Patronin) from *Drosophila* S2 cells. Input was 2.1% of total lysate. CoIP was performed independently twice.

These combined data point to a novel mechanism for ncMTOC function where Patronin, and perhaps also Nin, cooperate to recruit Msps to promote elongation of radial MTs, and do so independently of *γ*-tubulin (Figure 22).

**Figure 22.**
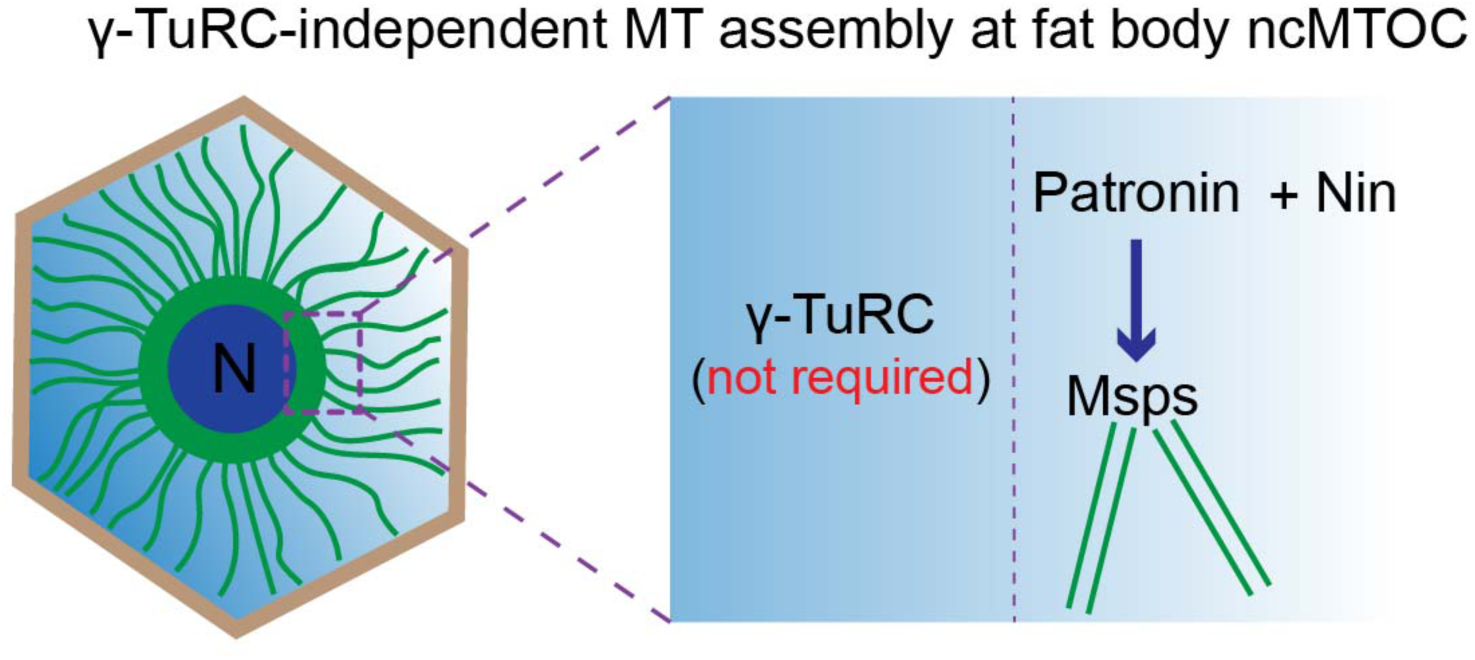
Working mechanism for MT assembly at the fat body ncMTOC. The fat body ncMTOC operates with a MT assembly mechanism distinct from the centrosome. The fat body ncMTOC does not require the widespread nucleator *γ*-TuRC. Instead, a unique MT assembly mechanism is proposed: Patronin and Nin, two MT minus-end proteins cooperate to stabilize and anchor circumferential MT seeds, and recruitment of the MT polymerase Msps elongates radial MTs. N: nucleus.

### The Nesprin Msp300 anchors the fat body ncMTOC at the nuclear surface and recruits Shot and Patronin

We investigated how the fat body ncMTOC is anchored to the nuclear surface and reasoned that a likely candidate is the Linker of Nucleoskeleton and Cytoskeleton (LINC) complex. The LINC complex connects cytoskeletal structures in the nucleoplasm with the cytoplasmic cytoskeleton via nuclear envelope-spanning proteins, which localize on the inner and outer nuclear envelopes and interact binding of their KASH and SUN domains within the inter-membrane space (Mejat and Misteli, 2010). KASH domain-containing Nesprins jut into the cytoplasm. *Drosophila* has two Nesprins: Klarsicht (Klar) and Msp300, both of which are localized to the nuclear envelope in fat body cells, along with the single SUN domain protein Klaroid (Koi) (Figure 23A). A null allele of *klar* or *koi* had little or moderate effect on nuclear centricity (Figure 8B, Table 1, and not shown). However, knockdown or mutation of *Msp300* dramatically disrupted nuclear centricity (Figure 23B, C) and MT assembly at the ncMTOC (Figure 23D, D’), demonstrating its requirement for a functional ncMTOC.

**Figure 23.**
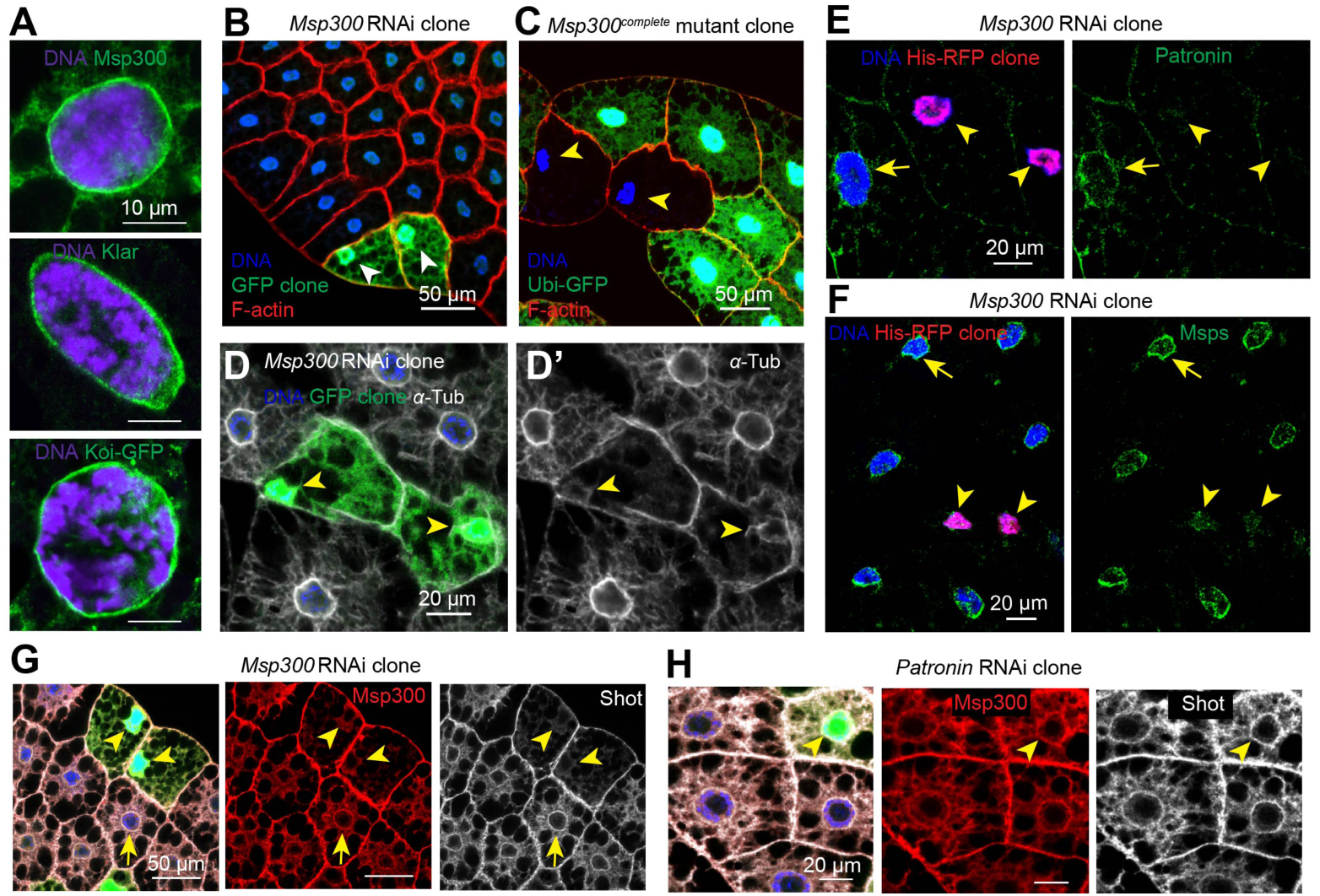
Msp300/Nesprin is required for MTOC assembly at the fat body nuclear surface by recruiting Shot, Patronin and Msps. (**A**) The LINC complex components Msp300, Klar and Koi are localized to the nuclear periphery in fat body cells. Note that Msp300 is perinuclear and also associated with cytoplasmic filaments, while Klar and Koi are perinuclear only. Scale bars: 10 μm. (**B, C**) Msp300 is required to maintain nuclear centricity (arrowheads) in fat body cells. *Msp300* RNAi knockdown clones are marked with GFP expression (**B**) and mutant clones are marked by loss of GFP expression (**C**). Quantification of nuclear centricity is shown in Figure 8B. (**D**) Fat body cell circumferential and radial MT arrays at the nuclear surface are diminished when *Msp300* is knocked down. Nuclei in RNAi knockdown clones are marked with GFP (arrowhead). Scale bar: 20 μm. (**E**) Patronin localization (anti-Patronin) to the nuclear surface is reduced (arrowhead for clone; arrow for control) in the *Msp300* RNAi knockdown clones marked with His-RFP. Quantification of Patronin perinuclear signal (not shown in the figure): 66.48 ± 10.21 (control, n=10) vs 2.983 ± 0.7863 (*Msp300* RNAi clone, n=9), *p*<0.0001 (****). Data are shown as means ± s.e.m. Scale bar: 20 μm. (**F**) Msps localization (anti-Msps) to the nuclear surface is reduced (arrowhead; arrow for control) in the *Msp300* RNAi knockdown clones marked with His-RFP. Quantification of Msps perinuclear intensity (not shown in the figure): 132.8 ± 11.13 (control, n=22) vs 30.5 ± 10.18 (*Msp300* RNAi clone, n=7), *p*<0.0001 (****). Data are shown as means ± s.e.m. Scale bar: 20 μm. (**G**) Shot (anti-Shot) is localized to the perinuclear MTOC (arrow) and its localization is disrupted in *Msp300* RNAi knockdown clones (arrowheads). Note that both perinuclear and cytoplasmic Msp300 are mostly eliminated by RNAi. Scale bar: 50 μm. (**H**) Shot and Msp300 localization to the perinuclear MTOC does not require Patronin as their localization appears normal in *Patronin* RNAi knockdown clone (arrowhead) by antibody staining. Scale bar: 20 µm.

Consistent with its role in controlling assembly of the ncMTOC, the localization of Patronin and Msps at the nuclear surface were blocked when *Msp300* was knocked down (Figure 23E, F). These results indicate that Msp300 organizes or anchors the ncMTOC on the nuclear surface, and we propose it does so by recruiting the key MT regulator Patronin.

We further showed that Short stop (Shot), the sole spectraplakin in *Drosophila* (Roper et al., 2002), was localized to the nuclear surface and was dependent on Msp300 for localization (Figure 23G). Shot was shown to associate with Patronin and to be required to localize it to ncMTOCs at the plasma membrane of the anterior oocyte and the apical surface of follicle cells (Khanal et al., 2016; Nashchekin et al., 2016). Knockdown of *Patronin* did not impact Msp300 or Shot localization (Figure 23H), consistent with a dependence of Patronin on Msp300 and/or Shot for its localization at the MTOC. These combined analyses suggest that Msp300 is the primary organizer of the perinuclear ncMTOC by recruiting Shot and Patronin. Patronin, together with Nin, generates circumferential MTs, and they recruit Msps to assemble radial MTs (see Figure 27).

### Shot depletion shifts the perinuclear ncMTOC to an ectopic MTOC

While Shot localization to the fat body perinuclear ncMTOC depends on Msp300 (Figure 23G), Msp300 localization to the nuclear periphery also depends on Shot; Msp300 was delocalized from the nuclear surface to the center of an ectopic cytoplasmic MTOC upon *shot* knockdown (Figure 24A, B). In contrast to *Msp300* knockdown, knockdown of *shot* resulted in a more severe phenotype: a remodeling of the perinuclear ncMTOC into a centrosome-like MTOC focus in the cytoplasm (Figure 24A). Besides Msp300, additional components of the fat body ncMTOC were also delocalized to the ectopic MTOC by *shot* knockdown, including Patronin and Msps (Figure 24B), whereas proteins not required for MTOC function such as Cnn and *γ*-tubulin were not (Figure 24C-E).

**Figure 24.**
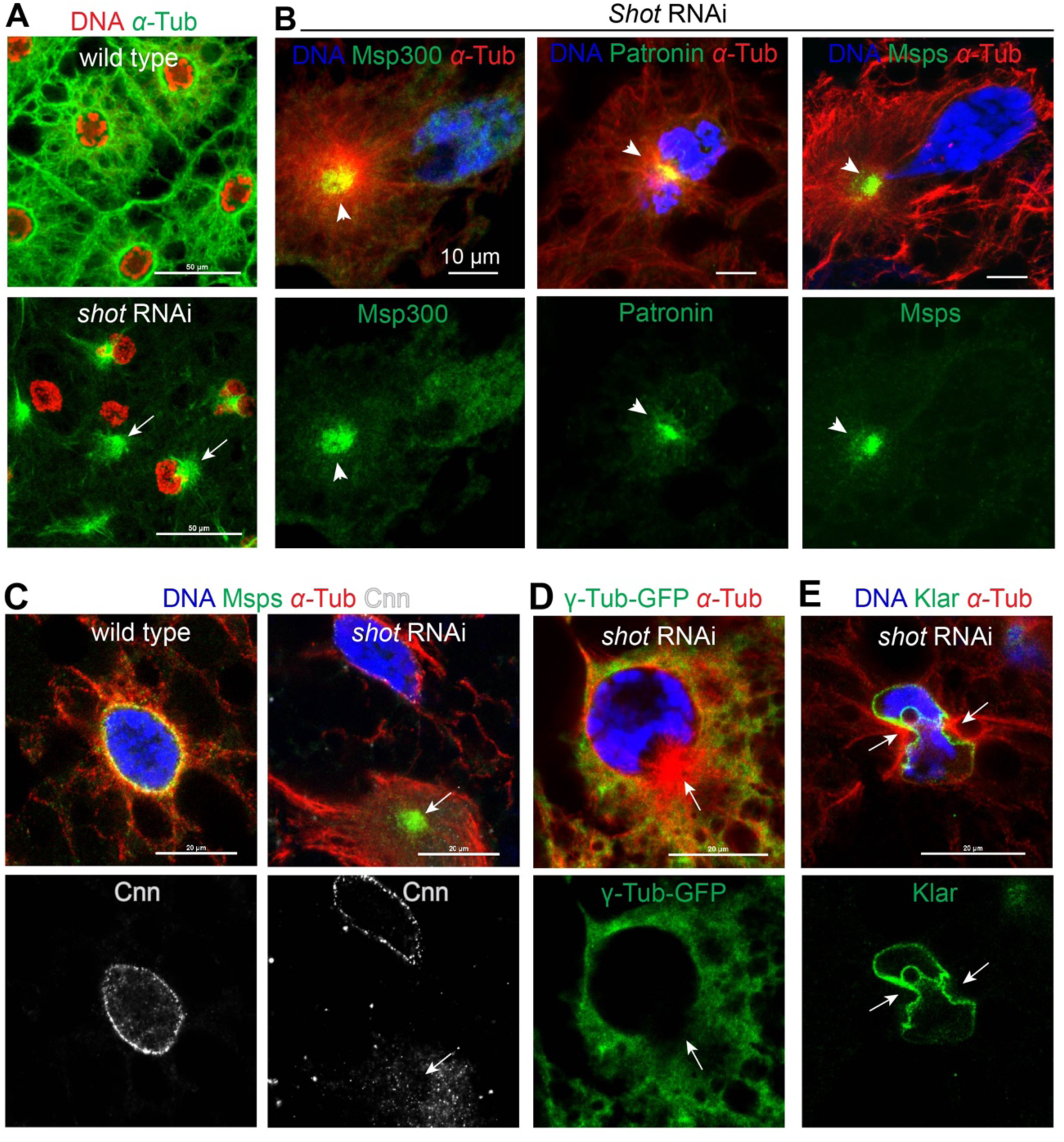
Shot depletion generates an ectopic centrosome-like MTOC that is enriched in Msp300, Patronin and Msps. (**A**) *shot* knockdown results in a reorganization of the MTOC from the nuclear surface to a centrosome-like focus in the cytoplasm (arrows). Scale bars: 50 μm. (**B**) Msp300, Patronin and Msps are delocalized to the ectopic MTOC (arrowheads) in *shot* knockdown fat body cells by antibody staining. Scale bars: 10 μm. (**C, D, E**) Cnn (**C**), *γ*Tub23C (**D**) and Klar (**E**) are not delocalized from the nuclear surface to the ectopic MTOC (arrows) generated by *shot* knockdown. Arrow in (**D**) indicates no *γ*Tub23CGFP enrichment in the ectopic centrosome-like MTOC. Scale bar: 20 μm.

Reasoning that Msp300 could be the primary regulator of the *shot* RNAi-induced ectopic MTOC, we co-depleted *Msp300* and *shot* in fat body cells. Further knockdown of *Msp300* in *shot* RNAi fat body significantly diminished the MT density of the centrosome-like foci (Figure 25), and blocked Patronin recruitment there (Figure 26). Co-depletion of *shot* with *Patronin* or *msps* also reduced the ectopic MT density and abrogated the formation of centrosome-like MTOC (Figure 25), further indicating that Msp300, Patronin and Msps are shared key MT regulators for the normal perinuclear MTOC and also for the ectopic MTOC that is induced by *shot* depletion. In addition to MT density change, it appears that *Patronin* double knockdown with *shot* also altered the ectopic MT foci morphology. Knockdown of *Patronin* and *shot* together caused a more disperse MT pattern of the residual MTOC compared to the concentrated MT focus in *shot* knockdown alone. Taken together, these data indicate that Shot and Msp300 are co-dependent for their assembly into the perinuclear ncMTOC, and the generation of an ectopic MTOC (that contains Patronin and Msps) after *shot* depletion indicates that Msp300 recruits Patronin and Msps independently of Shot (Figure 27).

**Figure 25.**
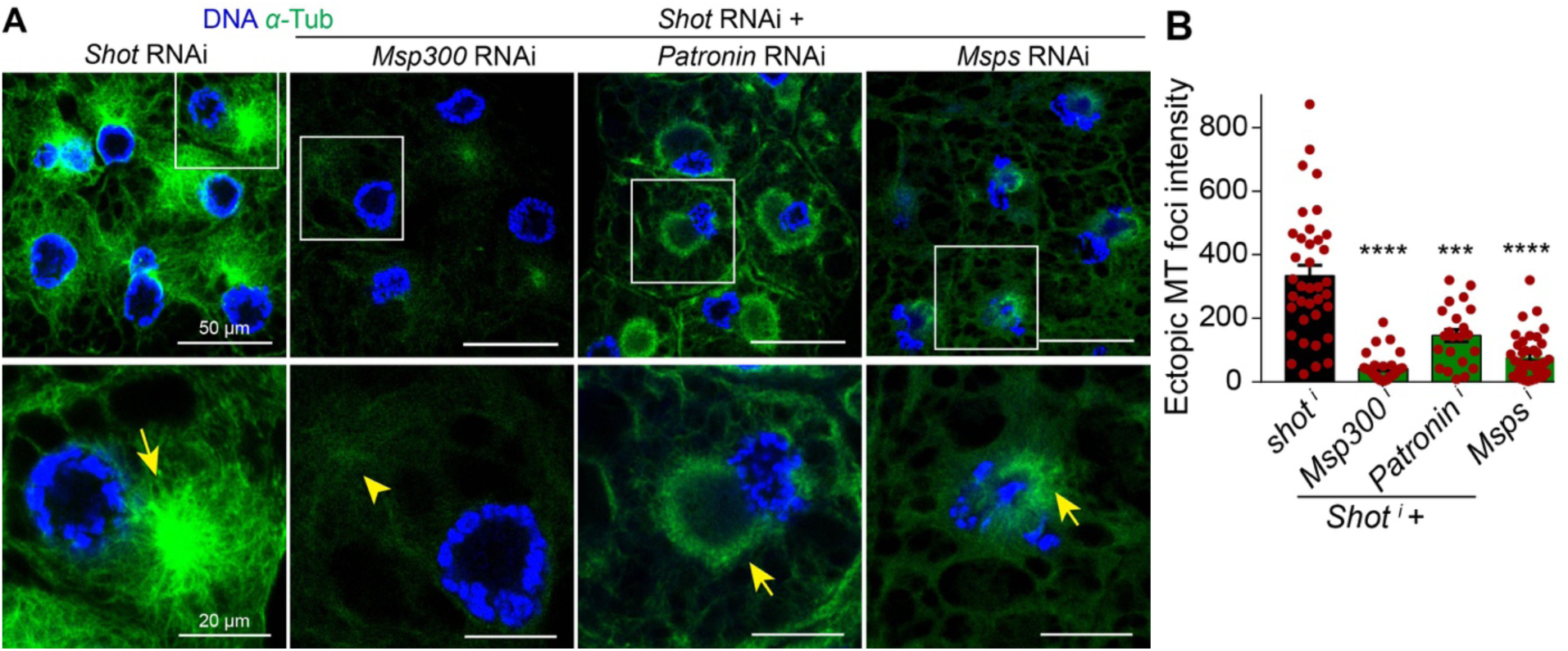
Additional depletion of Msp300, Patronin or Msps attenuates Shot RNAi-induced ectopic MTOC. (**A**) MT staining showing that *shot* RNAi generated a robust MT focus (arrow), and additional RNAi knockdown of *Msp300* by two fat body drivers (*SPARC-Gal4* + *Cg-Gal4*) in the *shot* RNAi background dramatically diminished cytoplasmic MT foci (arrowhead). *Patronin* and *shot* double knockdown by two fat body drivers (*SPARC-Gal4* + *Cg-Gal4*) reduced cytoplasmic MT foci, and also formed more disperse MTs around nucleus. *msps* and *shot* double knockdown by *SPARC-Gal4* attenuated cytoplasmic MT foci. Scale bars: 50 µm (top panel), 20 µm (bottom panel). (**B**) Quantification of ectopic MT foci shown in **A**.*shot* RNAi vs *shot* RNAi + *Msp300* RNAi, *p*<0.0001 (****), *shot* RNAi vs *shot* RNAi + *Patronin* RNAi, *p*=0.0001 (***), *shot* RNAi vs *shot* RNAi + *msps* RNAi, *p*<0.0001 (****). Data are shown as the means ± s.e.m. Statistics by two-tailed Student’s *t*-test.

**Figure 26.**
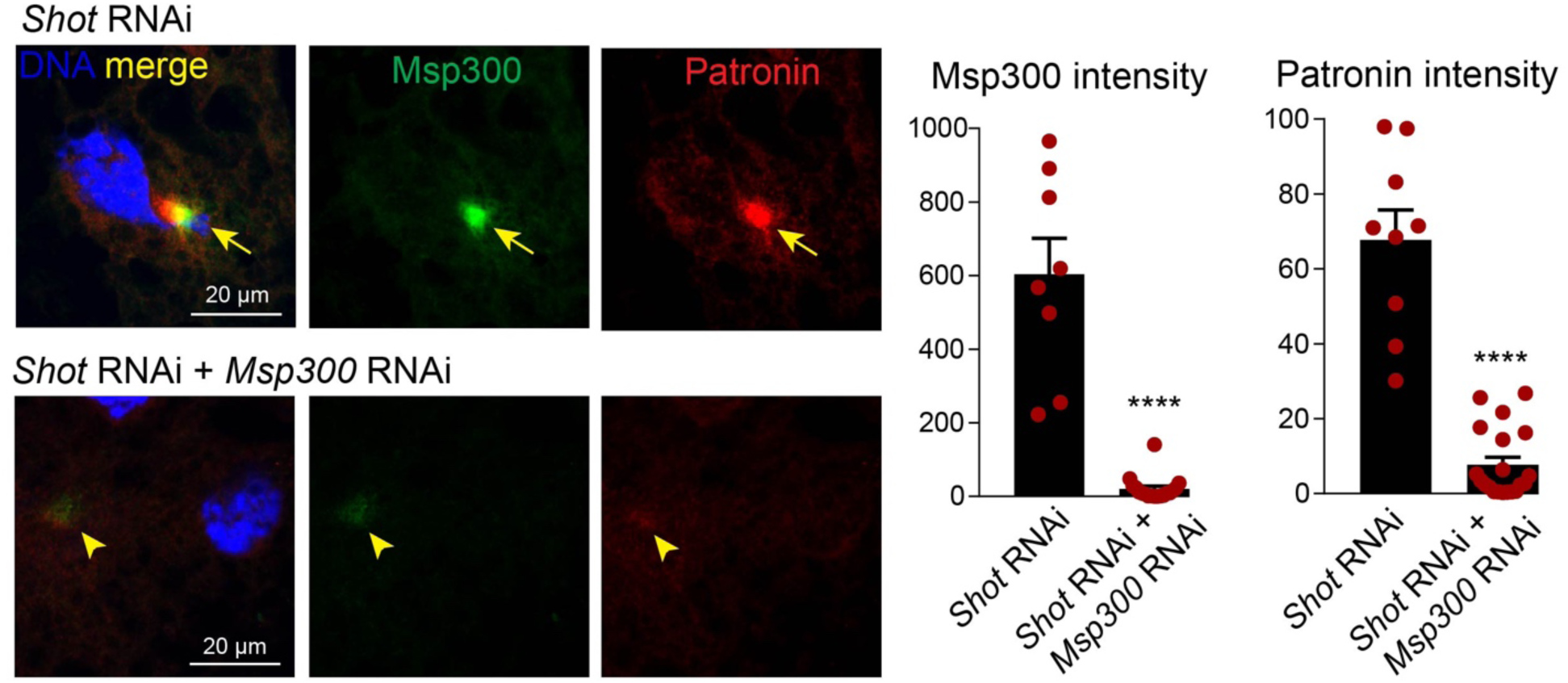
Msp300 could recruit Patronin independently of Shot. *Shot* RNAi delocalized Msp300 and Patronin (arrows), and additional *Msp300* RNAi in the background of *shot* RNAi efficiently knockdown Msp300 (arrowhead), and this also reduced Patronin recruitment to cytoplasmic foci (arrowhead). *Shot* RNAi vs *Shot* RNAi + *Msp300* RNAi, *p*<0.0001 (****) for Msp300 and Patronin foci intensity. Data are shown as the means ± s.e.m. Statistics by twotailed Student’s *t*-test. Scale bars: 20 μm.

**Figure 27.**
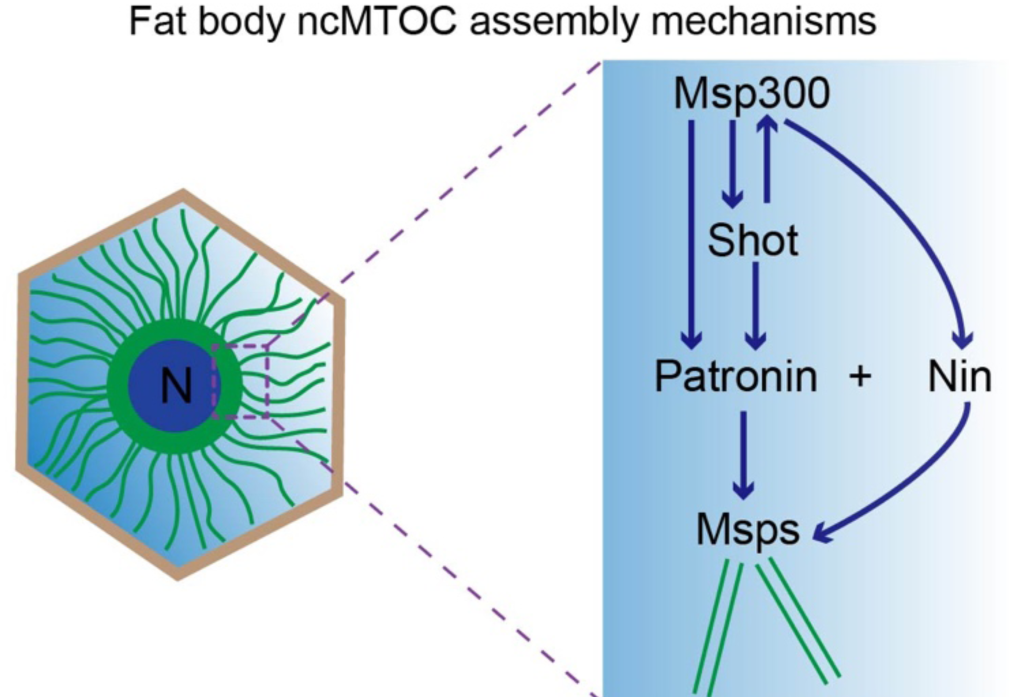
Assembly order/epistasis among Msp300, Shot, Patronin, Nin and Msps. Msp300 anchors ncMTOC to the nuclear surface and recruits Shot and Patronin. Msp300 and Shot are co-dependent for perinuclear localization. Msp300 could recruit Patronin independently of Shot.

Shot disruption generates additional pleiotropic phenotypes besides perturbing the perinuclear MTOC. Knockdown of *shot* also reorganized cortically enriched filamentous actin into large aggregates that accumulate in the proximity of the displaced MTOC (Figure 28A); deformed the nucleus (Figure 28B); reorganized many subcellular compartments including ER, Golgi, mitochondria, and endosomes (Figure 28C and not shown); and blocked secretion of basement membrane components (Figure 28D, E, E’), which abnormally accumulated in proximity to the collapsed MT network (Figure 28D). These severe disruptions involve more than Shot’s role at the ncMTOC, because knockdown of Msp300 disrupted Shot localization to the ncMTOC but did not result in the severe and pleiotropic effects seen with *shot* knockdown.

**Figure 28.**
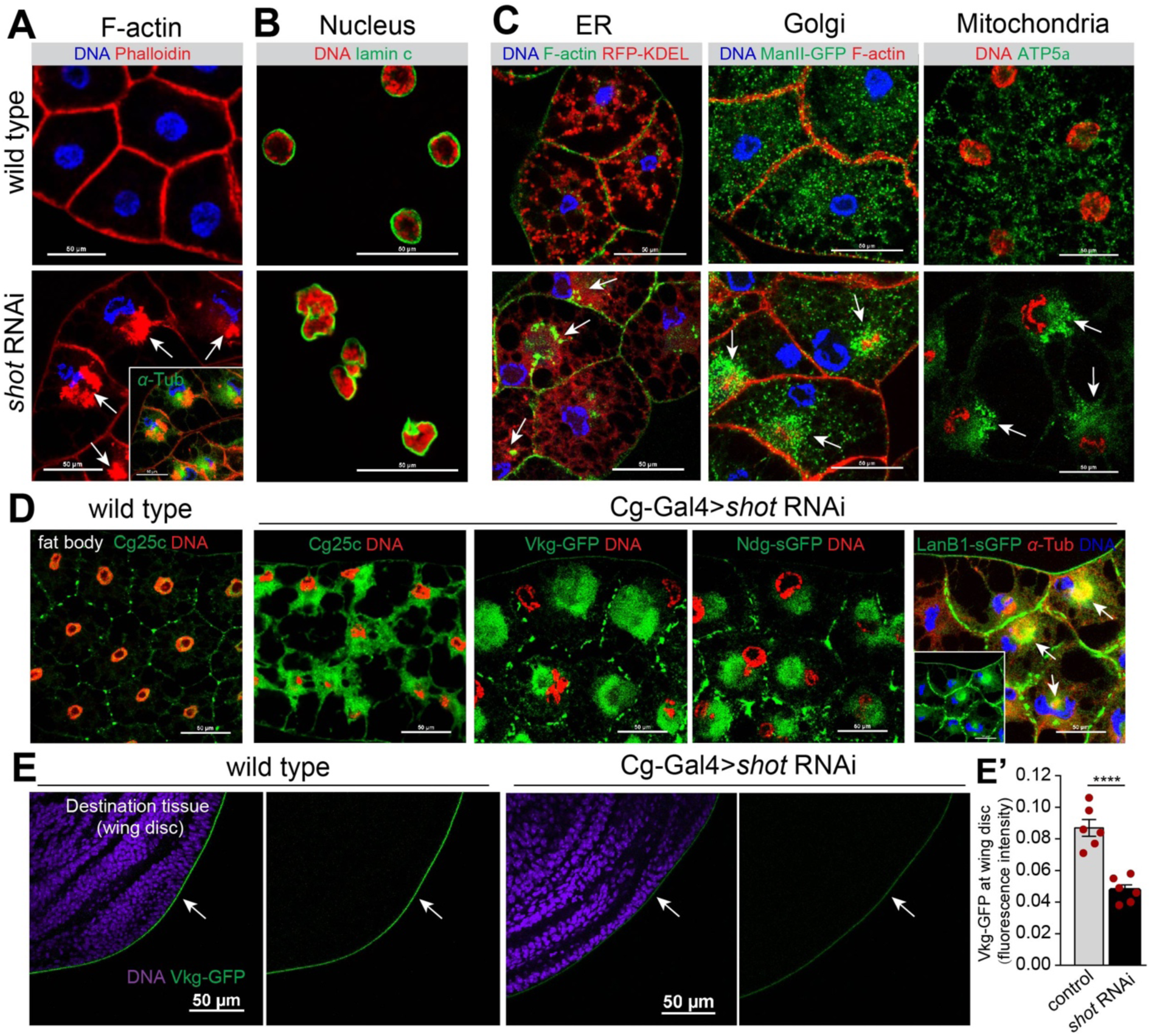
Shot depletion results in severe and pleiotropic disruption of subcellular compartments. (**A**) Actin collapses into an aggregate (arrows) near the MTOC after *shot* knockdown. Note that nuclear centricity is also affected. Inset shows the merged image with MTs. (**B, C**) Nuclear morphology (**B**) and organization of organelles (**C**) are disrupted in *shot* knockdown fat bodies. Arrows point to accumulation of ER, Golgi and mitochondria near actin aggregates. (**D**) *shot* knockdown results in accumulation of basement membrane components in the cytoplasm, proximal to the ectopic MTOC (arrows). (**E**) Deposition of collagen IV a2 (Vkg-GFP, arrows) on the wing disc is decreased when *shot* is knocked down in fat body cells. (**E’**) Quantitation of Vkg-GFP in (**E**). Data are shown as means ± s.e.m (n=6 wing discs). *P*<0.0001 (****) by two-tailed Student’s *t*-test. Scale bars: 50 μm (**A-E**).

### The fat body ncMTOC is essential for basement membrane (BM) secretion

The fat body is the major source of synthesis and secretion of the collagen IV trimer (Pastor-Pareja and Xu, 2011), whose subunits are encoded by *Collagen IV α1* (*Cg25c)* and *Collagen IV α2* (*Viking* (*vkg*)) (Figure 29A). A small fraction of secreted collagen IV is incorporated into fat body cell junctions as collagen IV intercellular concentrations (CIVICs) (Dai et al., 2017), but the majority is secreted, transported via the hemolymph, and deposited along with other BM components, like Perlecan (Trol), LanB1 and Nidogen (Ndg) at destination organs like imaginal discs and the brain (Figure 29A) (Pastor-Pareja and Xu, 2011).

**Figure 29.**
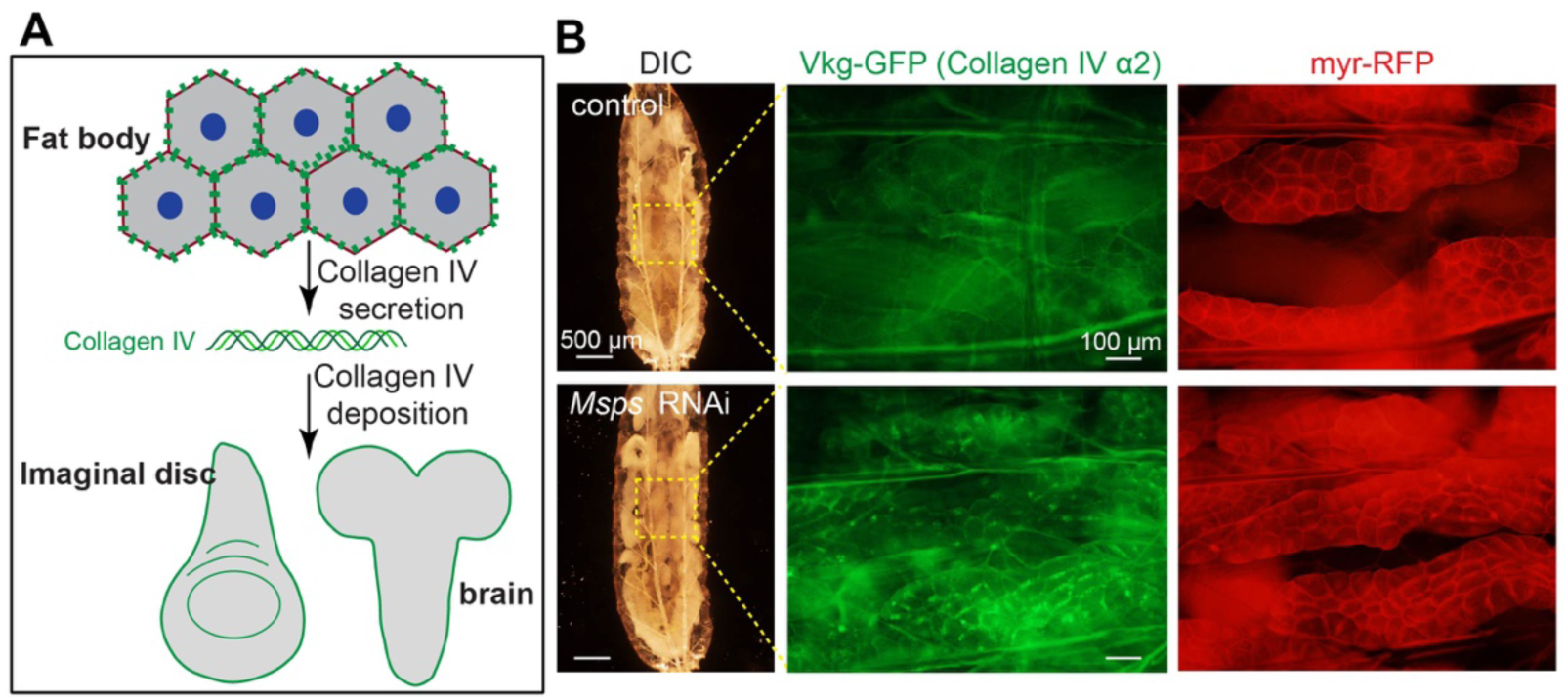
The fat body ncMTOC is essential for secretion of basement membrane proteins. (**A**) Diagram of basement membrane component collagen IV production in the fat body and its secretion and deposition at destination tissues as basement membrane. (**B**) DIC and fluorescent images of collagen IV a2 (Vkg-GFP) in live fat bodies labelled with the membrane marker myr-RFP from control larvae and larvae with fat body-specific knockdown of *Msps*. Scale bars: 500 μm (DIC images), 100 μm (fluorescent images).

When fat body MTs were disrupted or if the MTOC was impaired, collagen IV and other BM proteins accumulated to significantly higher levels at the plasma membrane of fat body cells and were visible in intact larvae (Figure 29B) or at dissected fat bodies (Figure 30A, B). Disruption of actin, on the other hand, did not result in accumulation of BM proteins in fat body (Figure 30A). As a consequence of fat body MT disruption and the accumulation of BM proteins at the plasma membrane, deposition of BM components at destination organs like the wing imaginal disc and brain were significantly reduced (Figure 31). Depletion of *γ*-TuRC components did not cause accumulation of BM components on the plasma membrane (Figure 30B), in accord with no involvement of the *γ*-TuRC in fat body ncMTOC assembly.

**Figure 30.**
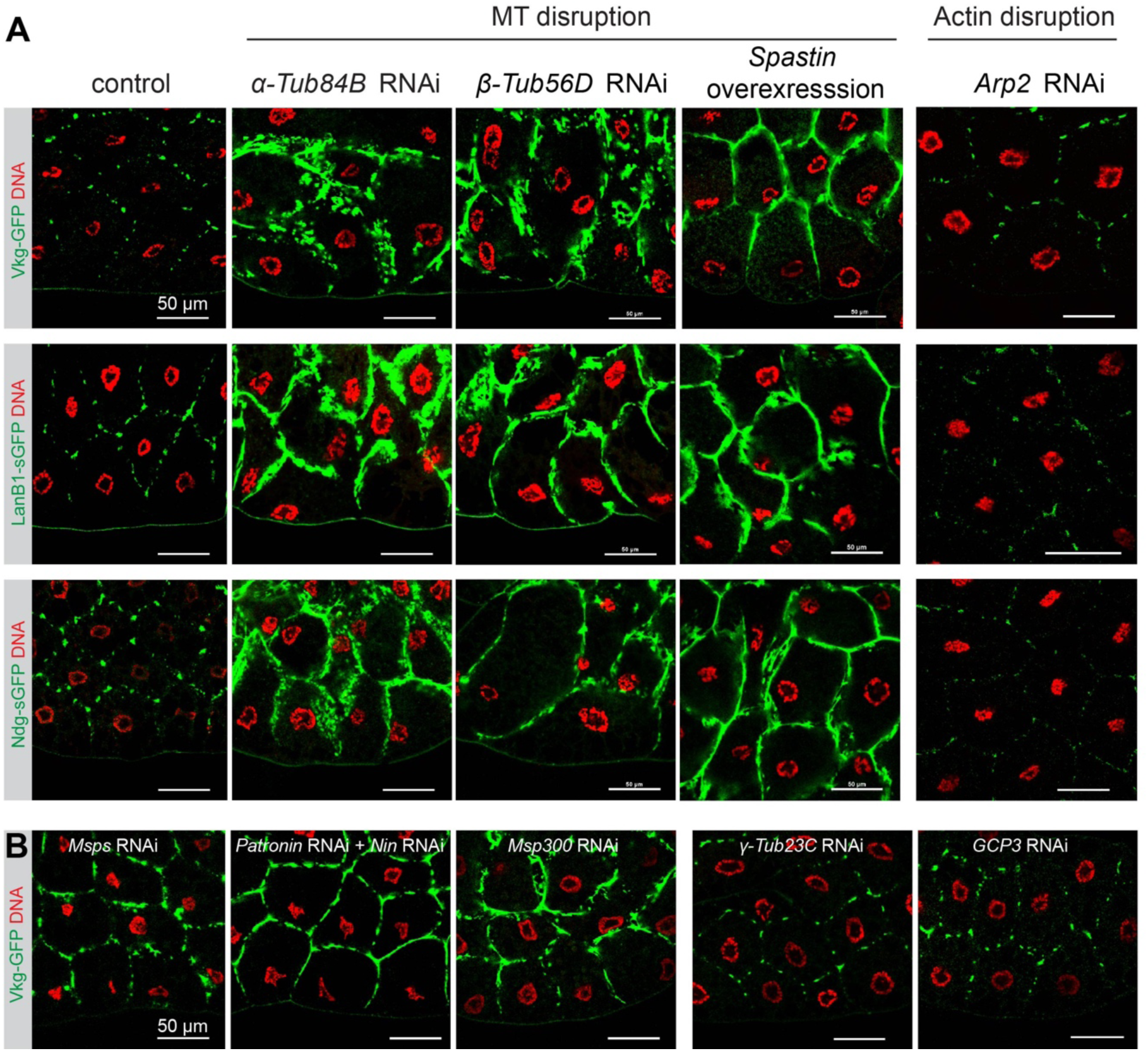
The fat body ncMTOC, but not actin is essential for secretion of basement membrane proteins. (**A**) Images showing fat body accumulation of indicated basement membrane components after the indicated MT or actin disruption in fat body cells. Vkg-GFP (collagen IV α2) is shown by GFP autofluorescence. LanB1-sGFP and Ndg-sGFP are stained with anti-GFP antibody. (**B**) Images showing fat body accumulation of Vkg-GFP after the indicated RNAi. Scale bars: 50 μm.

**Figure 31.**
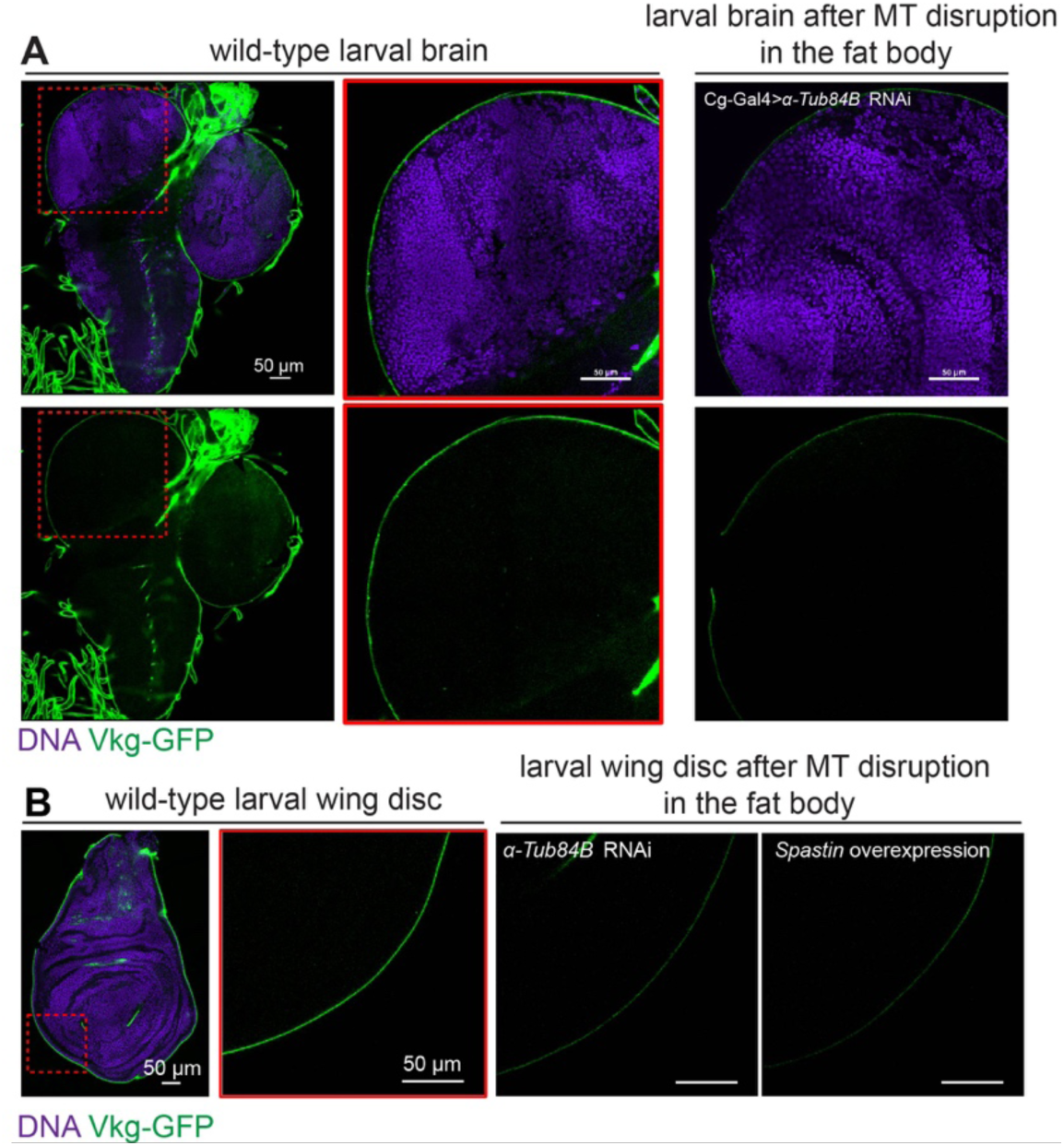
The fat body ncMTOC is essential for fat body secretion of basement membrane proteins to distant organs. (**A**) Images showing reduced brain deposition of Vkg-GFP after MT disruption in fat body cells by α*-Tub84B* RNAi. (**B**) Images showing reduced wing disc deposition of Vkg-GFP after MT disruption in fat body cells by α*-Tub84B* RNAi or *Spastin* overexpression. Scale bars: 50 μm.

### The fat body ncMTOC is essential to restrict plasma membrane growth and avert extracellular entrapment of basement membrane components

The accumulation of BM proteins at the plasma membrane following MT disruption is extracellular or pericellular, but not intracellular, as demonstrated by antibody binding to Vkg-GFP on fixed but non-permeabilized fat body cells (Figure 32). Thus, BM proteins are secreted but trapped outside of the plasma membrane. The extracellular entrapment of collagen indicates that MTs are not required for anterograde collagen trafficking or secretion *per se* across the plasma membrane, but somehow are responsible for their retention there.

**Figure 32.**
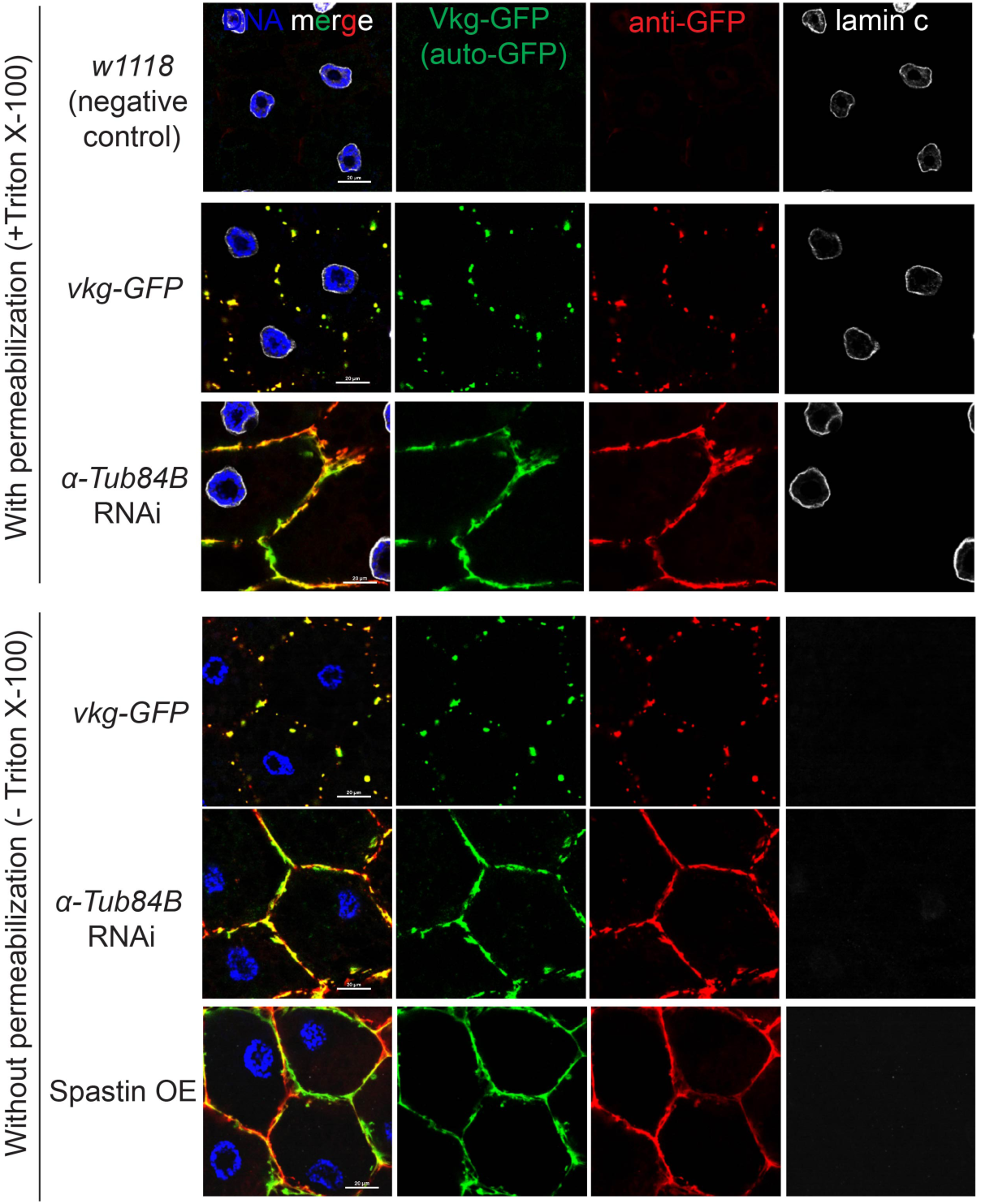
The accumulation of Vkg-GFP after MT disruption is extracellular, not intracellular. Images showing that the Vkg-GFP accumulation at the plasma membrane is extracellular in α*-Tub84B* RNAi fat body cells by immunostaining with or without detergent permeabilization. Vkg-GFP autofluorescence is shown green and GFP antibody staining is labeled red. Lamin C staining is a control for cell permeabilization. Scale bars: 20 μm.

This phenotype of extracellular BM entrapment was seen in fat body cells when endocytosis was disrupted (Zang et al., 2015). The balance of plasma membrane levels is controlled by endocytic membrane trafficking. In the absence of endocytosis, the plasma membrane overgrows, becomes highly convoluted, and results in extracellular entrapment of secreted collagen within the folds of excess plasma membrane (Zang et al., 2015). Disruption of the fat body MTOC caused similar overgrowth of the plasma membrane as revealed by the membrane-targeted marker myr-RFP (Figure 33A, B) or with the membrane dye CellMask (Figure 34), similar to knockdown of the endocytosis machinery from expression of dominant negative Rab5^S43N^ or *shi* (dynamin) knockdown (Figure 34A) (Zang et al., 2015). Knockdown of *Msp300* or *msps*, but not *γTub23C*, resulted in plasma membrane overgrowth and entrapment of collagen IV (Figure 33B).

**Figure 33.**
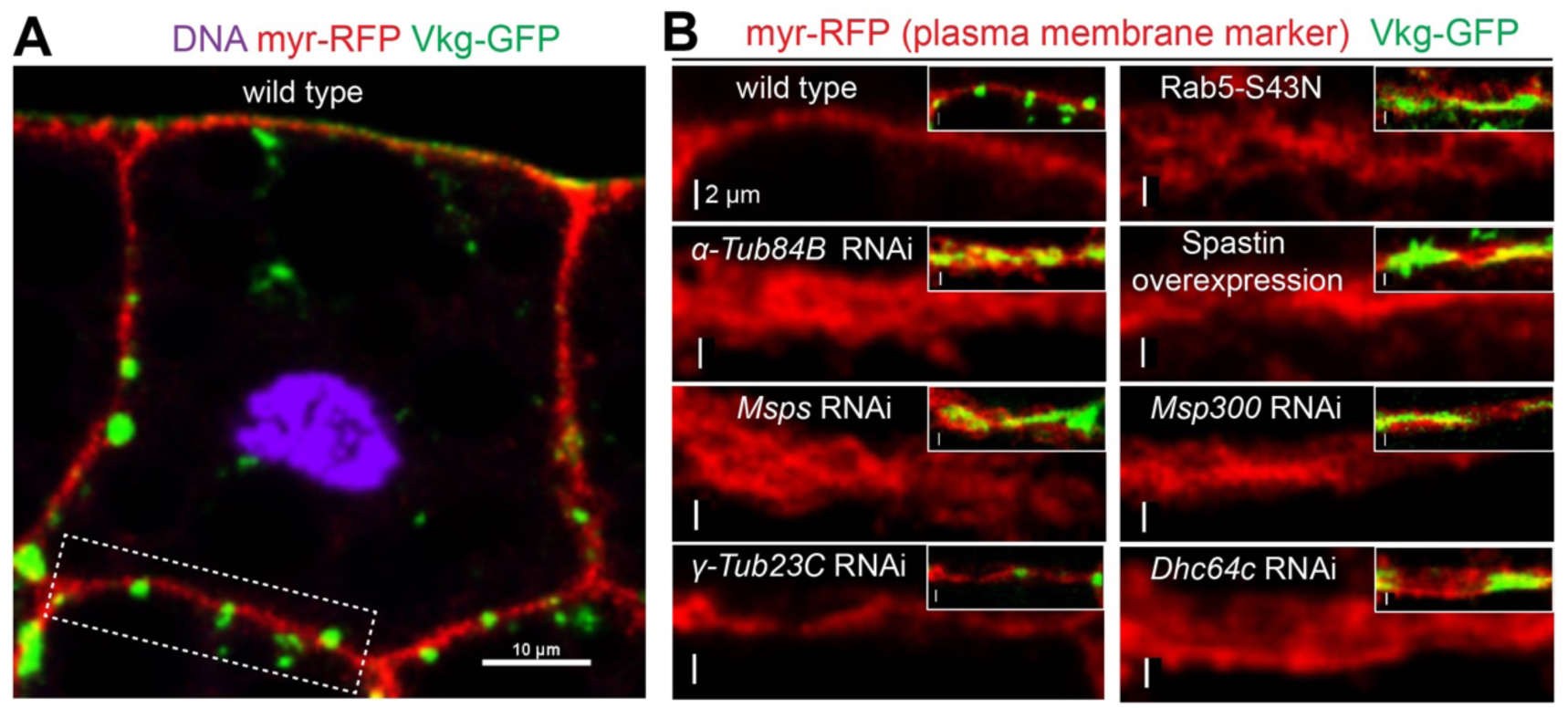
Fat body MT disruption causes “thickening” of the plasma membrane. (**A**) A fat body cell expressing Vkg-GFP and the plasma membrane marker myr-RFP. Scale bar, 10 μm. (**B**) Images showing that the plasma membrane marker myr-RFP becomes thicker due to plasma membrane overgrowth in fat body cells after disruption of MTs directly (α*-Tub84B* RNAi, *Spastin* overexpression) or by impairing the ncMTOC (*Msps* RNAi, *Msp300* RNAi), inactivation of dynein motor (*Dhc64c* RNAi) or endocytic machinery (Rab5-S43N, dominant negative Rab5, a positive control). Insets show merged images with Vkg-GFP at the plasma membrane. Scale bars: 2 μm.

**Figure 34.**
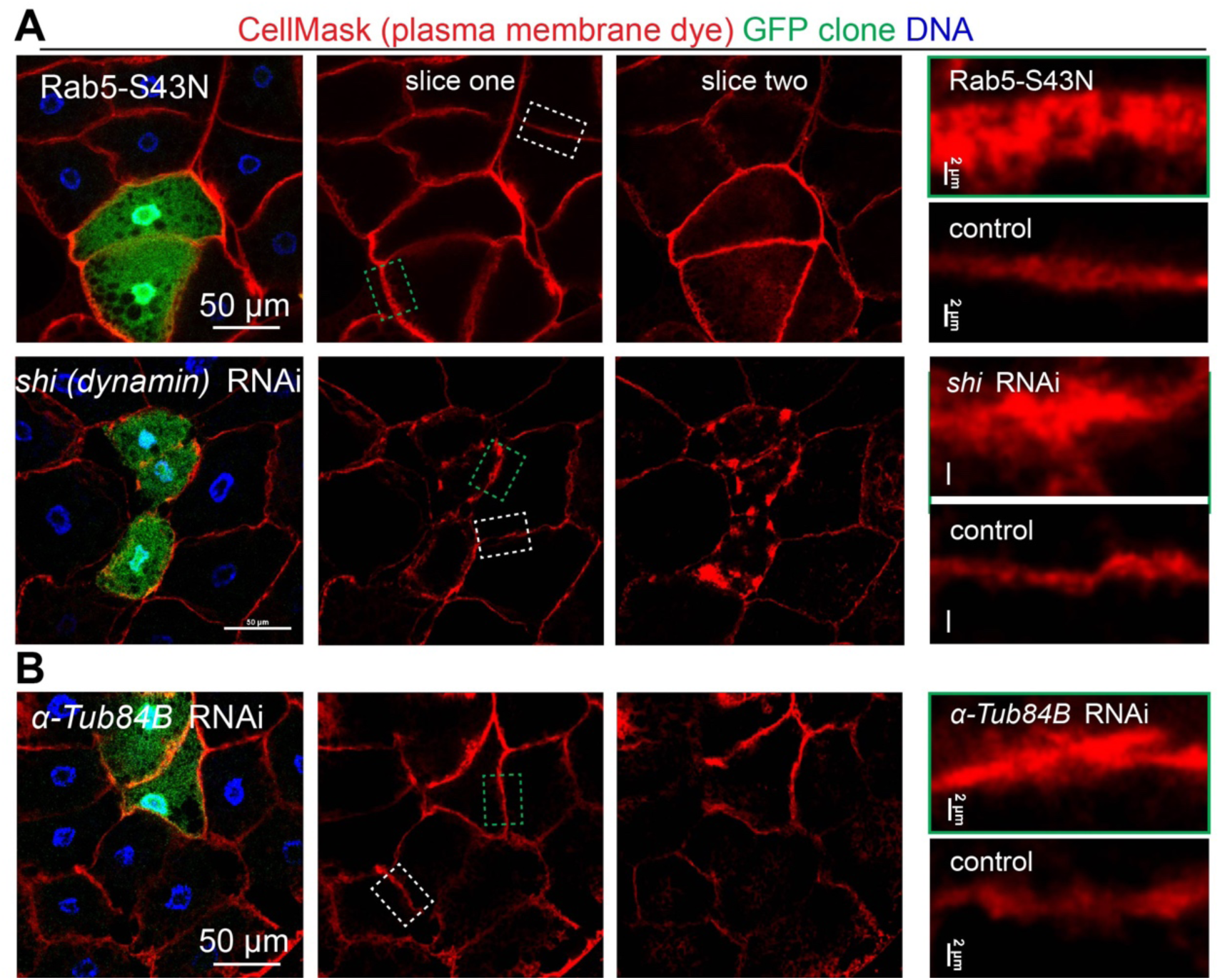
Fat body MT disruption and endocytosis blockage cause similar “thickening” of plasma membrane. (**A, B**) Images showing plasma membrane overgrowth using the membrane dye CellMask in GFP-marked fat body cells with disrupted endocytosis by overexpressing dominant negative Rab5 (Rab5-S43N) or knockdown of *shi* (**A**) or with disrupted MTs by knockdown α*-Tub84B* (**B**). Slices one and two are separate image sections from the same confocal image z-stack. The dashed boxes for control (white) and GFP+ clone (green) plasma membrane are shown in the panels on the right.

### The fat body ncMTOC supports retrograde dynein-dependent endosome trafficking to maintain proper plasma membrane growth

We tested the idea that the thickened plasma membrane arises due to a block in the trafficking of endocytic vesicles away from the plasma membrane when the fat body ncMTOC is impaired. Using GFP-Rab5 as an endosomal vesicle marker, we showed that in normal fat body cells GFP-Rab5 vesicles were enriched at the plasma membrane and also encircled the nucleus where the ncMTOC resides (Figure 35A). In sharp contrast, following disruption of the fat body ncMTOC, the perinuclear pool of GFP-Rab5 vesicles was diminished and instead became more highly enriched at the plasma membrane (Figure 35A). Thus, fat body MT arrays are required for retrograde membrane trafficking of endocytic vesicles from the plasma membrane to perinuclear sites (Figure 35B).

**Figure 35.**
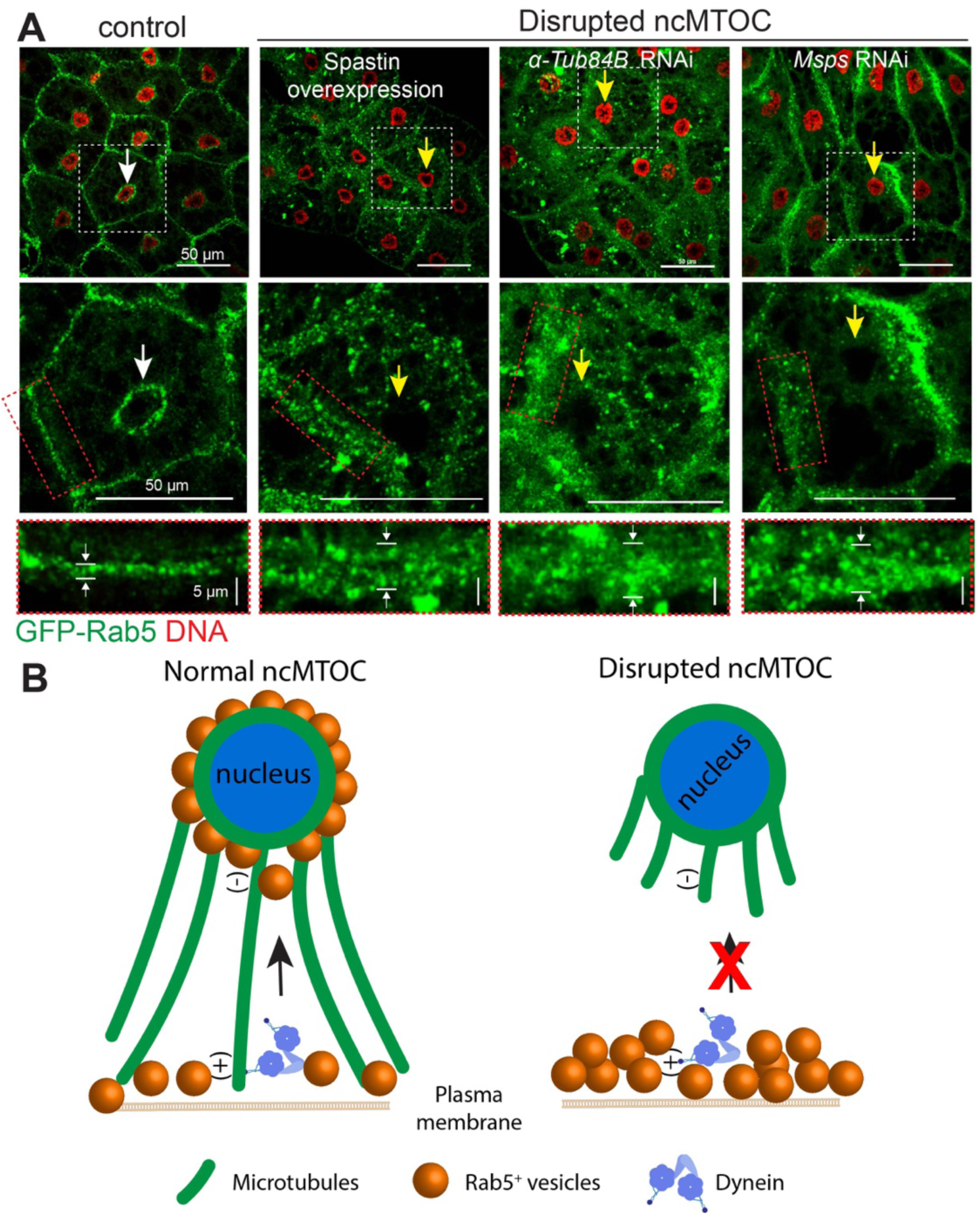
The fat body ncMTOC is essential to restrict plasma membrane growth. (**A**) GFP-Rab5 is normally distributed at two main sites: the plasma membrane and nuclear surface, but shifts to localizing predominantly near the plasma membrane after ncMTOC disruption. The arrows indicate the perinuclear pool of Rab5 and also the nuclear positioning (defective after disrupted ncMTOC). Scale bars: 50 μm top and middle, 5 μm bottom panel. (**B**) Illustration showing that disruption of the ncMTOC blocks retrograde membrane trafficking, leading to accumulation of endosomes at the plasma membrane.

Retrograde trafficking of endosomes requires the minus-end directed MT motor dynein in other systems (Granger et al., 2014). We tested the requirement for dynein in GFP-Rab5 distribution between the plasma membrane and the perinuclear sites in fat body cells. Loss of retrograde motor function, either by dynein RNAi or overexpression of dynamitin to inhibit dynein activity, shifted the perinuclear pool of GFP-Rab5 to the plasma membrane (Figure 36A, C). As a control, *shi* knockdown (disrupts endocytic budding) caused loss of plasma membrane GFP-Rab5 (Figure 37B, C). The blockage of endosomal trafficking by dynein inhibition phenocopied the effect of MT disruption on plasma membrane overgrowth (Figure 33B) and entrapment of collagen at the plasma membrane (Figure 37A), but not nuclear mispositioning (Figure 37B), consistent with dynein being required for trafficking upon MTs but not the organization of the MT array. When the Kinesin-1 motor was knocked down to block anterograde MT trafficking, there was no effect on BM secretion (Figure 37A) or nuclear positioning (Figure 37B). This indicates that a primary function for the fat body MTOC is to support retrograde endosomal trafficking by dynein.

**Figure 36.**
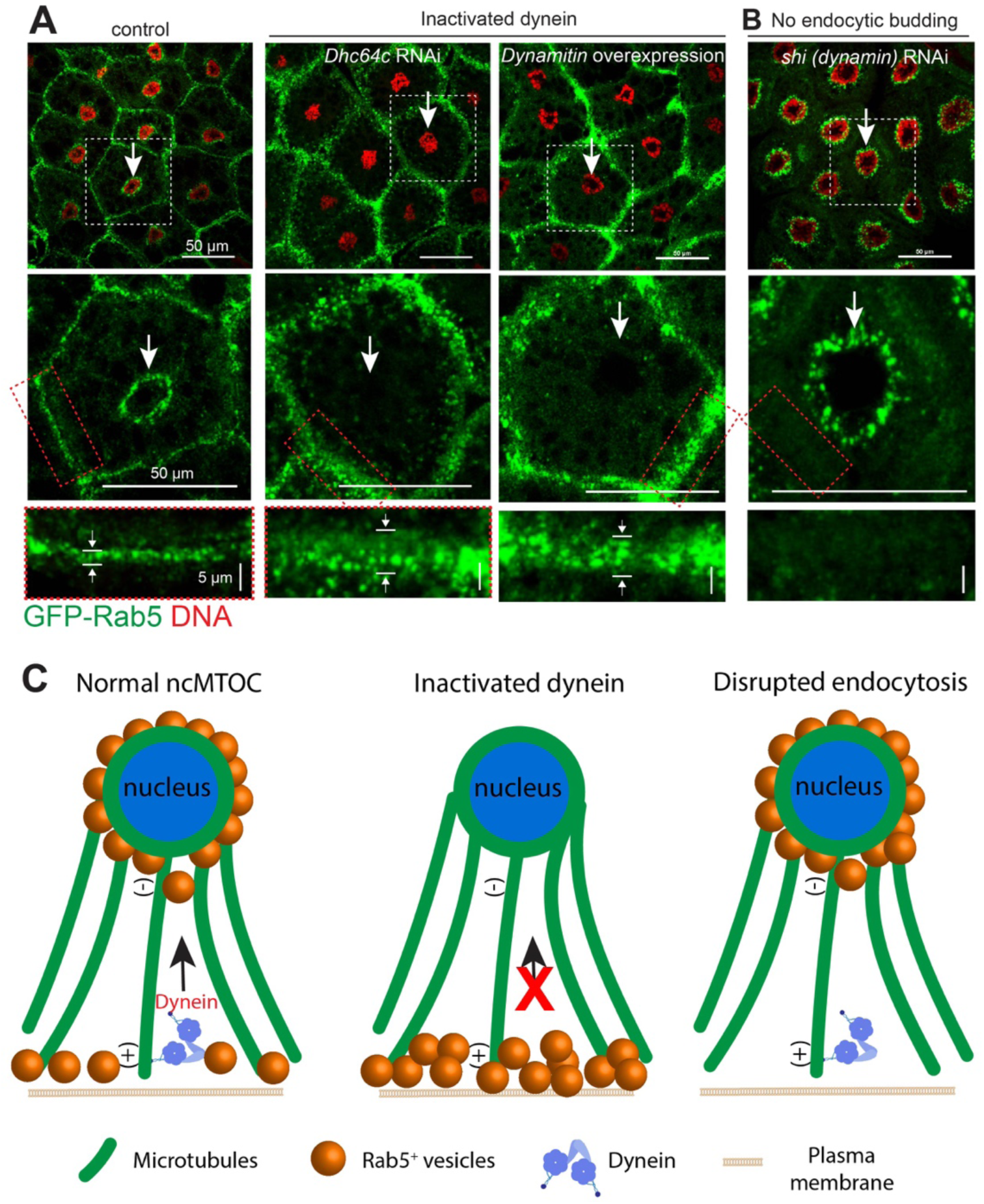
The dynein motor coordinates with the fat body ncMTOC for retrograde endosome trafficking to restrict plasma membrane growth. (**A**) Inactivation of dynein either by knockdown of *dynein heavy chain64c* (*Dhc64c*) or overexpression of inhibitory dynamitin caused Rab5 shifting from perinuclear site to plasma membrane (white arrows). Note nuclear positioning is normal. The control is reproduced from Figure 35A. (**B**) Blocking endocytosis machinery by *shi* (dynamin) knockdown caused loss of plasma membrane accumulation of Rab5. (**C**) Cartoon showing Rab5 localization after dynein inactivation or endocytosis disruption.

**Figure 37.**
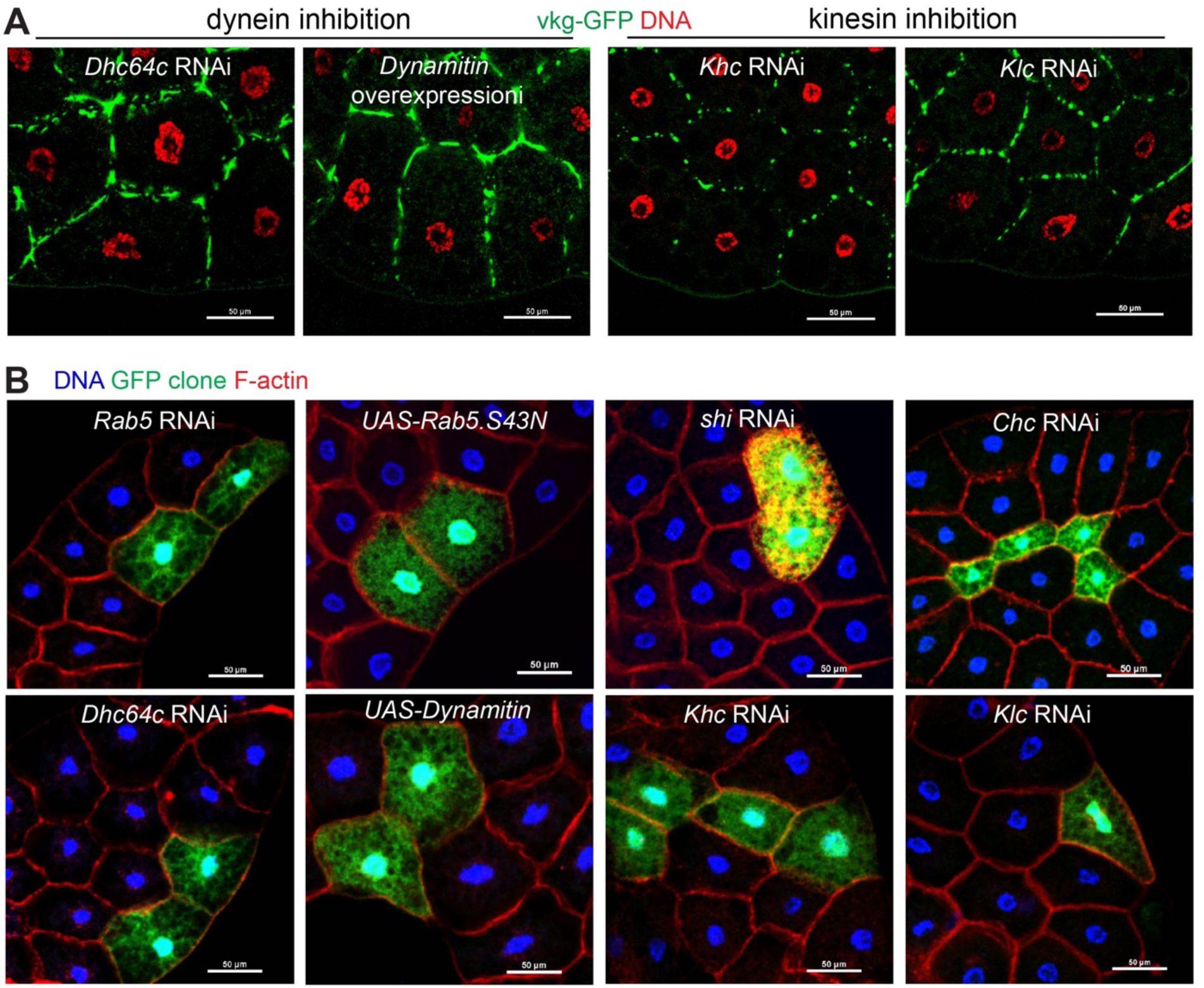
Retrograde trafficking is essential to restrict plasma membrane growth and secretion of BM proteins but does not affect nuclear positioning. (**A**) Images showing accumulation of Vkg-GFP at the the plasma membrane of fat body cells after inactivation of dynein activity with *Dhc64c* RNAi or Dynamitin overexpression, but not after inactivation of kinesin-1 by *kinesin heavy chain* (*Khc*) or *kinesin light chain* (*Klc*) RNAi. (**B**) Images showing normal nuclear positioning after inactivation of dynein or kinesin-1 motors (*Dhc64c* RNAi, UAS-dynamitin, *Khc* RNAi, *Klc* RNAi), or disruption of endocytic machinery (*Rab5* RNAi, *Rab5-S43N*, *shi* RNAi, *Clathrin heavy chain* (*Chc*) RNAi) in GFP-marked clones. Nuclear quantification is shown in Figure 8B. Note *Chc* knockdown cells are smaller, and *shi* cells have aberrant F-actin aggregation. Scale bars, 50 μm.

Taken together, we show two key functions for the fat body MTOC: nuclear positioning and retrograde endosomal trafficking. These two functions are separable because blockade of endosomal machinery or dynein motor activity caused membrane thickening and BM protein accumulation without affecting nuclear positioning (Figure 36, 37). These results demonstrate that nuclear positioning is directly controlled by the perinuclear ncMTOC, or specifically the radial MT array. However, successful collagen secretion relies on plasma membrane maintenance, a more complex process that requires not only the perinuclear ncMTOC-organized radial MT array as a track to facilitate retrograde endosomal trafficking, but also the endocytic machinery to generate plasma membrane-derived vesicles, and dynein to power their transport (Figure 38).

**Figure 38.**
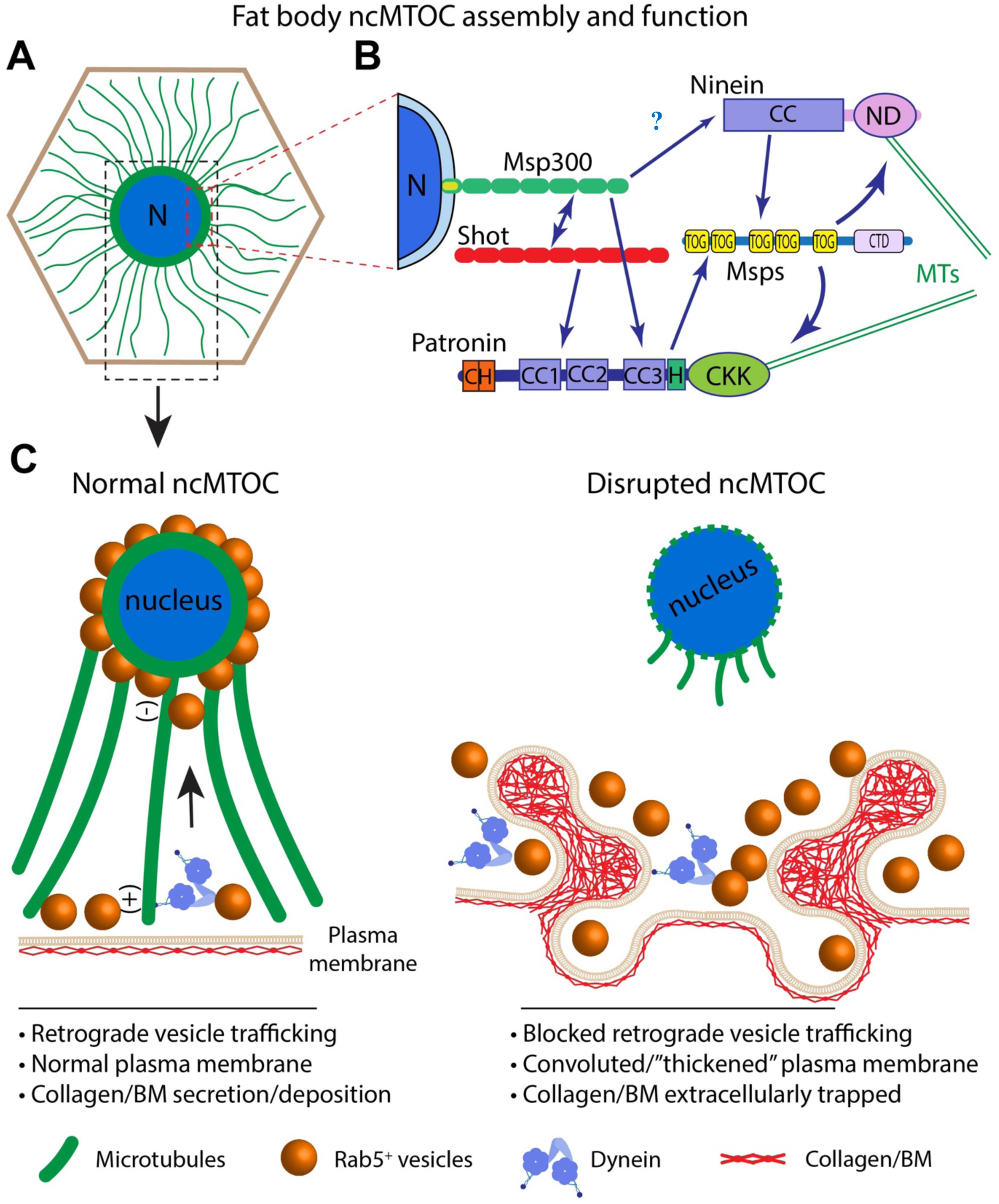
Summary and model for perinuclear non-centrosomal MTOC assembly and function in fat body. (**A**) *Drosophila* fat body cells, a differentiated cell type analogous to human adipocytes and liver, assemble a perinuclear ncMTOC with dense circumferential MTs and radial MTs. The radial MTs are polarized emanating from the nuclear surface (minus-end) towards the plasma membrane (plus-end). The perinuclear ncMTOC is distinct from the centrosome in composition. MT-regulatory proteins and many pericentriolar matrix (PCM) proteins, but not core centriolar proteins, are components of the ncMTOC (not depicted). N: nucleus. (**B**) The perinuclear ncMTOC has distinct architecture and MT assembly mechanisms. Msp300/Nesprin anchors the ncMTOC at nuclear surface, requiring and recruiting Shot, and epistatic to CAMSAP/Patronin. Patronin and Nin cooperate to recruit Msps for radial MT elongation independently of *γ*-tubulin. Domains are shown for each protein. Question marks indicate putative associations. (**C**) The perinuclear ncMTOC has two critical physiological functions; nuclear positioning and control of plasma membrane growth via retrograde trafficking. The fat body perinuclear ncMTOC controls endosomal retrograde trafficking in coordination with minus-end directed dynein to restrict plasma membrane overgrowth. Disruption of the ncMTOC or inactivation of dynein blocks retrograde membrane trafficking, leading to excessive plasma membrane growth and convoluted “thickened” plasma membranes. Consequentially, secreted BM components are trapped within the convoluted plasma membrane folds. The entrapment of BM proteins in fat body cells leads to reduced BM deposition in destination tissues, including imaginal discs and brains (not depicted).

## Discussion

The rich diversity of non-centrosomal MT arrays and subcellular ncMTOC sites has long been recognized, yet the molecular architecture of ncMTOCs and whether their makeup and assembly mechanisms are similar to the centrosome, whose structure and MT regulation has been extensively investigated, have been underexplored. Here we show that an ncMTOC assembles on the surface of the nucleus and organizes a highly stabilized radial array of MTs with their minus ends anchored at the nuclear surface in *Drosophila* fat body cells (Figure 38A). This ncMTOC is unique in its molecular architecture and the mechanisms by which MT assembly is regulated. The fat body perinuclear ncMTOC is anchored at the nucleus by Msp300, one of the two KASH-domain Nesprin proteins in *Drosophila*, and the spectraplakin Shot, which is interdependent with Msp300 for recruitment to the nuclear surface. Msp300 and/or Shot recruit the key MT regulator Patronin, which cooperates with Nin to recruit the MT polymerase Msps. *γ*-tubulin, however, is notably not involved. Based on these findings, a new paradigm for non-centrosomal MT assembly emerges (Figure 38B). Functionally, the fat body ncMTOC maintains nuclear centricity, and also serves a critical function in providing a trafficking conduit for dynein motor-based transport of endocytic vesicles, whose trafficking maintains the balance of plasma membrane growth. Maintaining proper balance of membrane growth is important to support the secretion of molecules beyond the fat body as large protein assemblies such as collagen IV and other BM components become trapped in the membrane folds resulting from plasma membrane overgrowth.

### A new paradigm for MT assembly at MTOCs

One prevailing model for generation of non-centrosomal MTs is that *γ*-tubulin dominates nucleation and also anchoring of newly formed MTs, as occurs at the centrosome. In some differentiated cell types, *γ*-tubulin complexes and other PCM proteins relocate from inactive centrosomes to non-centrosomal sites for MT nucleation or anchoring (Brodu et al., 2010; Feldman and Priess, 2012; Muroyama et al., 2016; Sanchez and Feldman, 2017). During spermatid morphogenesis, *γ*-tubulin complexes are recruited to the mitochondrial surface by a single protein, an alternative splice variant of Cnn. This converts mitochondria into an ncMTOC, epitomizing perhaps the simplest molecular platform for *γ*-TuRC–centered ncMTOC formation (Chen et al., 2017). Fat body cells lack centrosomes, and *γ*-tubulin is localized at the nuclear surface together with most of the major centrosomal PCM proteins. Surprisingly, neither *γ*-tubulin nor any of the PCM proteins are essential for the assembly of MTs at the MTOC, and mutant combinations we tested did not block MT assembly (although we did not examine every possible mutant pair combination).

Patronin/CAMSAP proteins have emerged recently as key MT regulators at ncMTOCs, though their exact functional mechanisms remain elusive (Goodwin and Vale, 2010; Wang et al., 2015b; Khanal et al., 2016; Nashchekin et al., 2016; Muroyama and Lechler, 2017; Sanchez and Feldman, 2017; Wu and Akhmanova, 2017; Martin and Akhmanova, 2018; Takeda et al., 2018; Tillery et al., 2018). Patronin protects and stablizes MT minus ends by antagonizing Kinesin-13 MT depolymerases (Goodwin and Vale, 2010; Hendershott and Vale, 2014; Atherton et al., 2017). CAMSAP2 is proposed to stabilize new MTs that are nucleated by *γ*-tubulin in neurons (Yau et al., 2014). CAMSAP2 and 3 bind directly to Katanin which appears to limit the extent of CAMSAP association with MT minus ends (Jiang et al., 2014). Here we show that Patronin controls circumferential MTs, with no significant impact on radial MTs, consistent with no significant effect on nuclear positioning in *Patronin* knockdown cells. This conflicts with a recent study reporting that Patronin is required for nuclear positioning in the fat body (Sun et al., 2019). Furthermore, we show that Patronin appears to not be antagonized by Kinesin-13 in the fat body. Moreover, organization of the MT array does not require the MT severing enzymes Kat60 or Kat-60L1, also in conflict with a report that MT severing by Kat-60L1 is required for perinuclear MT assembly in fat body cells (Sun et al., 2019). Instead, we show that Nin functions in parallel with Patronin to generate the fat body ncMTOC without the functional dependence of the *γ*-tubulin complex. In *C. elegans* epidermal cells, the Nin ortholog NOCA-1 is recruited by the *γ*-tubulin complex, and they work together and redundantly with Patronin for the establishment of a plasma cell membrane-localized ncMTOC (Wang et al., 2015b). In contrast to the Patronin redundancy with the Nin ortholog in *C. elegans*, our findings show that Patronin and Nin cooperate without the involvement of *γ*-tubulin. Whether the Patronin-Nin pair of MT regulators functions at other ncMTOCs remains to be determined.

Patronin/CAMSAPs are established stabilizers of MT minus ends which likely explains how they function to promote MT assembly at ncMTOCs, yet how Nin works is unknown as its role as a MT anchor is unclear. We show that co-depletion of Patronin and Nin significantly reduces the recruitment of Msps, a MT polymerase recently identified to function *in vivo* as a MT nucleator cooperatively with *γ*-tubulin. Since *γ*-tubulin and other known Msps cofactors are not involved, these data point to a novel paradigm for MT assembly at an MTOC where we propose that Patronin and Nin are required for assembly, anchoring and/or stabilization of MT seeds that are elongated by Msps (Fig 38A, B).

### A Nesprin-Shot complex anchors the MTOC at the nuclear surface

Msp300/Nesprin is the molecular foundation that assembles the ncMTOC on the nuclear surface of fat body cells, and we show that it is codependent on Shot for its assembly at the nuclear membrane. Msp300 and/or Shot also recruit Patronin (Fig 38B), but whether this is due to direct association(s) remains to be established. Msp300 is in the spectrin/spectraplakin protein family (Roper et al., 2002) and shares structural domain features with the spectraplakin Shot, which was shown to associate with Patronin as do spectraplakin orthologs in vertebrates (Khanal et al., 2016; Nashchekin et al., 2016; Noordstra et al., 2016). It is therefore possible that Msp300, like Shot, directly binds Patronin.

Mammalian and *Drosophila* muscle myotubes also have a perinuclear MTOC (Bugnard et al., 2005; Fant et al., 2009; Srsen et al., 2009; Espigat-Georger et al., 2016; Gimpel et al., 2017). The fat body and muscle perinuclear ncMTOCs share some MTOC components (*γ*-tubulin, Pericentrin, Nin), and the MT arrays in muscle and fat body cells are both stabilized, and resistant to cold treatment (Mao et al., 2017). Nesprin/Msp300 is a critical regulator of the perinuclear ncMTOC in both mammalian and fly muscle (Elhanany-Tamir et al., 2012; Wang et al., 2015a; Espigat-Georger et al., 2016; Gimpel et al., 2017), and in mammalian muscle Nesprin is required to recruit centrosomal components to the nuclear surface, including Akap450, *γ*-tubulin, CDK5RAP2, Pericentrin and PCM-1 (Espigat-Georger et al., 2016; Gimpel et al., 2017), and Akap450, but not CDK5RAP2, Pericentrin or PCM-1, is essential to nucleate MTs in MT regrowth assays (Gimpel et al., 2017). In *Drosophila* muscle, Msp300/Nesprin localizes to the nuclear surface and works cooperatively with Klar in the organization of the MTOC and myonuclear positioning (Elhanany-Tamir et al., 2012). In the fat body however, a *klar* null mutant has a small but insignificant effect on fat body nuclear positioning (Fig. 3a, Table 1). Furthermore, the fat body perinuclear MTOC differs from the muscle MTOC in that the muscle requires ensconsin/MAP7 (Metzger et al., 2012), EB1 (Wang et al., 2015a), CLIP-190 (Folker et al., 2012), Kinesin-1 (Metzger et al., 2012), and Dynein (Folker et al., 2012) for myonuclear positioning/nuclear morphology, but these proteins are not required for fat body MTOC organization or nuclear positioning. In addition, Nesprins can anchor MTs through associations with MT motors in other systems (Lee and Burke, 2018), but the role described here for Msp300 is distinct from those mechanisms because in the fat body Msp300 is a structural platform for assembly of the ncMTOC at the nuclear surface, seemingly independent of Kinesin-1 or Dynein motors. Morever, depletion of Nesprin-1 or SUN1/2 in mammalian muscle cells shifts the perinuclear MTOC to ectopic MTOCs associated with perinuclear Golgi complexes (Gimpel et al., 2017), an MTOC switch similar but distinctily different from *shot* depletion in fat body cells.

Msp300 is expressed widely in *Drosophila* (Technau and Roth, 2008; Xie and Fischer, 2008; Elhanany-Tamir et al., 2012), yet why Msp300/Nesprin organizes a perinuclear MTOC in a cell type-specific manner is unclear. It may involve tissue/cell-type specific isoforms of Msp300/Nesprin resulting from alternative splicing. Nesprins switch isoforms at the nuclear envelope to control nuclear positioning during muscle development (Randles et al., 2010), and a smaller isoform of Nesprin-1, Nesprin-1α, was upregulated and required for the mammalian muscle perinuclear MTOC (Gimpel et al., 2017). While a similar Msp300 isoform could account for the fat body perinuclear MTOC, alternatively, involvement of a unique co-factor, such as an isoform of Shot (which also has numerous alternative splice products), could conceivably connect Msp300 at the nuclear surface to the MT assembly machinery (Patronin, Nin, and Msps).

### The fat body ncMTOC controls nuclear positioning and plasma membrane growth

The critical function for the ncMTOC in fat body cells is the establishment of radial MTs that are necessary for endocytosis to restrict plasma membrane growth (Fig 38C). The MTOC is also critical for nuclear positioning, but whether nuclear positioning is important was not established as we were unable to separate it from MT organization and the critical role for the radial MT array in endocytic vesicle trafficking. However, it is clear that maintaining nuclear positioning and the MT array is not sufficient to control endosomal trafficking without the MT minus-end directed motor dynein, which is known to be essential for endosomal trafficking in other systems (Granger et al., 2014) as we show it is in the fat body. This was evident in the requirement for dynein for Rab5 trafficking and to restrict membrane growth. When the MTOC is disrupted, secretion of collagen and other BM proteins are impaired as a consequence of plasma membrane overgrowth due to an imbalance in membrane trafficking from blocked retrograde endosomal trafficking. This is consistent with electron microscopic analysis of fat body plasma membranes following endocytic block, where the plasma membrane becomes highly convoluted and traps secreted BM proteins and prevents them from exiting the fat body surface (Dai et al., 2017). The ncMTOC is essential to provide trafficking conduits for endocytic vesicles to restrict plasma membrane growth, and a failure to restrict membrane growth traps BM proteins in the convoluted and thickened membrane and blocks their release (Fig 38C).

In summary, the work we present here defines an essential ncMTOC in fat body cells whose regulation involves a new paradigm for non-centrosomal MT assembly. This novel ncMTOC controls nuclear positioning and plays a vital role in supporting secretion of large assemblies like collagen. The ncMTOC supports collagen secretion by balancing plasma membrane growth through retrograde dynein-controlled trafficking of plasma membrane-derived endosomal vesicles along the radial MTs.

## Methods and Materials

### *Drosophila* stocks

Information for the source, identifier, and original reference for all fly strains used, including mutants, RNAi lines, transgenic lines and drivers were listed in Supplementary Table 2. Flies were maintained on standard food and incubated at 25°C, except that RNAi-mediated knockdowns were crossed at 29°C.

### Plasmid constructs and generation of UASp-CFP-Msps *Drosophila* strain

The coding sequences for Msps were amplified by PCR from cDNA clone LP04448 (Acc# BT023496), cloned into pENTR-DTopo (ThermoFisher), and sequenced before shuttling into pPCW-attB (gift from Michael Buszczak) to generate UASp-CFP-Msps, and into pPHW-attB to generate UASp-HA-Msps. UASp-CFP-Msps plasmid was injected into embryos for targeted insertion on chromosome 3 at ZH-96E by Bestgene, Inc.

### Generation of mutant and FLP-out expression clones in fat bodies

Gain-of-function (Flp-out) clones were generated by crossing virgin females: *hsFlp; UAS-Dcr-2; Act>CD2>Gal4, UAS-GFP /TM6B* or *hsFlp; Act>CD2>Gal4, UAS-His-RFP/TM6B* to UAS-driven RNAi lines or other transgenes. FLP-mediated excision of the CD2 insert induces gene knockdown or overexpression driven by *Act-Gal4* with *UAS-Dcr-2* at 29°C in GFP-marked cells. These clones were generated in early embryos by shift to 29°C, and fat body precursor cells as evidenced by clones that typically include multiple cells due to mitotic expansion of the clone prior to differentiation.

hsFLP/Flippase Recognition Target (FRT)-mediated *Msp300* loss-of-function clones in larval fat body were induced in 0–6 h embryos by a 1-h heat shock at 37°C using an FRT-linked *Msp300^complete^* mutant allele. Mutant clones were marked by loss of GFP.

### Antibodies

Mouse anti-α-tubulin (DM1A; 1:1000 for indirect immunofluorescence staining [IF]; Cat#T9026, Sigma-Aldrich, St. Louis, MO), rat monoclonal anti-α-tubulin (YL1/2; 1:1000 for IF; Cat#MA1-80017, Thermo Fisher, Waltham, MA), FITC-conjugated α-tubulin (DM1A; 1:100 for IF; Cat#F2168, Sigma); mouse anti-β-tubulin (E7; 1:100 for IF; Developmental Studies Hybridoma Bank (DSHB), University of Iowa); mouse anti-acetylated tubulin (6-11B-1; 1:1000 for IF; Cat#6793, Sigma), mouse monoclonal anti-glutamylated tubulin (B3; 1:200 for IF; Cat#T9822, Sigma), mouse anti-*γ*-tubulin (GTU88; 1:1000 for IF; 1:10,000 for WB; Cat#T6557, Sigma), rabbit or guinea pig anti-Cnn (1:1000 for IF; 1:10,000 for WB; (Kao and Megraw, 2009)), mouse monoclonal anti-lamin C (LC28.26; 1:100 for IF; DSHB), mouse monoclonal anti-β-Gal (40-1a; 1:50 for IF; DSHB), mouse monoclonal anti-GFP (3E6; 1:1000 for IF; Cat#A-11120, Invitrogen), chicken anti-GFP (1:500 for IF; Cat#GFP-1020, Aves labs), rabbit anti-RFP (1:500 for IF; Cat#AB3216, Chemicon), anti-HA (HA-7; 1:40,000 for WB; Cat#H9658, Sigma), rabbit anti-TPI (1:10,000 for WB, Cat#sc-30145, Santa Cruz Biotechnology), rabbit anti-Msps (1:500 for IF, (Kao and Megraw, 2009)), guinea pig anti-Msp300 (1:100 for IF; (Elhanany-Tamir et al., 2012)), mouse anti-ATP5*α* (15H4C4; 1:1000 for IF; Cat#ab14748, Abcam), mouse anti-Klar (9C10; 1:100 for IF; DSHB), mouse anti-Shot mAbRod1 (1:200 for IF; DSHB), guinea pig anti-Ninein (1:100 for IF; 1: 1000 for WB (Iampietro et al., 2014), rabbit anti-Patronin (1:200 for IF; 1:1000 for WB (Goodwin and Vale, 2010)), rabbit anti-Spd2 (1:1000 for IF (Dix and Raff, 2007)), rabbit anti-ensconsin (1:5000 for staining, (Barlan et al., 2013)), rabbit anti-Grip84/GCP2 (1:500 for IF;(Oegema et al., 1999)), rabbit anti-Grip91/GCP3 (1:500 for IF;(Oegema et al., 1999)); rabbit anti-Grip128/GCP5 (1:500 for IF;(Oegema et al., 1999)); rabbit anti-centrocortin (1:1000 for IF (Kao and Megraw, 2009)); guinea pig anti-Cg25C (1:200 for IF; (Shahab et al., 2015)).

See more details for all the antibodies used in Supplementary Table 3.

### Immunostaining

Most stainings were done in fat bodies from late third instar wandering larvae, except that early stages of larvae were included in Figure 9, 10.

MTOC components staining in larval fat bodies was performed after methanol fixation as previously described with minor modification (Zheng et al., 2016). Fat bodies were dissected from late third instar wandering larvae in 1× phosphate-buffered saline (PBS). The dissected fat bodies from one larvae were transferred to 8 µl of PEM solution (100 mM PIPES, pH 6.9, 1 mM EGTA, and 2 mM MgSO_4_) on a slide. A 22 × 22 mm siliconized coverslip containing 1 μl of 18.5% formaldehyde in PEM was placed on the slide, allowing the weight of the coverslip to flatten the fat bodies for 10 s, followed by snap freezing of the slide in liquid nitrogen. The coverslip was pried off using a razor blade and the slide with attached fat bodies was immediately fixed in −20°C methanol for 10 min, followed by PBS rinse. A hydrophobic ring was drawn around the fixed fat bodies using a Super PAP Pen (Immunotech, Monrovia, CA). The fat bodies were incubated with 50 µl of antibody solution (1×PBS, 5% BSA and 0.1% saponin) in a humid chamber overnight at 4°C for primary antibody and 2 hr at room temperature for secondary antibody.

Fat body co-staining of microtubules and F-actin was performed after fixation with paraformaldehyde (PFA). Briefly, dissected fat bodies were fixed for 10 min in 50 µl of 4% PFA within a hydrophobic ring drawn on a slide. The fixed fat bodies were washed twice with PBS, followed by blocking with PBSBT (1×PBS, 0.1% Triton X-100, and 1% BSA) for 1 h and incubated with FITC-conjugated DM1A (1:200, Sigma) and CF568-conjugated Phalloidin (1:200, Cat#00044, Biotium) in PBSBT for 2 h in the hydrophobic ring.

For plasma membrane staining with CellMask Orange (C10045, Invitrogen), dissected fat bodies were directly stained with CellMask Orange (1:1000) for 5 min without inclusion of detergents. After washing three times with PBS, the stained fat bodies were then fixed with 4% PFA for 10 min and washed three times again after fixation. Mounted samples were immediately imaged.

Non-permeabilization anti-GFP staining of larval fat bodies expressing Vkg-GFP was performed similarly in 4% PFA-fixed fat bodies as above described, except that no detergent was used during the whole wash, block and antibody incubation.

### Transmission electron microscopy (TEM)

TEM analysis of the MTs at the fat body ncMTOC was performed essentially as described (Chen et al., 2015), except that fat bodies from 30 third instar larve were fixed for 72 hrs in 1 ml Karnovsky’s fixative (EM Sciences Cat#15720). Embedding, staining, sectioning and preparation of grids were performed by the Core Facility at UT Southwestern Medical Center. After three rinses with 0.1 M sodium cacodylate buffer, samples were embedded in 3% agarose and sliced into small blocks (1mm^3^), rinsed with the same buffer three times and post-fixed with 1% osmium tetroxide and 0.8 % potassium ferricyanide in 0.1 M sodium cacodylate buffer for 1.5 h at room temperature. Samples were rinsed with water and *en bloc* stained with 4% uranyl acetate in 50% ethanol for 2 h. They were then dehydrated with increasing concentration of ethanol, transitioned into propylene oxide, infiltrated with Embed-812 resin and polymerized in a 60°C oven overnight. Blocks were sectioned with a diamond knife (Diatome) on a Leica Ultracut 7 ultramicrotome (Leica Microsystems) and collected onto copper grids, post stained with 2% aqueous Uranyl acetate and lead citrate. Images were acquired on a Phillips CM120 Biotwin transmission electron microscope at the Florida State University Biological Science Imaging Resource (BSIR).

### Coimmuniprecipitation (CoIP)

Co-IP of HA-Msps and Patronin was performed with Drosophila S2 cells. S2 cells were cultured in Shields and Sangs M3 medium (Sigma cat# S3652) with 10% fetal bovine serum (Gibco cat#10437-010) at room temperature. pMT-GAL4 (DGRC cat#1042) was used to induce expression of pUASp-HA-Msps. Plasmids (pMT-GAL4 plus or minus pUASp-HA-Msps) were transfected using Lipofectamine 3000 (Thermo Fisher) in 6-well dishes. After 48 hrs of culture, expression was induced by addition of 1mM CuSO_4_ for 24 hr. Cells were harvested and lysed in 500 ul lysis buffer (50mM Tris pH 7.5, 150mM NaCL, 1mM EDTA, 05% NP-40) for 10 min on ice. Immunoprecipitation was performed using anti-HA-Agarose (Sigma cat#2095) following the manufacturer’s recommended protocol. For western blots, input lanes represent 2.1% of the total lysates and IP lanes represent 62.5% of the total immunoprecipitate. Mouse anti-HA (HA-7, Sigma cat#H9658, 1:40,000) and rabbit anti-Patronin (1:500)(Goodwin and Vale, 2010) were used to detect HA-Msps and endogenous Patronin, respectively. Western blotting was performed as described below.

### Western blot

Fat bodies or brains dissected from five wandering larvae were lysed respectively in 50 µL or 10 µL of 2×SDS-PAGE buffer (100 mM Tris-HCl, pH 6.8, 4% SDS, 0.02% bromophenol blue, 20% glycerol, 5% β-mercaptoethanol). 10 whole larvae at wandering stage were lysed with motorized pestle in 200 µL of 2×SDS-PAGE loading buffer. After boiling for 5 min, larval lysates were centrifuged at 13,200g for 5 min to clear pellets. 10 µL of larval lysates were loaded for SDS-PAGE gel electrophoresis, followed by semi-dry transfer to nitrocellulose membrane. The membranes were blocked with 5% nonfat milk in 1×TBS for 1 h at room temperature and then probed with primary antibodies diluted in 1×TBS containing 0.1% Tween (1×TBST) overnight at 4°C. After washing with 1×TBST three times, the membranes were incubated with secondary antibodies conjugated with IRDye-800CW or IRDye-680LT (1:20,000) for 1 h at room temperature. Blots were scanned on an Odyssey Infrared Imaging system (LI-COR Bioscience, Lincoln, NE). Signals were quantified using Li-Cor Image Studio software. Images were processed in Adobe Photoshop CS4 and presented in monochrome.

### Microtubule regrowth assay

Wild-type fat bodies were dissected and placed in 15 µl of 10 mg/ml fibrinogen (EMD#341573) in D-PBS (Dulbecco’s PBS, Invitrogen) on a clean slide within a hydrophobic ring drawn with a super PAP pen. 2 µl of thrombin solution (Sigma T9549-50UN; 100 U/ml in D-PBS) was pipetted to the fibrinogen solution to induce a fibrin clot. The fat bodies were treated with 30 µM vinblastine in Shields and Sangs M3 medium (Sigma # S3652) for 1-2 h to induce MT disassembly at room temperature. The vinblastine was washed out in ice-cold D-PBS (0-4°C, or at a constant 4°C) for 1 h, followed by a time-course recovery at 25°C. The fat bodies were fixed at the respective time points in −20°C methanol for 10 min and stained with anti-α-tubulin and DAPI to label MTs and nuclei, respectively. Microtubule regrowth in the *γ-Tub23C* RNAi clone experiment (Figure 12) was fixed with 4% PFA to enable staining with Alexa568-phalloidin.

### Image acquisition

Fixed fat body samples were imaged using a Nikon A1 laser scanning confocal microscope (Nikon, Japan) using a 60×/1.49 NA oil immersion objective. All confocal images were captured with a spacing of 0.25 μm or 0.5 μm between *z*-sections using Nikon NIS-Elements AR software (version 4.6) and are presented as maximum intensity projections of *z* stacks. Images of larval wing discs and brains were acquired by stitching together images acquired with 25% overlap using in Elements using a 60×/1.49NA oil immersion objective. Gamma-correction was applied to images.

Live imaging of intact fat body in whole larvae expressing vkg-GFP and myr-RFP was performed by a Macro Zoom microscope (MVX10, Olympus, Japan) equipped with DP72 camera. DIC view of whole larvae was imaged at 32-fold magnification. Fluorescent imaging of Vkg-GFP and myr-RFP expressed in larval fat bodies at single-cell resolution was performed at 130-fold magnification. Representative still frames were shown in Figure 29B.

Live imaging of nuclear positioning/movement in intact fat body expressing myr-RFP and Histone-GFP was performed on whole live third instar wandering stage larvae restrained between a 22×22 mm and a 24×40 mm 1.5 coverslip using clear ¼-inch wide office tape. Images were captured every 3.0 s using 488 nm and 568 nm laser excitation with a 20×/0.75 NA Plan Apo objective on a Nikon A1 confocal microscope with the pinhole open to 8.0. The time-lapse image sequence was converted to .avi video at 20 fps.

### Quantification analysis

Nuclear positioning, or centricity analysis, in fat bodies was measured as the distance between the cell’s geometric center and nucleus’ geometric center using Nikon NIS-Elements AR software (version 4.6). Cell and nuclear geometric centers were measured using the “centroid” tool after auto- or manual thresholding of cell boundary (cortical F-actin by Phalloidin staining) and nucleus (Hoechst staining). The distance between cell and nuclear geometric centers was measured using the length tool in NIS-Elements.

Fluorescent intensity of circumferential and radial MTs, Msps or Patronin in GFP or His-RFP-marked clones was analyzed in Image J. Integrated density and area for circumferential or radial MTs was measured in 8-bit inverted monochrome images. Mean fluorescence intensity was measured as integrated density per unit area using the measure tool in ImageJ on 8-bit monochrome inverted and thresholded images. For each RNAi knockdown, analysis was performed in control and GFP-marked clones for side-by-side comparison.

Vkg-GFP fluorescent intensity in larval wing discs was quantified in Image J. Mean fluorescent intensity of GFP auto-fluorescence was measured after GFP signal thresholding in 8-bit inverted images.

### Statistics

Data are presented as mean ± s.e.m. Dissected fat bodies from at least five larvae were included for nuclear centricity analysis. At least four fat body clones were included for fluorescent intensity analysis. At least four wing discs were analyzed for Vkg-GFP fluorescent intensity analysis. The statistical significance was determined by two-tailed Student’s *t*-test using Prism 7 (Graphpad, San Diego, CA). Differences were considered statistically significant when *p*<0.05. * denotes *p*<0.05, ** for *p*<0.01, *** for *p*<0.001 and **** for *p*<0.0001 as shown in figures and figure legends.

## Supporting information

Supplemental Movie 1-wild type

Supplemental Movie 2-Msp300RNAi

## Acknowledgements

We thank Megraw lab members for numerous discussions and critiques of the manuscript, and three anonymous reviewers for constructive recommendations that improved the work. We thank Melissa Rolls, Wu-Min Deng, Jordan Raff, Ron Vale, Talila Volk, Nina Sherwood, Renata Basto, Nasser Rusan, Eric Lécuyer, Tomer Avidor-Reiss, Maurice Ringuette, Dorothy Lerit, Steve Rogers, James Wakefield, Hiroyuki Nakanishi, and Volodya Gelfand for antibodies and Drosophila stocks. We thank the Bloomington Drosophila Stock Center, Vienna Drosophila Resource Center, Kyoto Stock Center for Drosophila stocks and the Developmental Studies Hybridoma Bank, University of Iowa, for antibodies. We are grateful for funding from NIH grants R15GM119078 and R15HD099648 (T.L.M).

